# A novel mutation (E83Q) unlocks the pathogenicity of human alpha-synuclein fibrils and recapitulates its pathological diversity

**DOI:** 10.1101/2021.11.21.469421

**Authors:** Senthil T. Kumar, Anne-Laure Mahul-Mellier, Ramanath Narayana Hegde, Rani Moons, Pedro Magalhães, Alain Ibáñez de Opakua, Gwladys Rivière, Iman Rostami, Sonia Donzelli, Markus Zweckstetter, Frank Sobott, Hilal A. Lashuel

## Abstract

A novel mutation (E83Q), the first in the NAC domain of alpha-synuclein (aSyn), was recently identified in a patient with dementia with Lewy bodies. We investigated the effects of this mutation on the aggregation of aSyn monomers and the structure, morphology, dynamic, and seeding activity of the aSyn fibrils in neurons. We found that it dramatically accelerates aSyn fibrillization and results in the formation of fibrils with distinct structural and dynamic properties. In cells, this mutation is associated with higher levels of aSyn, accumulation of pS129, and increased toxicity. In a neuronal seeding model of Lewy bodies (LB) formation, the E83Q mutation significantly enhances the internalization of fibrils into neurons, induce higher seeding activity and results in the formation of diverse aSyn pathologies, including the formation of LB-like inclusions that recapitulate the immunohistochemical and morphological features of brainstem LBs observed in PD patient brains.

**Teaser:** A novel mutation (E83Q) exacerbates alpha-synuclein aggregation and toxicity and reproduces PD pathological diversity.

## Introduction

The accumulation of fibrillar and aggregated forms of the presynaptic protein alpha-synuclein (aSyn) within Lewy bodies (LBs) and Lewy neurites (LNs) is a characteristic hallmark of many synucleinopathies, including Parkinson’s disease (PD), multiple system atrophy (MSA), and dementia with Lewy bodies (DLB). Several missense point mutations in the *SNCA* gene, which encodes for aSyn, have been linked to familial forms of PD: A53T (*1–3*), A30P (*4*), E46K (*5*), H50Q (*6, 7*), A53E (*8*), A53V (*9*), and A30G (*10*). Additionally, the G51D mutation has been linked to a form of synucleinopathy with shared characteristics between PD and MSA (*7, 11*). Furthermore, *SNCA* gene multiplications were shown to be sufficient to cause PD and DLB (*12–17*). Although aSyn mutation carriers are rare, studies on the biochemical, cellular, aggregative, and toxic properties of these mutants have provided valuable insights into the mechanisms of aSyn aggregation and PD pathology. These studies have also suggested that the various mutations may act via distinct mechanisms.

Recently, Kapasi et al. (2020) reported the discovery of a novel *SNCA* mutation encoding for a glutamic acid-to-glutamine (E83Q) substitution in a patient with DLB and atypical frontotemporal lobar degeneration (*18*). Post-mortem neuroimaging and histology of the patient’s brain revealed a widespread LB and LN pathology, with severe atrophy of the frontotemporal lobes that correlates with cognitive impairment. As reported in some MSA (*19*), the brain of the E83Q mutation carrier showed higher LB pathology in the hippocampal neurons than the substantia nigra where much less pathology was detected. In addition, severe LB pathology was also detected in the cortex and other brain regions, including the thalamus and the basal ganglia. There was no evidence of Tau or TDP-43 pathology in the brain of this patient. Of note, the autopsy report of the patient’s father included a diagnosis of Pick’s disease. The presence of the *SNCA* E83Q mutation in the father was later confirmed (*18*).

Unlike all previously reported synucleinopathy-related mutations, which invariably occur in the N-terminal region spanning residues 30–53, and with most clustering between amino acids 46–53 (Fig. 1A), the E83Q mutation is within the non-amyloid component (NAC) domain (residues 61–95), which plays a critical role in catalyzing aSyn oligomerization and fibril formation (*20, 21*). Furthermore, the substitution of glutamate by glutamine at residue 83 results in the removal of a negative charge from the highly hydrophobic NAC domain, which contains only three charged residues (two glutamic acid and one lysine). These observations, combined with the fact that this mutation is associated with DLB instead of PD, suggest that it may influence the structure, aggregation, and pathogenicity of aSyn *via* mechanisms distinct from those of other mutations (*18*) and, thus, could offer valuable insights into the molecular mechanisms of aSyn pathology and neurodegeneration in synucleinopathies such as DLB.

**Fig. 1.**
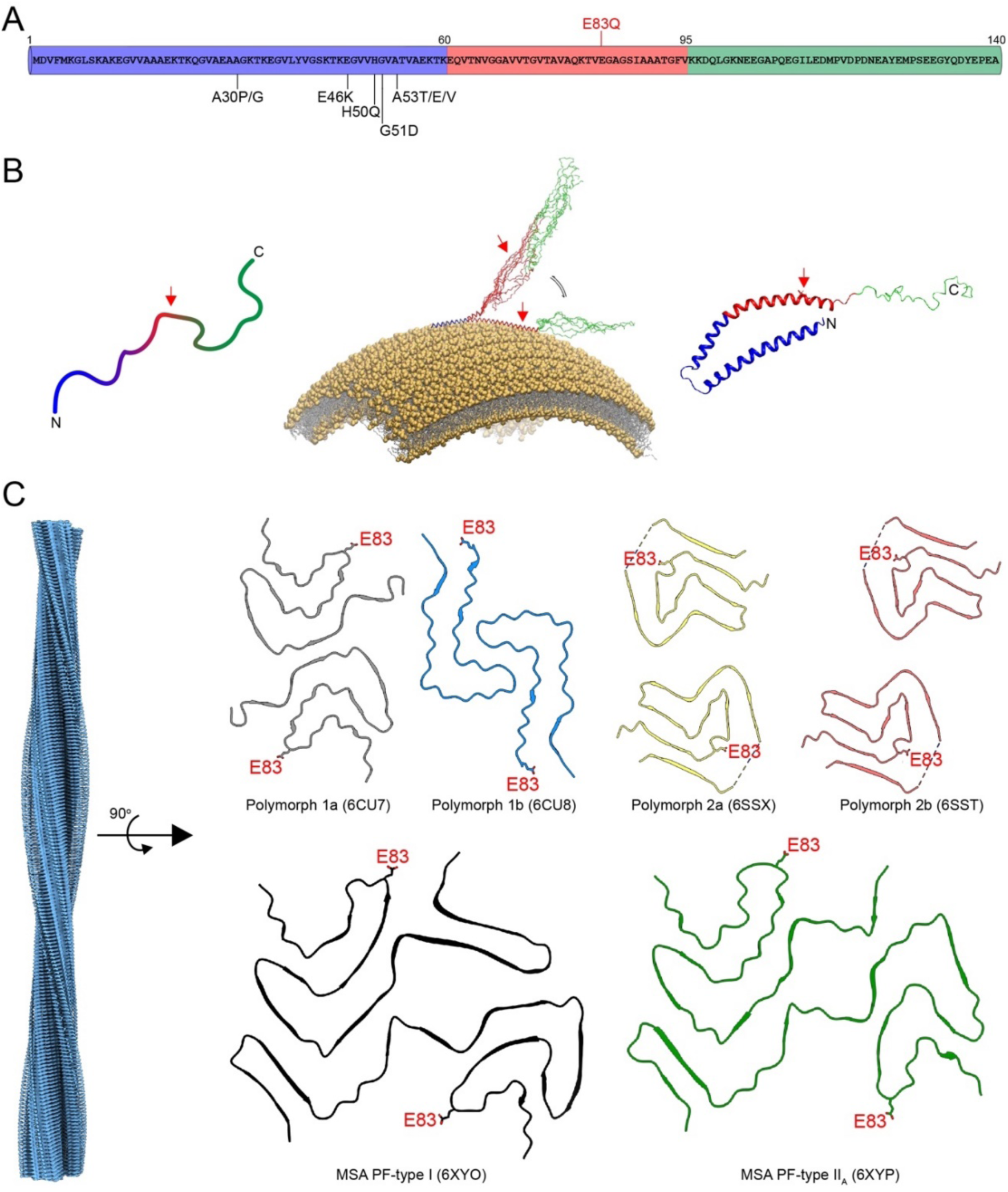
The localization of the E83Q mutation on the different forms of aSyn. **(A)** Schematic representation of the primary structure of human WT aSyn showing the three regions [N-terminal (blue), NAC (red), and C-terminal (green) region] and locations of known familial mutations. **(B)** Illustration of a disordered conformation of monomeric aSyn (left) and its α-helical conformations when membrane-bound (center and right). The schematic illustration of the structure of aSyn bound to membrane is adapted from Runfola et al. 2020 (*37*). **(C)** Location of E83 on the cryo-EM structures of *in vitro* prepared full-length aSyn fibril polymorphs 1a (PDB-6CU7), 1b (PDB-6CU8), 2a (PDB-6SSX), and 2b (PDB-6SST) and of human MSA brain-derived aSyn fibrils Type-I (PDB-6XYO) and Type-IIA (PDB-6XYP). The E83 residue lies in the loop region on the outer proximities of protofilaments, except the polymorph 2a and 2b.

Herein, biochemical and biophysical approaches, including nano-electrospray ionization mass spectrometry (nESI-MS), electron microscopy, and solid-state NMR (ssNMR) spectroscopy, were combined with cellular models to determine the effect of the E83Q mutation on: 1) the conformation and membrane binding properties of monomeric aSyn; 2) the aggregation kinetics of aSyn, as well as the morphology and structural properties of aSyn fibrils *in vitro*; 3) the subcellular localization, aggregation, and inclusion formation of aSyn in mammalian cells lines; and 4) the seeding activity and formation of *de novo* fibrils and LB-like inclusions in a neuronal seeding models of synucleinopathies (*22*).

Our *in vitro* studies demonstrated that the E83Q mutation significantly accelerated aSyn fibrillization. Fibrils generated from E83Q aSyn monomers exhibited distinct structural properties and stability compared to those formed by the wild-type (WT) protein. Overexpression of E83Q, but not WT aSyn, in mammalian cells (HEK293 or HeLa) resulted in a gain of toxic functions that are not dependent on aSyn fibrillization, but that appears to be associated with increased oligomerization. Finally, in a neuronal seeding model of LB formation (*22–24*), the E83Q mutation dramatically increased the seeding activity of human preformed fibrils (PFFs) and promoted the formation of LB-like inclusions with diverse morphological features, resembling the diversity of aSyn pathology in PD brains (*25–29*). Unlike mouse PFFs, which induce the formation of diffuse LB-like inclusions, the E83Q PFFs induced the formation of LB-like inclusions with a ring-like organization that recapitulates the immunohistochemical, structural, and morphological features of *bona fide* brainstem LBs observed in brains affected by late-stage PD (*25, 26, 30-32*).

## Results

### The E83Q mutation disrupts transient, long-range interactions of the aSyn monomer ensembles

To investigate the effect of the E83Q mutation on the conformation of aSyn monomers, we compared the circular dichroism (CD) spectra of purified WT and E83Q aSyn from *E. coli* (Fig. S1). WT and E83Q aSyn showed identical CD spectra, with a minimum at ∼198 nm, consistent with a predominantly disordered conformation for both proteins (Fig. 2A, solid lines).

**Fig. 2.**
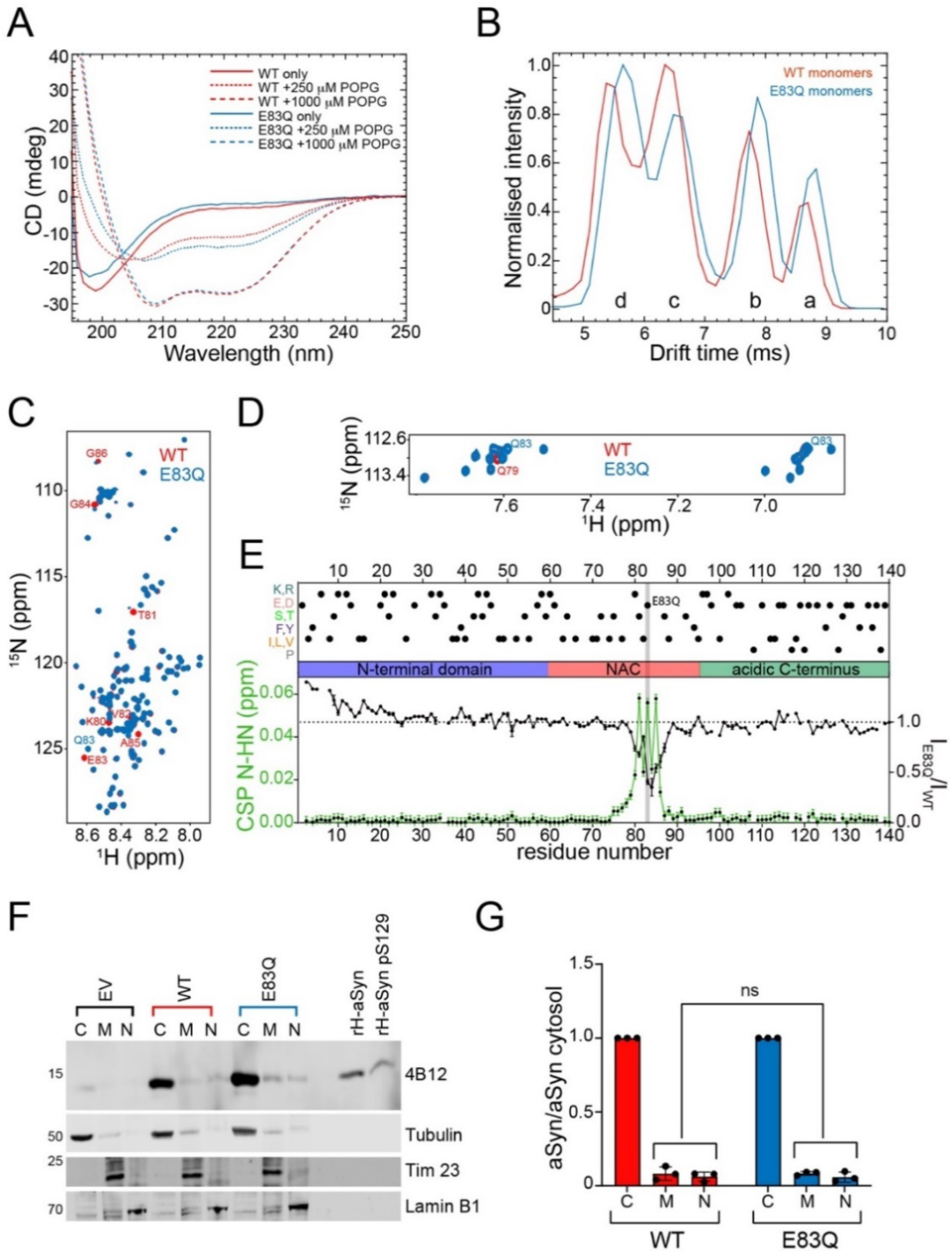
E83Q mutation affects the conformational ensembles of free aSyn monomers but not its helical conformation when bound to lipid vesicles. **(A)** CD spectra of monomeric WT and E83Q aSyn incubated with or without POPG vesicles. **(B)** Drift time plot comparison of 8+ charge state in WT (red) and E83Q mutant (blue) aSyn. Both the WT and the E83Q aSyn mutant display four unique conformational populations (labeled “a” to “d” from the most extended to the most compacted, respectively). **(C)** Backbone amide region of the ^1^H/^15^N-HMQC of WT (red) and E83Q mutant (blue) aSyn. The most perturbed residues are labelled. **(D)** Asparagine and glutamine side chain regions of the ^1^H/^15^N-HMQC of WT (red) and E83Q mutant (blue) aSyn. The new peaks of Q83 and the perturbed Q79 are labelled. **(E)** N-HN chemical shift perturbations (green) and intensity ratios (black) between WT and E83Q aSyn based on the spectrum in (C). Above the distinct domains of aSyn (N-terminal lipid-binding α-helix, NAC, and the acidic C-terminus), the most representative residues are shown in dots. The mutated residue is highlighted in grey. **(F)** Western blotting of aSyn from cytosolic (C), membranous (M), and nuclear (N) fractions of HEK293 cells expressing WT or E83Q aSyn or empty vector (EV) for 48 h (representative of 3 separate experiments). aSyn protein levels were detected with the 4B12 antibody (epitope 103-108). The purity of each subcellular fraction was determined by analyzing specific cytosolic (tubulin), nuclear (Lamin B1), and membranous (Tim 23) markers. **(G)** A plot showing the fold change of total aSyn compared aSyn levels in the cytosol (C), membrane (M), and nucleus (N), quantified using densitometry from E. Changes in the protein distribution level of WT and E83Q aSyn were non-significant (NS) for all fractions.

In solution, aSyn exists in an array of dynamic conformations (Fig. 1B) (*33, 34*). To determine whether the E83Q mutation alters the distribution of aSyn monomer conformations, we performed native nESI-MS combined with ion mobility. Ion mobility reports on the rotationally averaged shape and size (i.e., compactness) of ionized protein, providing conformational fingerprints per charge state. Data for each m/z peak was obtained as previously described (*35, 36*) and visualized in drift time plots. In Fig. 2B, drift time plots of the 8+ charge state of WT and E83Q are overlaid to compare the conformation distributions.

We observed four distinct conformational populations at this charge state for both WT and E83Q monomers. The conformations for the WT monomers were identical to those previously found (*35*). The E83Q mutant showed a slight shift towards the two more extended conformations (a and b), and within the distributions of compact states (c and d) there was a shift towards the most compact state (d), which was reflected in the disparities of the peak intensities (Fig. 2B). To investigate whether monomer conformations are stabilized or destabilized by the mutation, collisional activation experiments were performed for 7+, 8+, and 11+ charge states. The results are discussed in detail in the supplementary information (Fig. S2) and show subtle perturbations in conformational stability, depending on the charge state. Some individual conformational sub-states are slightly stabilized, while others appear with somewhat reduced stability

To further delineate the effect of the E83Q mutation on the conformational ensemble of the aSyn monomer in solution, we compared the nuclear magnetic resonance (NMR) spectra of 15N-labeled WT and E83Q aSyn proteins. The differences observed in the chemical shifts (chemical shift perturbations, CSPs) of the 1H/15N-HMQC and 1H/13C HSQC spectra of the mutant were present locally around the mutation (Fig. 2C–E and Fig. S3A–C). The CSPs were small even for the mutated residue, as expected for the simple change from an -OH to -NH2 group upon mutation of a glutamic acid to a glutamine. However, the perturbation was not limited to the immediate vicinity of the site of mutation (Fig. 2D). In the glutamine and asparagine sidechain region (Fig. 2D), we observed, besides the new peak of the new residue Q83, a CSP in one of the protons of residue Q79. This might be attributed to the presence of a polar interaction between Q79 and residue 83 and can also provide a rationale for the more extended CSP for residues N-terminal to the site of mutation. Notably, the peak intensity analysis of the same spectra revealed a decrease in intensity in the region of the mutation and an increase in NMR signal intensity for residues 1-23 at the N-terminus of aSyn (Fig. 2E). Because the 23 N-terminal residues of aSyn are predominantly positively charged, the changes in NMR signal intensities in this region might arise from changes in transient long-range electrostatic interactions of the aSyn ensembles due to the removal of the negative charge at position 83. Thus, both nESI-MS and NMR spectroscopy indicate that the E83Q mutation perturbs the conformational ensemble of aSyn in solution.

### The E83Q mutation does not significantly disrupt aSyn interactions with membranes

Since E83Q mutation lies in the distal region of aSyn that binds to membranes (Fig. 1B), we investigated if the E83Q mutation affects aSyn’s binding properties to membranes. It is known that WT aSyn adopts α-helical conformations upon binding to lipid membranes (*34, 37*). Therefore, we used CD spectroscopy to compare the propensity of WT and E83Q aSyn to bind to liposomes of distinct lipid composition [POPG and DOPE:DOPS:DOPC (5:3:2)]. As seen in Fig. 2A and S3D–F, we observed an identical secondary structural transition from unfolded to α-helical conformations for both proteins, irrespective of lipid vesicle composition or ratios of lipids to aSyn. These observations indicate that the E83Q mutation does not affect the conformation of membrane-bound aSyn or the binding of aSyn to membranes.

To validate these findings physiologically, we assessed aSyn-membrane binding properties in cells. We transiently overexpressed E83Q or WT aSyn in HEK293 cells and assessed the aSyn distributions in the cytosolic, membranous, and nuclear subcellular fractions. As shown in Fig. 2F-G, both E83Q and WT aSyn are predominantly localized in the cytosolic compartment and accumulate minimally in the membrane or nuclear fractions. (The epitopes of the antibodies used in this study can be found in Fig. S4). Altogether, our data indicate that although the E83Q mutation is located in the membrane-binding domain, it does not strongly alter its binding to lipid vesicles or intracellular membranes.

### The E83Q mutation accelerates the aggregation kinetics of aSyn *in vitro*

To investigate the effect of the E83Q mutation on aSyn aggregation, we compared the fibrillization kinetics of WT and E83Q aSyn at five initial protein concentrations ranging from 10 to 50 μM. Irrespective of the initial concentration, the E83Q mutant exhibited faster aggregation kinetics relative to the WT protein (Fig. 3A-B). Analysis of the lag phase from the kinetic curves revealed that the E83Q mutant exhibited ∼10-fold faster aggregation kinetics at all concentrations (Fig. 3C). The WT counterpart exhibited a gradual decrease in the lag phase, from ∼23 h at 10 μM to ∼11 h at 50 μM initial aSyn concentration, whereas the E83Q mutant at these concentrations showed lag times of ∼2.9 h and ∼0.9 h, respectively (Fig. 3C). To further validate the faster aggregation of E83Q relative to WT aSyn, the aggregation of the proteins was monitored in the absence of ThT using the sedimentation assay; namely, by quantifying the remaining soluble aSyn species as a function of time (Fig. 3D-E). Similar to the data from the ThT aggregation kinetics (Fig. 3A-B), E83Q showed significantly faster aggregation propensities than WT aSyn at all concentrations (Fig. 3D-E).

**Fig. 3.**
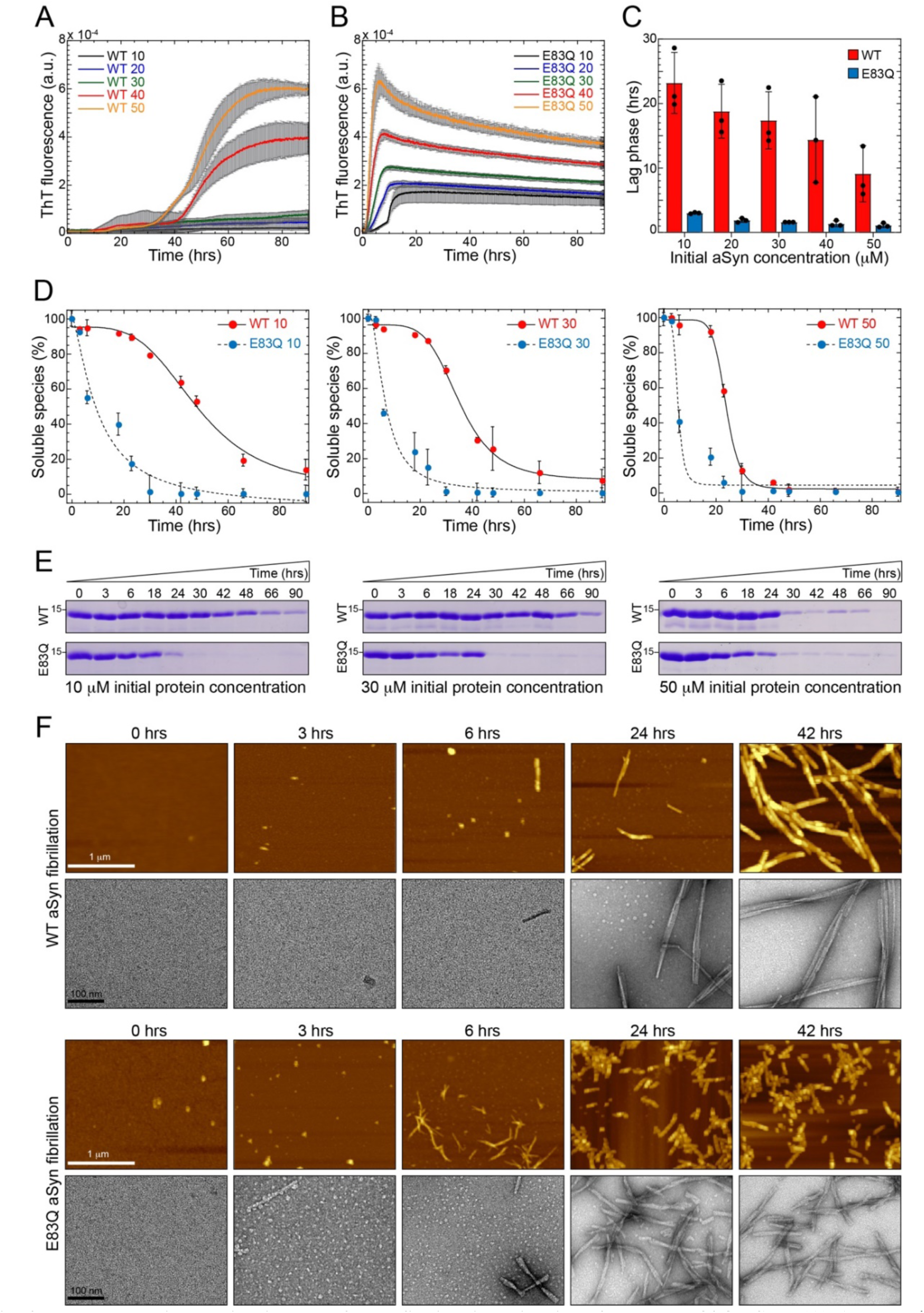
Increased oligomerization and faster fibrillation kinetics of mutant E83Q aSyn. **(A)** Aggregation kinetics of WT and **(B)** E83Q aSyn at different initial concentrations (10, 20, 30, 40, and 50 μM) monitored by ThT fluorescence. **(C)** Bar graph of lag phase extracted from the ThT aggregation kinetics shown by WT (A) and E83Q (B) (mean ± SEM, n=3). **(D)** Solubility assay of the WT and E83Q aSyn aggregations at different time-points at varying initial concentrations of 10, 30, and 50 μM. **(E)** SDS-PAGE analysis of the same samples from initial concentration. Scale bars = 1 μm for all the AFM images and 100 nm for all the TEM images. **(F)** AFM and TEM images of time-dependent WT and E83Q aSyn aggregation from 50 μM.

To determine whether the E83Q mutation influences early oligomerization events, we monitored changes in the aggregation state of E83Q and WT aSyn as a function of time by atomic force microscopy (AFM) and transmission electron microscopy (TEM) (Fig. 3F). No aggregate/oligomeric structures were observed immediately after the resuspension of either WT or E83Q monomeric proteins (0 h). At later time-points (3 h and 6 h), the E83Q, but not WT aSyn, showed a significant accumulation of oligomers. At ∼24 h of aggregation, the WT aSyn formed a mixed population of oligomeric and fibrillar structures, whereas the E83Q mutant formed predominantly shorter fibrillar structures. After ∼42 h, WT aSyn was observed to form mainly straight and long fibrillar structures, whereas its E83Q-mutated counterpart formed a preponderance of short fibrillar structures that were significantly shorter (by an average of 180 ± 97 nm) than those of WT aSyn (Fig. 3F).These findings are consistent with the results from the ThT and sedimentation assay and demonstrate that the E83Q mutation accelerates aSyn oligomerization and fibril formation and alters the size distribution and dimensions of the fibrillar structures.

### Fibrils of E83Q mutant show distinct morphology, stability, and structural features

Given that E83 occurs within the core of aSyn fibrils, we speculated that its mutation to glutamine might influence the structural properties or dynamics of aSyn fibrils. To test this hypothesis, AFM and TEM were used to quantify and compare the morphological properties of WT and E83Q aSyn fibrils. As shown in Fig. 4, both WT and E83Q fibrils were polymorphic and had single-filament (twisted) and multi-filament (stacked) morphologies. The WT fibrils consistently exceeded 1 μm in length (Figs. 4A-F and S5A), whereas E83Q fibrils were much shorter, with an average length of 180 ± 97 nm (Figs. 4A-F). These observations are highly reproducible using different batches of E83Q aSyn proteins (Fig. S5B-F). Furthermore, WT aSyn showed a broader height distribution, with an average height of 8.9 ± 2.1 nm, whereas E83Q fibrils had an average height of 10.9 ± 2.5 nm. The E83Q and WT aSyn formed fibrils of diverse yet overlapping structures (Fig. 4D-E). However, an in-depth analysis of the fibril widths of the different polymorphs revealed that the population of twisted fibrils displayed the highest variation in width between WT and E83Q aSyn (WT: 18.0 ± 1.9 nm; E83Q: 13.3 ± 1.8 nm). The other two morphologies exhibited nearly identical widths (single: ∼8 nm; stacked: ∼17 nm) between WT and E83Q aSyn fibrils (Fig. 4G).

**Fig. 4.**
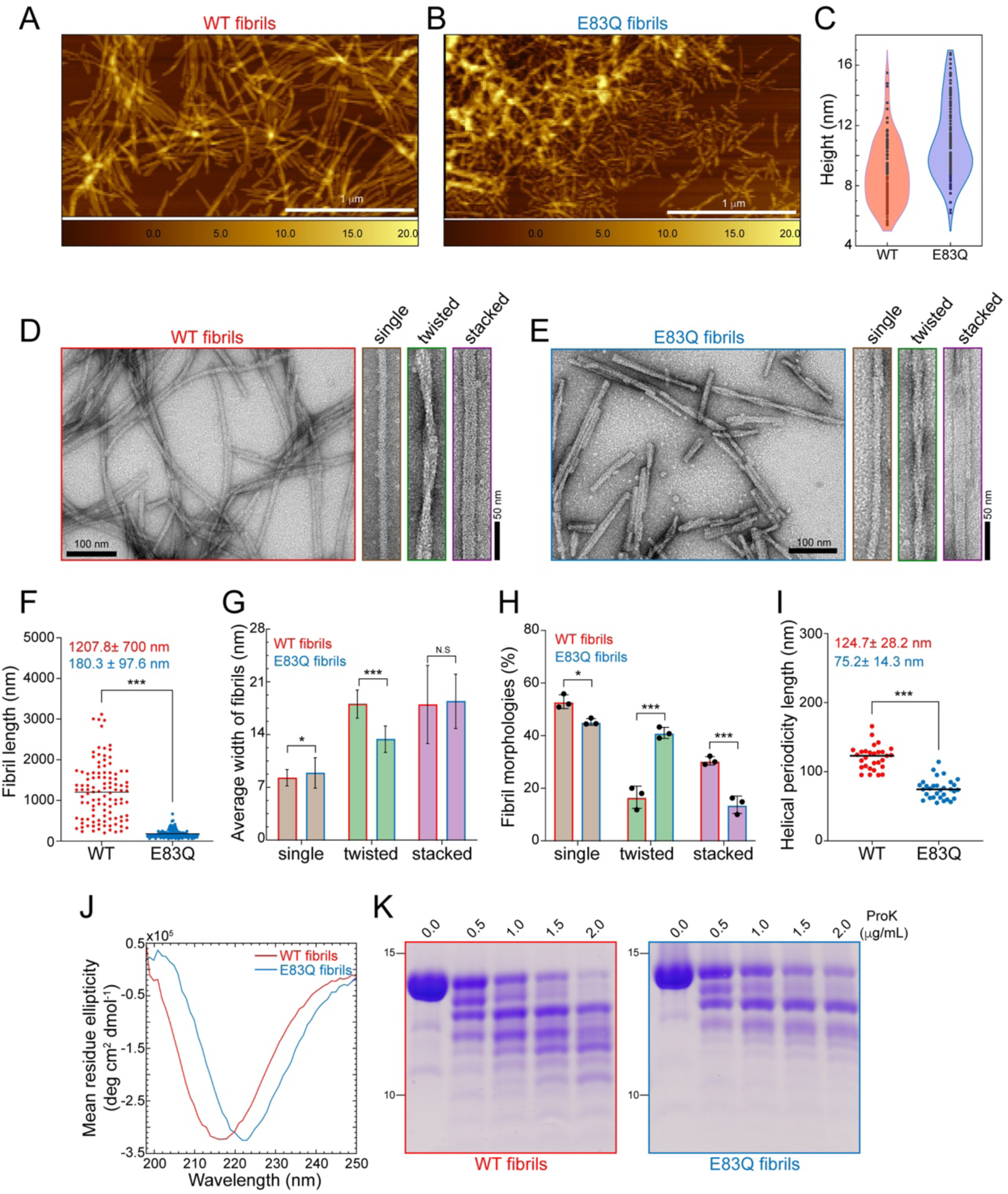
Distinct morphological and structural properties of E83Q aSyn fibrils comparing to WT fibrils. **(A, B)** AFM images of WT (A) and E83Q (B) aSyn fibrils. **(C)** Violin plot showing the AFM-based height analysis estimated from A and B, with an average of 8.9 ± 2.1 nm for WT and 10.9 ± 2.5 nm for E83Q fibrils. **(D, E)** TEM images of WT (D) and E83Q (E) aSyn fibrils. TEM montages in both D and E show the single, twisted, and stacked morphologies. **(F)** Dot plot showing the length distribution of WT and E83Q aSyn fibrils estimated from the individual fibrils (n= 108 for WT and 191 for E83Q fibrils) from various TEM images. Unpaired T-test statistical method was used for the comparison of WT vs E83Q fibrils. p<0.0005=***. **(G)** Bar graph of TEM-based analysis of average width of different morphologies of WT and E83Q fibrils. WT fibrils (single: 8.2 ± 1.0 nm (n=87); twisted: 18.0 ± 1.9 nm (n=30); stacked: 17.9 ± 5.2 nm (n=47)) and E83Q fibrils (single: 8.8 ± 2.0 nm (n=79); twisted: 13.3 ± 1.8 nm (n=68); stacked: 18.4 ± 3.6 nm (n=29)). Statistics are carried out using T-test (in R, function t.test). N.S (non-significant), p<0.05=*, p<0.0005=***. **(H)** Bar graph displays the distribution of different fibril morphologies between WT and E83Q. The graph represents the mean ± SD of three independent experiments. 2-way ANOVA shows a significant statistical interaction between the WT vs E83Q mutation and their different morphological distributions. Multiple comparison analysis with Sidak’s correction. *P < 0.05, **P < 0.005, ***P < 0.001. **(I)** Dot plot showing the helical periodicity length of twisted fibrils of WT and E83Q aSyn (n= 30 for WT and 31 for E83Q fibrils). Unpaired T-test statistical method was used for the comparison of WT vs E83Q fibrils. p<0.0005=***. **(J)** CD spectra of WT and E83Q aSyn fibrils. **(K)** SDS-PAGE analysis of ProK digestion of WT (left) and E83Q (right) aSyn fibrils.

Interestingly, while analyzing the relative frequencies of different morphologies, we observed different distributions of each of the three fibril morphologies between the two proteins (Fig. 4H). In the case of the E83Q mutant, the single-fibril morphologies predominated (∼45%), followed by the twisted (∼39%) and stacked fibril morphologies (∼16%). Although the single-fibril morphologies also predominated (∼53%) for the WT protein, the second most common morphology (∼29%) was stacked fibrils, followed by twisted fibrils (∼18%). Analysis of the helical periodicity of the twisted fibril population showed significant differences between the two proteins. The E83Q fibrils showed an average periodicity length of 75.2 ± 14.3 nm, compared to 124.7 ± 28.2 nm for the WT fibrils (Fig. 4I).

Next, we analyzed the secondary structure of fibrils using CD spectroscopy. The CD spectra of WT and E83Q aSyn fibrils showed the single peak minimum for both fibrils, suggesting the enrichment of β-sheet structures. However, a major shift of the CD minimum at 222 nm was observed for E83Q fibrils and at 217 nm for WT fibrils. This suggests the existence of pronounced differences in the arrangement/packing of *β*-sheet structures between aSyn molecules in the fibrils (Fig. 4J). Together, these findings demonstrate that the E83Q mutation significantly alters the distribution and the structural and morphological properties of aSyn fibril conformations.

To further validate our findings, we compared the proteinase K (ProK) digestion profile of WT and E83Q fibrils. To ensure the absence of cross-contamination of oligomeric or monomeric aSyn, we centrifuged the aSyn fibril samples, removed the supernatant, and resuspended the pellet containing the fibrillar structures in PBS (*38*). After a 30-min incubation of the same concentration of WT and E83Q aSyn fibrils with the increasing concentrations of ProK, the reaction mixtures were visualized by SDS-PAGE. The results revealed major differences in the stability and proteolysis pattern between WT and E83Q aSyn fibrils (Fig. 4K). While the protein band at ∼15 kDa from WT aSyn degraded almost completely at 2 μg/mL of ProK, the same band from the E83Q mutant displayed a stronger resistance to ProK proteolysis, suggesting greater stability of E83Q aSyn fibrils relative to WT aSyn. Although the band at ∼ 12 kDa was observed in both WT and E83Q samples, additional bands appeared at high concentrations of ProK in the WT, but not in the E83Q, fibril sample. At the highest concentration of ProK (2 μg/mL), we observed 12 bands for WT fibrils and only 8 bands for E83Q fibrils (Figs. 4K and S5G-H). These differences in the ProK proteolysis profile of E83Q aSyn fibrils are indicative of a distinct fibrillar structure. Taking together, the data from AFM/TEM imaging and ProK digestion analyses suggest that the fibrils generated from the E83Q-mutated aSyn exhibit distinct morphological and structural properties from their WT counterparts.

Next, to further probe the differences in the molecular structure of amyloid fibrils formed by E83Q and WT aSyn, we prepared ^13^C, ^15^N-labeled proteins (Fig. S5I-J) and characterized the aggregated proteins, which were prepared under identical conditions (Fig. S5F, K-L), using ssNMR spectroscopy. We observed high-resolution two-dimensional ^13^C-^13^C dipolar-assisted rotational resonance (DARR) and ^15^N-^13^Cα (NCA) spectra for both WT and E83Q aSyn (Fig. 5 A-B). DARR and NCA experiments employ cross-polarization steps and therefore only detect residues from the rigid cross-*β*-structure core of amyloid fibrils. The high quality of the DARR and NCA spectra demonstrates that both E83Q and WT aSyn aggregate into structurally well-defined amyloid fibrils. Moreover, detailed comparison of the cross-peak patterns observed for the two proteins reveals pronounced differences in the position and intensity of many signals. This is particularly apparent in the spectral region of the ^13^C-^13^C DARR spectrum, in which Cα-C*β* cross-peaks of threonine and serine residues from the rigid fibrillar core appear (Fig. 5A, zoom). The comparison provides residue-specific support for differences in the molecular structure of the cross-*β*-structure core of amyloid fibrils formed by E83Q and WT aSyn.

**Fig. 5.**
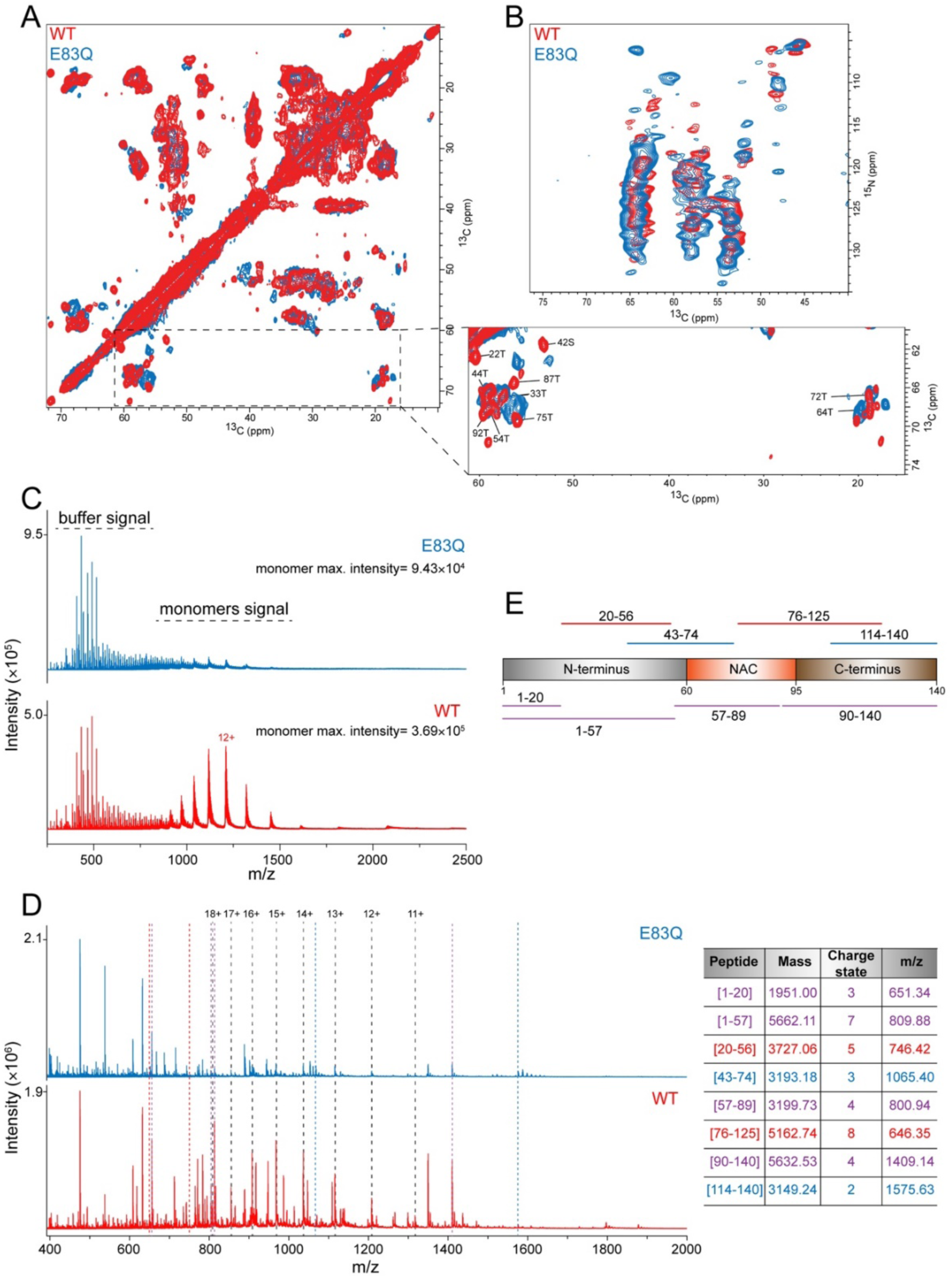
ssNMR and nESI-MS experiments on WT and E83Q fibrils. **(A and B)** Superposition of two-dimensional ^13^C-^13^C DARR **(A)** and NCA spectra **(B)** of E83Q aSyn (blue) and WT (red) aSyn fibrils. Tentative assignments, which were transferred from previous resonance assignments (BMRB id: 18860), of C*α*-C*β* cross peaks of threonine and serine residues in WT aSyn are indicated in the zoom in of (A). **(C)** Comparison of intensities of monomers releasing from the fibril during the nESI-MS experiments on fibrils. We also observed the signals of buffer components at a similar intensity between samples. **(D)** Peptide spectra of WT and E83Q fibrils after 5 minutes digestion with 0.5 µg/µL proK. Peaks related to intact monomer charge states are indicated with black dashed lines. The presence of undigested monomer in both cases, even at very low intensity, indicates that secondary cleavage of peptides is kept to a minimum which is desired to retain structural information from the fibril. Colored dotted lines indicate selected peaks that are present in both conditions (purple), only for WT (red) or only for E83Q (blue). The table lists the selected peaks with their respective mass and linked peptide fragments. Colors are identical to those of the dotted lines in B. **(E)** A scheme depicts the full sequence of aSyn and the peptides (from D) commonly found for both WT and E83Q (purple), while others were unique to one of the two fibrils (red for WT and blue for E83Q).

Finally, to investigate the structural basis underlying the differences in the ProK digestion profile of E83Q and WT aSyn fibrils dynamics, we performed nESI-MS on the digestion products. We first analyzed the fibrils without ProK treatment. Interestingly, we also observed a peak pattern corresponding to monomers, suggesting their release from the fibrils of WT and E83Q aSyn, respectively. It has been shown in other amyloid systems that a dynamic equilibrium exists at the fibril ends involving the dissociation and re-association of the monomers (*39, 40*). Here, the stark difference in the concentration of monomers released (∼4-fold) indicates a difference in the fibrils’ stability, suggesting that monomers were released from WT aSyn fibril ends more freely than those released from E83Q aSyn fibril ends (Fig. 5C).

We next analyzed the WT and E83Q aSyn fibrils after ProK treatment and detected several peptide peaks (Fig. 5D). The observed peak pattern differed between the WT and E83Q fibrils, with more intense peptide signals seen in the WT fibrils. Following manual deconvolution of the peaks, peptide masses could then be linked to amino acid sequences using the PAWS tool. We observed differences in the peptide regions when mapping the identified peptides onto the aSyn protein sequence, indicating structural discrepancies between WT and E83Q fibrils. Interestingly, we detected peptides commonly found for both WT and E83Q aSyn (representative peptides in purple), while others were unique to one of the two fibrils (representative peptides in red for WT and blue for E83Q, Fig. 5D-E).

The peptides identified in both WT and E83Q aSyn fibrils were derived from the first half of the N-terminus [amino acids (aa) 1–20], the entire N-terminus (aa 1–57), in addition to others encompassing nearly the complete NAC domain (aa 57–89) or the fully intact C-terminus (aa 90–140). In the case of WT aSyn fibrils, peptides were detected that derived from the second half of the N-terminus (aa 20–56), and one peptide was detected that covered a part of the NAC region up to the first half of the C-terminus (aa 76–125). Only the E83Q aSyn fibrils showed a peptide covering the last part of the N-terminal region together with the first half of the NAC- domain (aa 43–74) and a peptide that contained the last part of the C-terminus (aa 114–140). Altogether, we identified peptides covering the three distinct domains of the aSyn sequence as intact fragments and unique peptides specific for either WT or E83Q aSyn, indicating differences in the accessibility of specific cleavage sites between both structures, resulting from the structural differences between the fibrils. These findings point to significant differences in the dynamics, stability, and structural properties between E83Q and WT aSyn fibrils.

### E83Q, but not WT, aSyn is toxic to mammalian cells and increases the formation of puncta structures

To determine whether the E83Q mutation enhances the aggregation propensity of aSyn and results in the formation of intracellular inclusions, we performed immunocytochemistry (ICC) and confocal imaging in M17, HEK293, and HeLa cells overexpressing WT or E83Q aSyn. The aggregate formation was assessed via ICC between 24 h and 96 h post-transfection. Co-staining of the cells using total aSyn antibodies (1-20 or 134-138) in combination with pS129 antibodies did not reveal the presence of the typical large aSyn aggregates in any of the three mammalian cell lines overexpressing WT aSyn (Figs. 6A, S6A-B). The overexpression of E83Q aSyn was insufficient to induce spontaneous aggregation of aSyn in either neuronal-like (M17) or non-neuronal-like mammalian cell lines (HEK293 and HeLa; Figs. 6A and S6A-B). Furthermore, Western blot (WB) analyses showed that aSyn carrying the E83Q mutation was not found significantly enriched in the insoluble cellular fraction (Fig. 6B). Besides, small puncta were consistently observed in cells overexpressing E83Q aSyn, but not in those transfected with WT aSyn. Among the three cell lines tested, these puncta structures were preferentially formed in HeLa cells (Figs. 6B and S6H). However, our ICC demonstrated that these puncta were not positive for pS129, a known marker for aSyn fibrils and pathological aggregates (Figs. 6A and S6B). WB analyses also confirmed the absence of a pS129 signal in the insoluble fractions of the cells overexpressing E83Q aSyn (Figs. 6B and S6C-D).

**Fig. 6.**
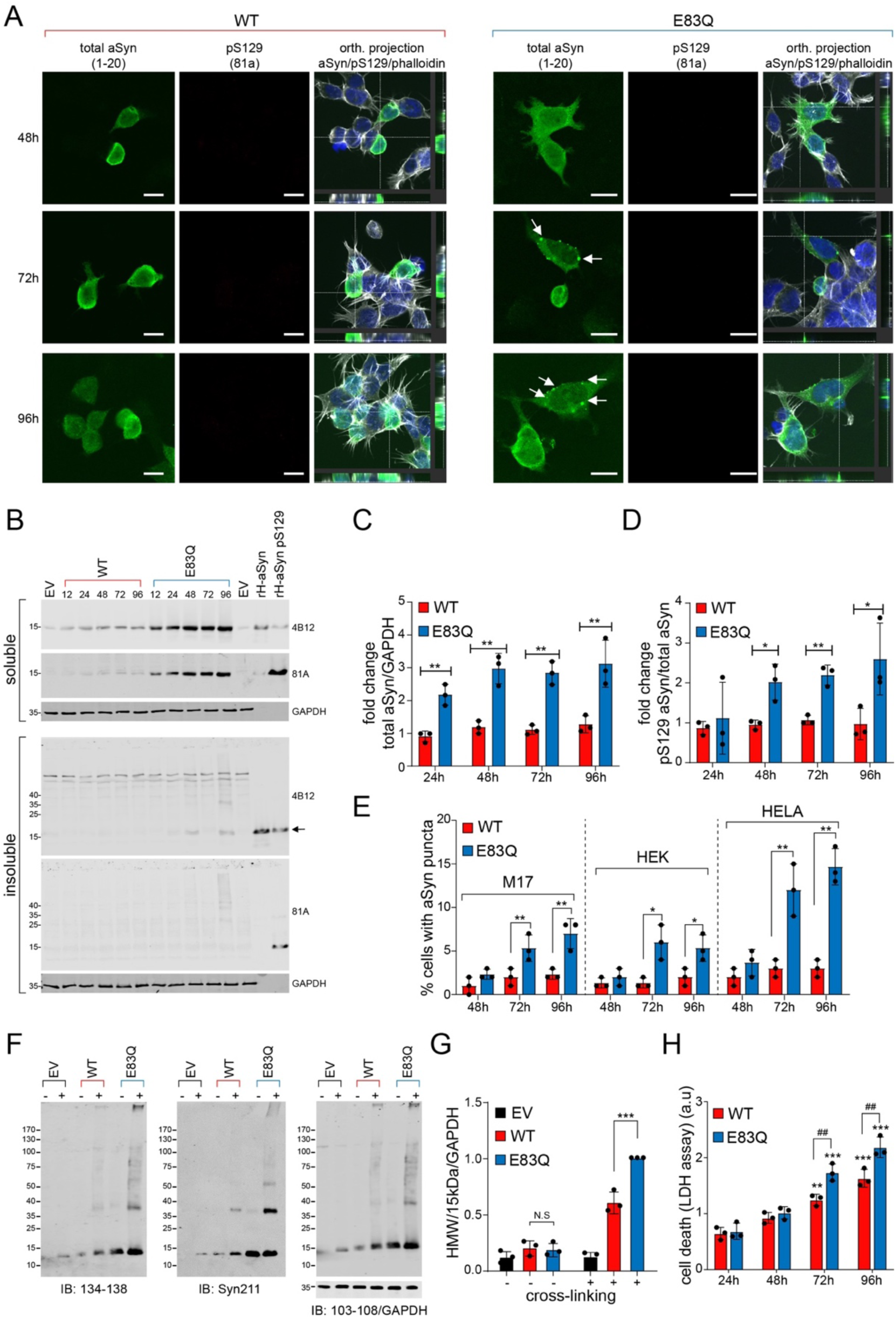
Overexpression of E83Q aSyn in immortal mammalian cells does not significantly alter cellular properties. **(A)** ICC of M17 cells transfected with either WT or E83Q aSyn plasmids for the indicated time. The aSyn inclusions were not detected in these cells (larger fields of view are depicted in Fig. S6H). White arrows point to the puncta-like structures. Scale bars = 10 µM. **(B)** WB of detergent soluble and insoluble fractions of lysed M17 cells transfected with either WT or E83Q aSyn plasmids for the indicated time. The low concentration of aSyn in the insoluble fractions from E83Q-expressing cells is indicated by arrows. **(C)** Graph showing the expression levels of total WT and E83Q aSyn from the M17 soluble fractions (n=3). **(D)** Graph showing the pS129 levels in WT aSyn and E83Q expressing M17 soluble fraction (n=3). **(E)** Graph showing the % of cells with WT and E83Q aSyn puncta (as indicated in A by white arrows) from all three mammalian cell lines. C–E represent the mean ± SD of a minimum of 3 independent experiments. \**p*<0.05, \*\**p*<0.05, \*\*\**p*<0.0005 (ANOVA followed by Tukey [HSD] post-hoc test, WT vs E83Q at each time-point). **(F)** WB analysis following the cross-linking (DSG) analysis of HeLa cells overexpressing either an empty vector (EV), WT aSyn, or E83Q aSyn constructs. High molecular weight (HMW) species of aSyn are observed in the overexpression of WT and E83Q aSyn in the presence of a cross-linker. **(G)** Graph showing the quantification of HMW aSyn bands detected at and above 25 kDa to the top of the gel from F. Data represent the mean ± SD of 3 independent experiments. N.S (non-significant), \*\*\**p*<0.0005 [ANOVA followed by Tukey [HSD] post-hoc test, non-treated cells (WT vs. E83Q) and treated cells with DSG (WT+ vs E83Q+)] in both cell lines. “+” cells were treated with 1 mM DSG; “–” cells with DMSO. **(H)** Graph displaying the aSyn-mediated toxicity measured by quantifying the concentration of LDH released into the HeLa cell culture media at the indicated times. Data represent the mean ± SD of 3 independent experiments. \**p*<0.05 [ANOVA followed by Tukey [HSD] post-hoc test, (WT vs. E83Q).

On the other hand, it was noteworthy that aSyn was significantly upregulated in the soluble cellular fraction of the cell lines overexpressing E83Q (Figs. 6B-D, S6C-D and F-G). Altogether, our data show that the E83Q mutation did not promote the formation of pathological-like pS129-positive aggregates in mammalian cell lines; it did, however, increase the number of small, dot-like structures (Fig. 6E), which may represent the accumulation of multimeric/oligomeric species. This finding possibly reflects a greater propensity of E83Q to form oligomers (e.g., dimers and tetramers) intracellularly.

To test this hypothesis and determine the size distribution of the species formed in these puncta structures, we conducted size-exclusion chromatography (SEC) on the cellular extracts from HEK and HeLa cells overexpressing either WT or E83Q aSyn. WB analyses of the SEC fractions confirmed that both E83Q and WT aSyn species were mostly detected as monomers (∼15 kDa; Fig. S7). To investigate the possibility that the E83Q oligomers could be unstable and disassociate on the column, we performed protein cross-linking (*41–44*). The SEC analyses of the cellular extracts treated with disuccinimidyl glutarate (DSG) confirmed that in both cell lines overexpressing WT or E83Q, the majority of aSyn species eluted in a similar volume as the unfolded human recombinant aSyn (Fig. S8A-B). We then performed WB analyses to assess the level of oligomerization in cells. In the control cells [empty vector (EV)] treated with DSG or DMSO, the monomeric aSyn was either not observed or only weakly detected, and no oligomeric bands were detected (Figs. 6F and S8C).

In the absence of the cross-linking agent, WT and E83Q aSyn overexpressed in HeLa and HEK cells were both detected as a prominent single band (∼15 kDa) corresponding to the molecular weight of the aSyn monomer. However, DSG treatment of the cells overexpressing WT aSyn revealed the presence of several high-molecular-weight (HMW) aSyn species, including a smear above 130 kDa, in addition to the main aSyn monomer band, as previously described in cross-linking studies (*43, 45*). The HMW bands and smear above 130 kDa were also detected in the soluble fraction of the cells overexpressing E83Q, with a similar MW as the WT counterparts. As the E83Q mutant is expressed at the protein level, but not at the mRNA level (Fig. S6E), at higher steady-state levels than aSyn WT in HeLa cells (Fig. 6B), the level of the HMW signal was normalized to the total aSyn level expressed in cells. This showed significantly higher levels of the HMW aSyn species in cells overexpressing E83Q (Figs. 6F-G and S8C). Although the HMW species, such as the dimers and oligomers, seem to be minor species, our data suggest increased oligomerization within cells, especially when overexpressing E83Q aSyn. Finally, we assessed cell viability over time in mammalian cells overexpressing WT or E83Q aSyn using the lactate dehydrogenase (LDH) toxicity assay. Our cell death assay showed that overexpression of aSyn WT and E83Q induced a significant increase in toxicity over time, as evidenced by an increase in the loss of plasma membrane permeability from 96h for WT overexpressing cells and starting already at 72h for E83Q cells (Figs. 6H and S6I). Moreover, in both M17 and HEK cells, E83Q overexpression induced higher cell death than WT aSyn at 96h (Fig. S6I). Such differences could be attributed either to the higher expression levels of E83Q in these cells or to the greater propensity of E83Q aSyn to oligomerize. The HMW bands and smear above 130 kDa were also detected in the soluble fraction of the cells overexpressing E83Q, with a similar MW as the WT counterparts. It remains unclear if such differences could be attributed solely to the higher expression levels of E83Q in these cells or its greater propensity of E83Q aSyn to oligomerize, or both.

### WT and E83Q fibrils preferentially seed the aggregation of monomers with the same sequence

Since the E83Q mutation carrier was heterozygous, we also investigated the aggregation of the E83Q mutant in the presence of WT aSyn at varying molar ratios (4:1, 1:1 and 1:4). As shown in Fig. 7A, the presence of WT aSyn monomers resulted in a significant and concentration-dependent delay in the aggregation of E83Q monomers, as evidenced by the increase in lag time from ∼0.9h for only E83Q samples to ∼3.9h, ∼8.4h and ∼10.9h for samples containing a mixture of 4:1, 1:4 and 1:1 ratio of E83Q: WT aSyn, respectively. EM analysis of the samples at the end of the aggregation process showed fibrils in all the E83Q: WT aSyn mixtures (Fig. 7C). The fibrils formed in these mixtures exhibited a higher propensity to undergo lateral association. Interestingly, the samples containing an equimolar concentration of WT and E83Q monomers showed a multiphasic aggregation profile (Fig. 7A), suggesting a more complex aggregation process involving a complex interplay between different species of both proteins. Despite this, only fibrils were observed at the end of the aggregation process (Fig. 7C).

**Fig. 7.**
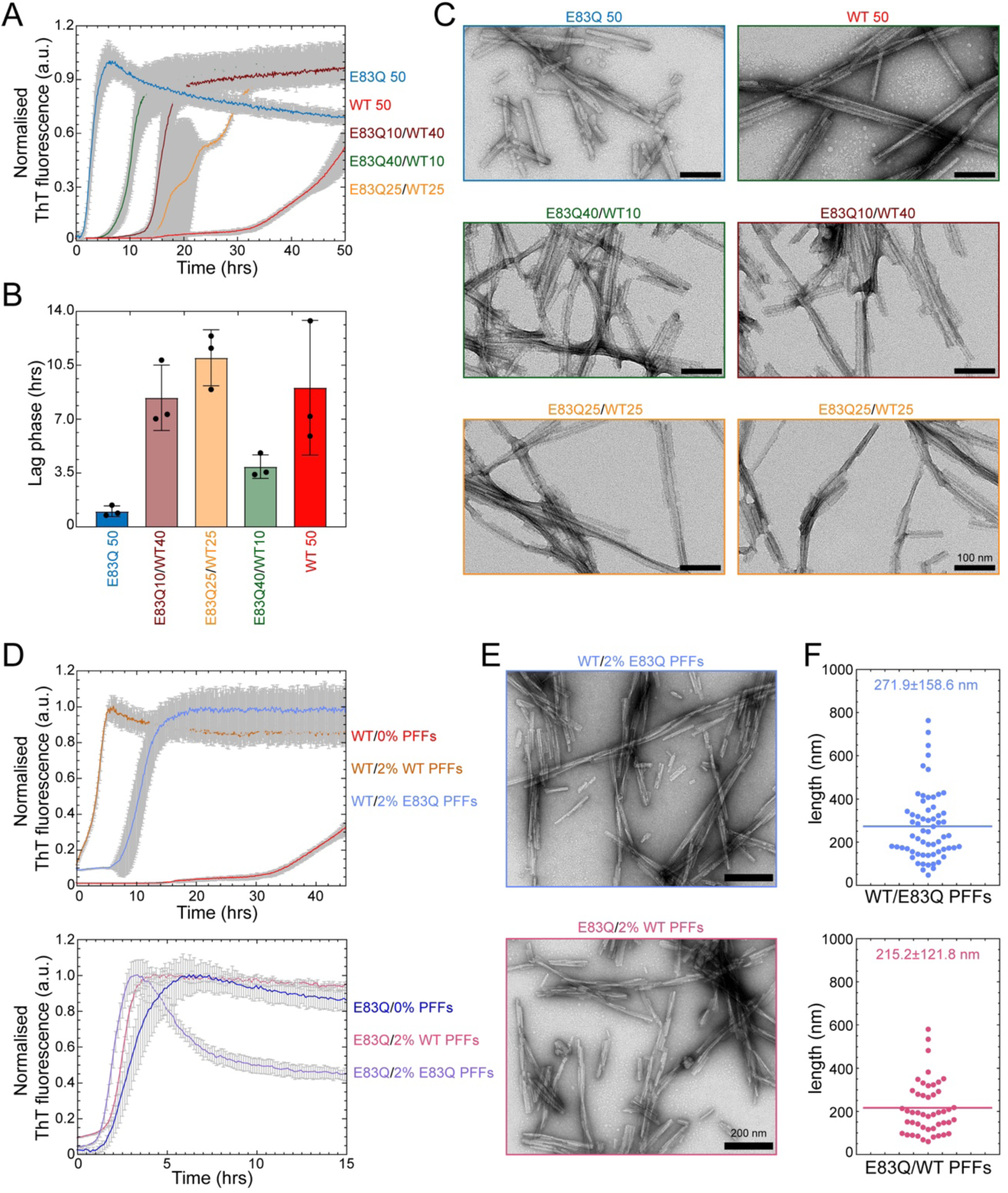
WT and E83Q fibrils preferentially seed the aggregation of monomers with the same sequence. **(A)** ThT based aggregation kinetics of cross-species aggregation of hWT monomers with hE83Q monomers at varying ratios. **(B)** Bar diagram showing the lag phase (hrs) extracted from the aggregation kinetics (from A). **(C)** Negatively stained EM images of the fibril samples at the end time-point of the ThT kinetics (from A) (Scale bar: 100 nm). **(D)** ThT based aggregation kinetics of cross-seeding of 50 μM concentration of hWT monomers (top) or hE83Q monomers (bottom) with 2% of hWT PFFs or h83Q PFFs. **(E-F)** EM analysis of end time-point (from E) samples of hWT monomers seeded with 2% hE83Q PFFs (top) and hE83Q monomers seeded with hWT PFFs (bottom) and their fibrils length distribution (F), respectively. Scale bars = 200 nm.

Next, we investigated and compared the ability of WT and E83Q fibrils to seed the aggregation of monomers of both proteins. The primary objective of this experiment was to determine if E83Q PFFs are able to transmit their morphological and structural properties to the next generation of fibrils and to assess the specificity and efficiency of cross-seeding by WT and E83Q PFFs. As shown in Fig. 7D, seeding by PFFs was always more efficient when both the PFF seeds and monomers were of the same sequence. WT aSyn monomers are more efficiently seeded by WT PFFs than E83Q PFFs (Fig. 7D, top panel). Similarly, the E83Q monomers are more efficiently seeded by E83Q PFFs (Fig. 7D, lower panel). EM analysis of fibril length in the two-seeded aggregation samples (Fig. 7E-F) showed the WT monomers seeded with E83Q PFFs resulted in fibrils with an average length similar to those observed for the aggregation of E83Q monomers alone. These observations suggest that the E83Q fibrils exhibit distinct conformations, preferentially seed E83Q monomers, and are able to pass some of their morphological and structural properties to WT aSyn monomers.

### Human E83Q aSyn PFFs show high seeding activity and increased toxicity in primary mouse neurons

Our *in vitro* biochemical and biophysical experiments established that the E83Q mutation alters the morphology, stability, and structural properties of aSyn fibrils (Figs. 3 and 4). Therefore, we sought to determine to what extent the E83Q mutation could influence the seeding activity of the pre-formed aSyn fibrils (biophysical characterization of the PFF preparations is shown in Fig. S9) in neurons. Given that postmortem neuropathological characterization of the brain of the E83Q mutation, carrier showed significantly higher LB pathology in the hippocampal and cortical areas than the substantia nigra (*18*), we compared the seeding activity of WT and E83Q PFFs in WT hippocampal and cortical primary neuronal cultures. Previous studies from our laboratory (*46*) and others (*47, 48*) have shown that aSyn PFFs seeding activity correlates with levels of aSyn expression in different neuron types, which explains why aSyn PFFs seed more efficiently in hippocampal neurons compared to cortical neurons, which express lower levels of aSyn. We used a neuronal seeding model, in which the addition of nanomolar concentration of PFFs primary neuronal culture triggers the formation of intracellular aggregates of endogenously expressed aSyn (Fig. S10) (*22–24*). We also sought to compare the seeding activity of WT and E83Q PFFs in DA neurons, but we failed to observe any seeding activity (data not shown). Therefore, we limited our seeding activity studies to hippocampal and cortical neurons.

As the seeding process requires the internalization of the PFFs into the neurons, we first determined whether the conformational properties of the E83Q PFFs alter the uptake or processing of human aSyn PFFs (e.g., phosphorylation or C-terminal cleavage of PFF) (*46*). To follow the fate of the PFFs independently of the seeding mechanism, we used aSyn knockout (KO) neurons in which the PFFs are unable to induce seeding due to the absence of endogenous aSyn (Fig. S11) (*22, 46*). ICC combined with confocal imaging confirmed that both human WT (hWT) and human E83Q (hE83Q) PFFs were readily internalized in the neurons via the endolysosomal pathway (LAMP1 positive vesicles) 24 h after their addition to the primary cultures (Fig. S11A) as previously reported in mouse PFF-treated neurons (*46, 49, 50*).

Next, we quantified the level of PFF internalization by quantifying the amount of internalized HMW (*22, 46*). The PFF levels were significantly higher in the insoluble fraction of the hE83Q PFF-treated KO neurons than in neurons treated with hWT PFFs, as indicated by the HMW band detected by the pan-synuclein antibody (SYN-1) (Fig. S11B-C). We next investigated whether the E83Q mutation alters the processing of the internalized PFFs. Within 24 h, both hWT and hE83Q PFFs were C-terminally truncated, as evidenced by the detection of the 12 kDa band. After 3 days, both hWT and hE83Q PFFs were fully cleaved, as shown by the complete loss of full-length aSyn (15 kDa; Fig. S11C). The truncated species were cleared over time, but a residual level remained up to 21 days after internalization of the PFFs into the neurons for both hWT and hE83Q PFFs (Fig. S11C). Finally, the internalized hWT or hE83Q PFFs were never phosphorylated at residue S129 in KO neurons (Fig. S11C). Altogether, our results demonstrate that the conformational properties of E83Q PFFs increase their uptake into neurons without interfering with the processing and clearance once internalized.

Next, we compared the seeding capacity of the hE83Q and hWT PFFs in WT murine hippocampal neurons. hE83Q or hWT PFFs were added to WT primary culture, and their seeding activity and ability to induce the formation of LB-like inclusions were investigated at day 7 (D7), D14, and D21 of treatment. As previously reported by our group (*22*) and others (*51*), the internalized PFF seeds never get phosphorylated at S129 residue at early or late time-points, up to 21 days, and the great majority are cleaved at residue 114 within 12-24 hours (*46*). Therefore, they do not interfere with the detection and quantification of newly formed fibrils by pS129 antibodies. ICC confirmed the formation of aggregates immunoreactive to aSyn pS129, both in hWT or hE83Q PFF-treated neurons at D14 and D21 (Fig. 8A). Quantification of the pS129 level by high content imaging (HCA) demonstrated that more pS129-seeded aggregates were formed in neurons treated with hE83Q PFFs than in those treated with hWT PFFs, both at D14 and D21 (Fig. 8B). Interestingly, the level of pS129-seeded aggregates had increased even further from D14 to D21 in hE83Q PFF-treated neurons but not in hWT PFF-treated neurons. Like the hippocampal neurons, the addition of hE83Q PFF to the cortical neurons induces the formation of pS129-seeded aggregates that accumulate over time (Figs. 8C and S12A-B). The level of pS129-seeded aggregates was also significantly higher in hE83Q PFF-treated cortical neurons than in the hWT PFF-treated cortical neurons at D14 and D21 (Figs. 8C and S12A-B). Altogether our findings suggest that the E83Q mutation strongly enhances the seeding activity of human aSyn PFFs in both the hippocampal and cortical neurons.

**Fig. 8.**
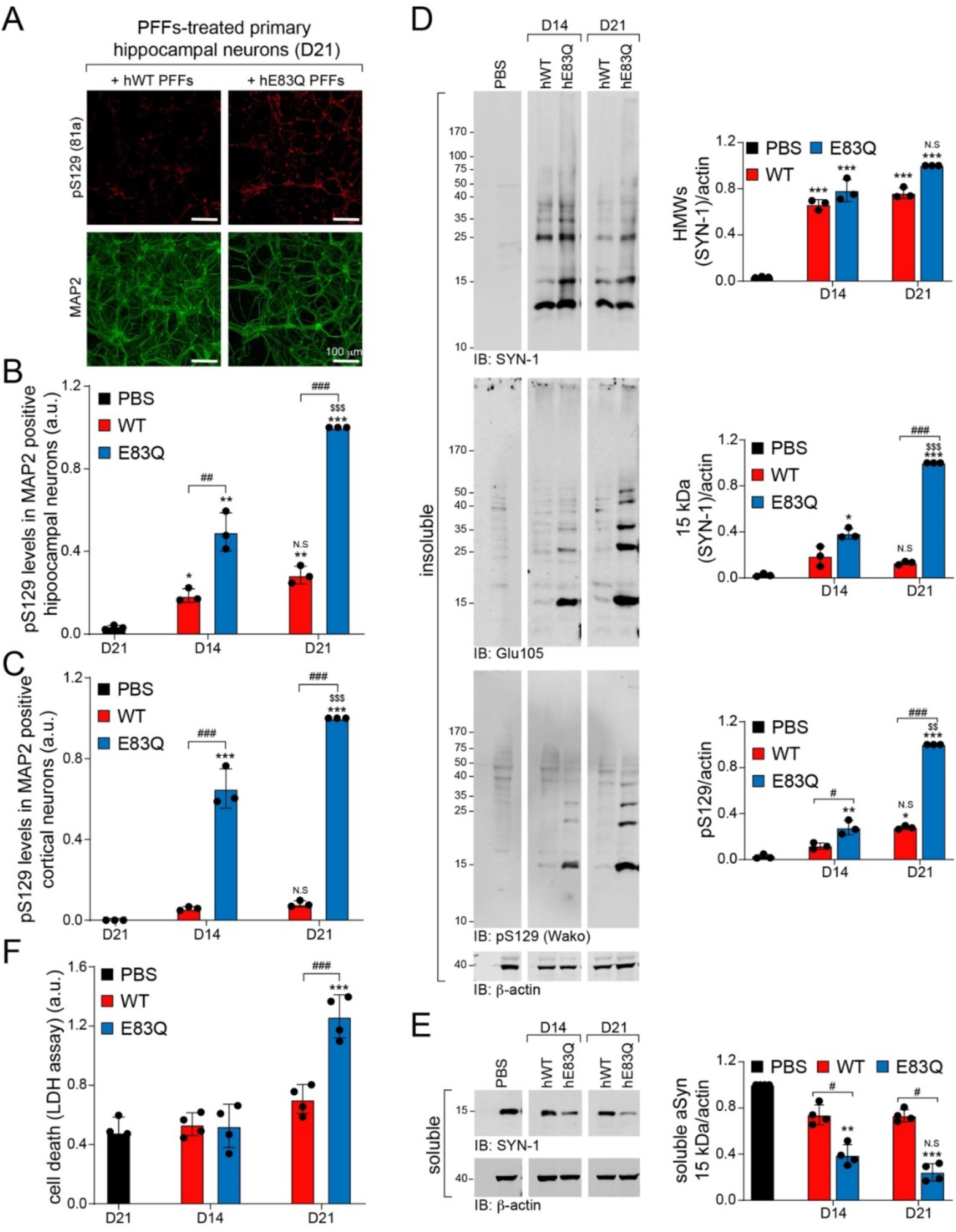
Human E83Q PFFs have a higher seeding capacity than human WT PFFs in WT hippocampal primary neurons. Temporal analysis of the level of aSyn seeded-aggregates formed in WT hippocampal neurons after the addition of 70 nM of hWT or hE83Q aSyn PFFs to the primary culture (at DIV 5) for 14 days (D14) or D21. Control neurons were treated with PBS. **(A-C)** The level of seeded aggregates was measured over time in PBS- or PFF-treated hippocampal (A-B) or cortical (C) neurons by high content analysis (HCA). Seeded-aggregates were detected by ICC using pS129 (81a) antibody, neurons were counterstained with microtubule-associated protein (MAP2) antibody, and nuclei were counterstained with DAPI. Scale bars = 100 μm. For each independent experiment (n=3), a minimum of two wells was acquired per condition, and nine fields of view were imaged per well. **(D-E)** Western blot of total aSyn, pS129, and actin, as detected by SYN-1, Glu105 (antibody specific for the detection of mouse aSyn), pS129 Wako, or pS129 MJFR-13 antibodies, respectively. In the insoluble fraction, HMW bands corresponding to the newly formed fibrils are detected from 25 kDa up to the top of the gel. **(F)** Cell death levels were assessed in WT neurons treated with PFFs (70 nm) at D14 and D21 using lactate dehydrogenase (LDH) release assay (n=≥3 experiments). For each independent experiment (n=5), triplicate wells were measured per condition. **(B–F)** The graphs represent the mean +/- SD of a minimum of 3 independent experiments. p<0.05=*, p<0.005=**, p<0.0005=*** (ANOVA followed by Tukey [HSD] post-hoc test, PBS vs. PFF-treated neurons). p<0.05=#, p<0.0005=### (ANOVA followed by Tukey [HSD] post-hoc test, hWT PFF-treated neurons vs. hE83Q PFF-treated neurons). p<0.005=$$, p<0.0005=$$$ (ANOVA followed by Tukey [HSD] post-hoc test, D14 vs. D21 PFF-treated neurons). N.S (non-specific, D14 vs. D21).

We further characterized the seeding level in PFF-treated neurons by WB analysis. As shown in Fig. S12C and reported previously, the aSyn PFFs seeding level is significantly higher in hippocampal neurons than in cortical neurons. Therefore, we performed the rest of our studies on the biochemical and morphological properties of the newly formed aggregates in primary hippocampal neurons. Given that SYN-1 detects both the PFFs and the seeded aggregates in the insoluble fraction, the pS129 antibody was used to discriminate the newly formed aSyn aggregates from the exogenously added PFFs, as the latter undergo C-terminal cleavage and are not subjected to phosphorylation at S129 in the neuronal seeding model (*22, 46, 52*). Consistent with the ICC and HCA measurements, pS129- positive seeded aggregates were barely detected in the insoluble fraction of the hWT PFF-treated neurons. This confirms the low degree of seeding in hWT-PFF treated neurons, which is at the limit of the WB software detection threshold (*22, 46, 52*).

Conversely, we observed an accumulation of aSyn-seeded aggregates in the insoluble fraction of the hE83Q PFF-treated neurons (Fig. 8D), which were positively stained with SYN-1, which recognizes the 15 kDa and HMW bands at ∼23, 37, 40, and 50 kDa, and with a specific antibody against pS129, which recognizes the 15 kDa band and the HMW bands at ∼23 and 35 kDa). As measured by HCA, the level of pS129 significantly increased between D14 and D21 in hE83Q PFF-treated neurons. Conversely, pS129-positive seeded aggregates were barely detected at D14 or D21 in hWT PFF-treated neurons (Fig. 8D). The detection of the pS129-positive seeded aggregates in the insoluble fraction was concomitant with the shift of endogenous aSyn from the soluble (Fig. 8E) to the insoluble fraction in the hE83Q PFF-treated neurons. Nevertheless, and despite a trend towards a downregulation of aSyn in the soluble fraction between D14 and D21, the difference in aSyn signal (SYN-1) was not significant (Fig. 8E).

Finally, LDH toxicity assays revealed neuronal cell death at D21 in the hE83Q-PFF treated neurons but not in hWT PFF-treated neurons (Fig. 8F). Altogether, our findings demonstrate that the E83Q-dependent changes in the structural and dynamic properties of human aSyn PFFs translate into increased uptake, seeding activity, and neurotoxicity.

### The E83Q mutation restores the capacity of human aSyn PFFs to induce the formation of LB-like inclusion in neurons

Next, we investigated and compared the shape and morphology of the pS129-positive seeded aggregates formed upon the addition of hWT or hE83Q PFF to the primary culture. It is well-established that human fibrils less efficiently seed the aggregation of endogenous mouse aSyn in primary neurons (*53, 54*). Therefore, the formation and the evolution of aSyn-seeded aggregates in mice primary neurons have been extensively studied using mouse aSyn PFFs (mWT PFFs) (*22-24, 46, 47, 50, 51, 53, 55-63*).

Previously, we and others have shown that the seeded aggregates, up to 14 days of formation, appeared predominantly as long filamentous-like structures that evolve into inclusions at D21, with different structural and morphological properties that we classified as filamentous (∼30%), ribbon-like (∼45%), or round LB-like inclusions (∼ 22%) (*22*). In studies using hWT PFFs to induce seeding in mouse primary neurons, the pS129-positive seeded aggregates were mostly detected as small puncta in neurites and as long fibrils in the cell bodies (*24, 52, 64-67*). The morphology of hWT PFF-seeded aggregates has never been assessed after D14. Therefore, we compared the morphology of aSyn inclusions formed by the human WT or E83Q seeds to that induced by the addition of mWT PFFs up to D21.

Firstly, based on pS129 staining, confocal imaging revealed that human aSyn PFF induced the formation of seeded-aggregates with different types of morphologies than those detected in mWT PFF-treated neurons. The morphological classification of the different types of seeded-aggregates formed in human or mouse PFF-treated neurons is depicted in Figs. 9A and S13A and their relative distribution at D14 and D21, respectively, in Figs. S13B (D14) and 9B (D21).

**Fig. 9.**
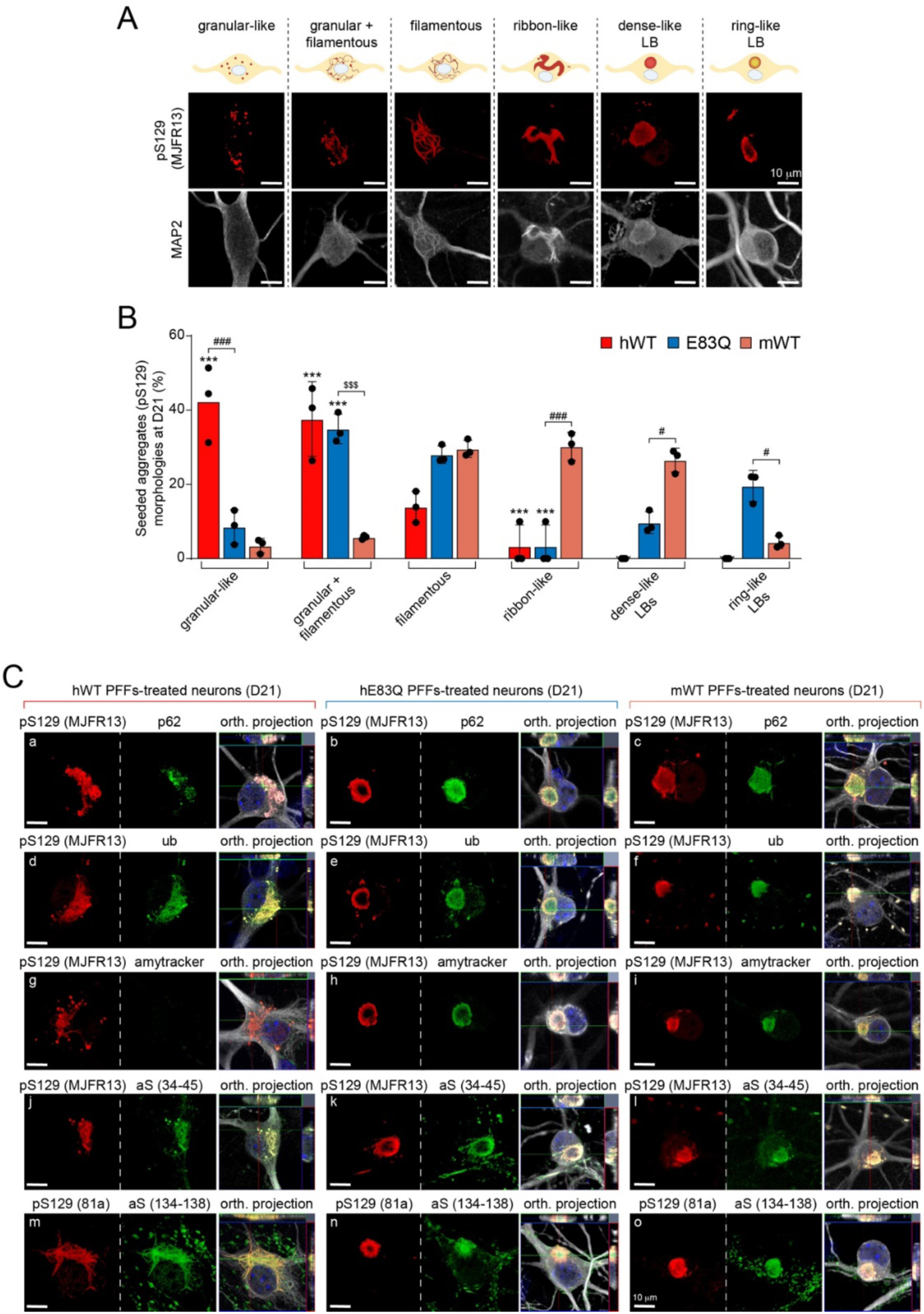
Comparative analysis of the morphological properties and diversity of aSyn aggregates and inclusions formed at 21 days after PFF treatment in hippocampal neurons. **(A)** LB-like pathological diversity was observed in PFF-treated hippocampal neurons. The representative images for the filamentous and the ribbon-like inclusions shown in this panel are also depicted in supplementary Figs. S13-S16 with additional markers of the LB-like inclusions. **(B)** Quantification of the morphologies of the aSyn seeded-aggregates observed by ICC in hippocampal primary neurons at D21. A minimum of 100 neurons was counted in each experiment (n=3). The graphs represent the mean +/- SD of a minimum of three independent experiments. p<0.0005=*** (ANOVA followed by Tukey [HSD] post-hoc test, mWT PFF-treated neurons vs. human PFF (hWT or E83Q)-treated neurons). p<0.0005=### (ANOVA followed by Tukey [HSD] post-hoc test, hE83Q PFF-treated neurons vs. mWT PFF or hWT PFF-treated neurons). p<0.0005=$$$ (ANOVA followed by Tukey [HSD] post-hoc test, hWT PFF-treated neurons vs. hE83Q PFF-treated neurons). **(C)** Aggregates were detected by ICC using pS129 (MJFR13 or 81a) in combination with p62 (a–c), ubiquitin (d– f), Amytracker dye (g–i), total N-ter aSyn (epitope: 34-45) (j–l), or total C-ter (epitope: 134-138) (m–o) aSyn antibodies in hWT PFFs (a, d, g, j, m), hE83Q PFFs (b, e, h, k, n), or mWT PFF-treated hippocampal neurons (c, f, i, l, o). Neurons were counterstained with MAP2 antibody and the nuclei with DAPI. Scale bars = 10 μm.

In mWT PFF-treated neurons at D14, the seeded aggregates appeared predominantly as filamentous-like inclusions (∼75%; Figs. S13, S14A-Ca,b, S14Ba,b) detected both in the neurites and neuronal cell bodies (*22, 24*). In addition to these filamentous aggregates (Figs. S14Ad, S14Bd,g), we observed the formation of aggregates that either exclusively appeared as granular-like aggregates or as a mixture of granular and filamentous aggregates at D14 (Figs. S13B, S14Ae-i, S14Bf,h-i, S13De,h,i) and D21 (Figs. 9B, S14Da,d,g,j, S14Ed,e,h,i, S14Ed,e,h,i, S14Fe,h,i) for human hWT and hE83Q PFF-treated neurons. However, these types of aggregates were not observed in mWT PFF-treated neurons at any time point.

In human WT or E83Q PFF-treated neurons, the ribbon-like structures, which are commonly detected in mouse PFF-treated neurons (∼10% of the seeded aggregates at D14 and ∼30% at D21 (*22*)) were not observed either at D14 (Figs. S13B, S14A-C) or D21 (Figs. 9B and S14Bc, Db, Gb). More importantly, while the LB-like inclusions were formed at a similar level in hE83Q PFF-treated as in mWT PFF-treated neurons at D14 (∼10%) (Fig. S13B) and D21 (∼28%; Fig. 9B), no LB-like inclusions were ever observed in hWT PFF-treated neurons (Figs. 9B and S13-16).

Altogether, our data suggest that the specific conformation adopted by the PFFs generated *in vitro* from aSyn carrying the E83Q mutation, as shown in our *in vitro* experiments, appears to interact more favorably with endogenous mouse aSyn monomers, leading to their efficient recruitment to fibrils. These *de novo* fibrils were able to mature into pathological LB-like inclusions in both cortical and hippocampal primary neurons. These findings show that this single E83Q mutation within the NAC region dramatically alters the structure and seeding activity of human aSyn PFFs in manners that facilitate the formation of LB-like inclusions, which was not possible with WT human PFFs.

### Seeded-aggregates formed upon the addition of human E83Q PFFs recapitulate the biochemical and architectural properties of classical brainstem LBs in hippocampal neurons

To further characterize the newly formed aSyn aggregates, the following LB-like biochemical properties were evaluated: 1) immunoreactivity at D14 and D21 of aSyn-seeded aggregates to ubiquitin (*68*) and p62 (*69*) (Figs. 9C and S14A-B, D, E); 2) binding to the fluorescent Amytracker tracer dye, which binds specifically to the β-sheet structure of amyloid-like protein aggregates (*70*) (Figs. 9C and S14C, F); and 3) immunoreactivity of aSyn-seeded aggregates to total aSyn (N- and C-terminal antibodies) over time (Figs. 9C and S15). Given that the formation of LB-like inclusions has not yet been investigated in human PFF-treated neurons, we systematically compared the properties and morphological features of the seeded aggregates formed at D14 and D21 in human PFF-treated neurons to those formed in mouse PFF-treated neurons (*22*).

First, all types of seeded-aggregates formed in hWT and hE83Q PFFs or in mWT PFF-treated neurons were positively stained by N- and C-terminal aSyn antibodies (Figs. 9C and S15) or by the ubiquitin antibody (Figs. 9C and S14B, E). While in the neuronal cell bodies of the mWT PFF-treated neurons, p62 (Figs. 9C and S14Aa-c, Da-c) and Amytracker dye (Figs. 9C and S14Ca-c, Fa-c) detected all of the seeded-aggregates, as previously reported (*22*), both markers barely colocalized with the granular-like or the mixture of granular and filamentous aggregates formed after the addition of the human PFFs at D14 (Figs. S14Ae,f,h,i and S14Ce,g-i) and D21 (Figs. 9C, S14De,g-i and S14Fe,g-i). The lack of recognition of these structures by p62 and Amytracker was even more pronounced in the hWT PFF-treated neurons (Figs. 9C, S14Ag–i, S14Cg–i, S14Dg–I and S14Fg–i) than in those treated with the E83Q-mutant PFFs. This suggests that some of the pS129-positive species formed by the human PFFs (WT or E83Q) represent non-fibrillar aggregates (i.e., oligomers).

Our morphological analyses revealed a different organization of the LB-like inclusions formed in mWT or hE83Q PFF-treated neurons. While the proportion of the LB-like inclusions detected in both types of neurons was similar at D14 (∼10%) and D21 (∼28%; Figs. 9B and S13B), a striking difference was seen in the morphology of these inclusions (Fig. S13C-E). In the mWT PFF-treated neurons, most of the LB-like inclusions appeared with a dense core (∼80–90% at D14 and D21; Figs. S13A, C) positively stained by pS129, p62 (Fig. S13Ca), ubiquitin (Fig. S13Cc) and the Amytracker dye (Fig. S13Ce). In rare cases, LB-like inclusions were organized with a ring-like structure (Fig. S13Cb,d,f), as previously observed (*22*). Conversely, in hE83Q PFF-treated neurons, the majority of the LB-like inclusions observed at D14 (∼65%, Figs. 9C and S13A) or D21 (∼70%, Fig. S13D-E) appeared with a ring-like organization in which the core was no longer detected by pS129 antibodies (Fig. S13Db,d,f) or by the Amytracker dye (Fig. S13Df). This may suggest that in these types of LB-like inclusions, aSyn fibrils are preferentially relocated to the periphery rather than the center of the inclusion.

Our findings corroborate those of a recent study that used super-resolution microscopy to show the accumulation of pS129-positive aSyn at the periphery of the nigral LB (*71*). Such organization, with a highly dense and organized shell of phosphorylated aSyn at the periphery of the inclusions, also resembles the brainstem LB inclusions observed in human brain tissues from PD patients at stages 3–5 of disease progression (*25, 26, 30-32*).

As the diversity of human PD pathology extends beyond the classical halo-like LBs or the diffuse cortical LBs, we also assessed the level of Lewy Neurite (LN)-like pathology in the PFF-treated neurons. HCA-based quantification demonstrates that between D14 and D21, the level of the pS129 seeded-aggregates significantly increases in both the neurites and the cell bodies of the hE83Q PFF-treated neurons (Fig. S13F-G). In contrast, in the mPFF- treated neurons, the number of neuronal cell bodies containing pS129-positive aggregates reaches its maximum at D14 (Fig. S13G), suggesting that in these neurons, the main changes observed between D14 and D21 are associated mainly with their structural reorganization and conversion into LB-like inclusions (Figs. 9 and S13).

Finally, the p62 (Fig. S13Db) and ubiquitin (Fig. S13Dd) antibodies recognized not only the periphery of the inclusions but also the center. Interestingly, cytoskeletal proteins such as MAP2 (Fig. S13Db,d,f), neurofilaments (NFLs) (Fig. S16Cf), and the mitochondrial marker Tom20 were also detected both at the periphery and in the center of these inclusions (Fig. S16Df). Our findings align with those of previous studies showing the recruitment and the differential distribution of cytoskeletal proteins and organelles in human LB inclusions from the substantia nigra (*26, 72–76*), the hippocampal CA2 region (*77*), or the stellate ganglion (*78*).

It is noteworthy that recent analysis of the proteome of purified LBs or LB-enriched preparations from human brain tissues (*79–82*) showed that 65-80% of the proteins overlap with the proteome of the LB-like inclusions that form in our neuronal model (unpublished data). Altogether, our findings demonstrate that the PFFs generated from the E83Q mutant induced the formation and maturation of LB-like inclusions in hippocampal neurons that appear to recapitulate some of the morphological, biochemical, and architectural features of LBs found in the brain of patients with PD and related synucleinopathies.

### The E83Q mutation induced the formation of seeded aggregates with distinct morphology in cortical neurons

Finally, we investigated and compared the seeding activity of WT and E83Q PFFs in cortical neurons. As shown in Figure 10, both WT and E83Q human aSyn PFFs induced the formation of pS129 positive aggregates of different morphologies (Fig. 10A-B), similar to those observed in the hippocampal neurons (Fig. 9A-B).

**Fig. 10.**
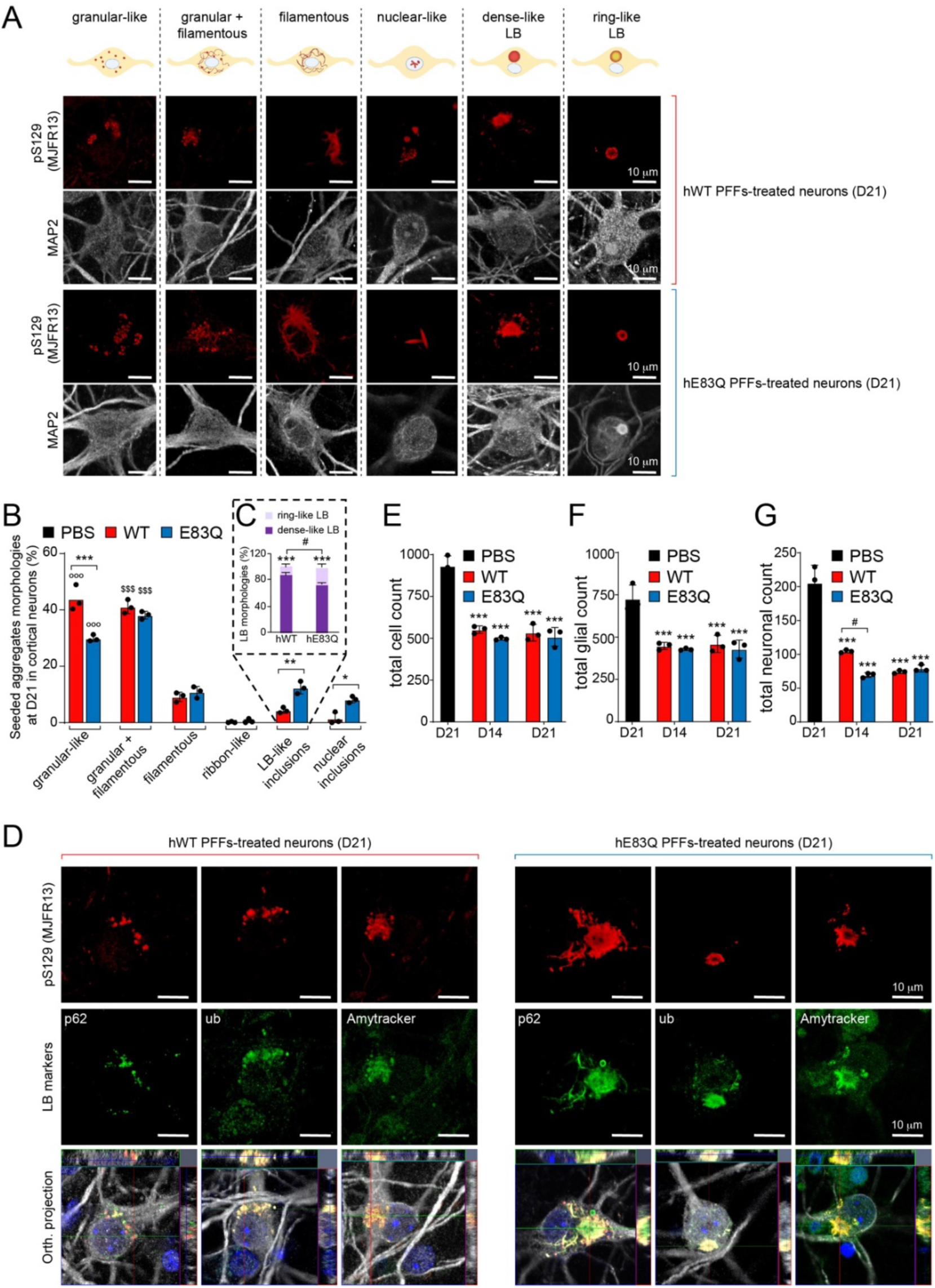
Comparative analysis of the morphological properties and diversity of aSyn aggregates and inclusions formed at 21 days after PFF treatment in cortical neurons. **(A)** LB-like pathological diversity was observed in PFF-treated cortical neurons. The representative images for the different types of seeded aggregates are also shown in Supplementary Fig. S17 with additional markers of the LB-like inclusions. **(B)** Quantification of the morphologies of the aSyn seeded-aggregates observed by ICC in primary neurons at D21. A minimum of 100 neurons was counted in each experiment (n=3). The graphs represent the mean +/- SD of a minimum of three independent experiments. p<0.0005=***, p<0.005 and p<0.05=* (ANOVA followed by Tukey [HSD] post-hoc test, hWT PFF-treated neurons vs. E83Q PFF- treated neurons). p<0.0005=°°° (ANOVA followed by Tukey [HSD] post-hoc test, granular-like aggregates vs. other types of seeded-aggregates). p<0.0005=$$$ (ANOVA followed by Tukey [HSD] post-hoc test, granular and filamentous-like aggregates vs. other types of seeded-aggregates). (**C**). Quantification of the morphologies of the LB-like inclusions (dense core vs. ring-like structure) observed by ICC in primary neurons at D21. A minimum of 50 neurons was counted in each experiment (n=3). The graphs represent the mean +/- SD of a minimum of three independent experiments. p<0.0005=*** (ANOVA followed by Tukey [HSD] post- hoc test, dense core vs. ring-like structure). p<0.0005=### (ANOVA followed by Tukey [HSD] post-hoc test, ring-like structures counted in hWT PFF-treated neurons vs. E83Q PFF-treated neurons). **(D)** Aggregates were detected by ICC using pS129 (MJFR13) in combination with p62, ubiquitin and the Amytracker dye (additional representative images are shown in Fig. S17). Neurons were counterstained with MAP2 antibody and the nuclei with DAPI. Scale bars = 10 μm. The properties and morphological features of the LB-like inclusions formed in the cortical neurons (see also Fig. S17) were similar to that observed in the hippocampal neurons (Figs. 9 and S13-S16). The LB inclusions with a dense core were positively stained by pS129 (see also S17), p62 (see also Fig. S17A), ubiquitin (see also Fig. S17B), and the Amytracker dye (see also, Fig. S17C), while the core of the LB-like inclusions with a ring-like organization was positively stained by p62 and ubiquitin but no longer detected by pS129 antibodies or by the Amytracker dye (see also, Fig. S17). (**E-G**) Quantification of cell death in PBS- and PFF-treated neurons was performed at D14 and D21. At the indicated time, neurons were fixed and ICC performed using NeuN antibody to specifically stain the neuronal nuclei and DAPI staining to label the nuclei of all cells in the primary culture (Fig S18). The graphs represent the mean +/- SD of three independent experiments. For each independent experiment, 3 fields of view (FOV) per condition were acquired at a 10x magnification (∼400 to 1’000 cells per FOV were counted). For each FOV, the count of the total cell population (E) was based on the number of DAPI positive nuclei and the count of neurons (F) on the number of NeuN positive nuclei. Cells were counted as glial (G) if they were DAPI positive and NeuN negative. p<0.0005=*** (ANOVA followed by Tukey [HSD] post-hoc test, PBS vs. PFF- treated neurons). p<0.05=# (ANOVA followed by Tukey [HSD] post-hoc test, hWT PFF-treated neurons vs. E83Q PFF-treated neurons).

The newly formed aggregates in hWT or E83Q PFF-treated cortical neurons appeared predominantly as granular-like aggregates (∼45% hWT; ∼30% E83Q) or as a mixture of granular and filamentous-like aggregates (∼41% hWT, ∼35% E83Q) that accumulate in the neuronal cell bodies (Figs. 10A-B and S17). As for the filamentous-like aggregates (∼10% hWT; ∼10% E83Q), they were found both in the neurites and the neuronal cell bodies (Figs. 10A-B and S17).

More importantly, while no LB-like inclusions were ever observed in hippocampal neurons treated with the hWT PFF, approximately 5% of newly formed aggregates in cortiacal neurons were detected as LB-like inclusions (Figs. 10A-B and S17). Interestingly, the proportion of LB-inclusions was significantly higher in cortical neurons treated with E83Q PFF (∼15%). Moreover, at D21, the majority of the LB-like inclusions in hWT PFF- (∼85%) and E83Q PFF- (∼70%) treated neurons exhibited a dense core (Figs. 10C and S17). In addition, a significantly higher number of LB inclusions with a ring-like organization were detected in cortical neurons treated with E83Q (∼30%) than with hWT (∼15%) PFFs (Figs. 10 and S17).

All types of seeded-aggregates formed in hWT and hE83Q PFF-treated cortical neurons were positively stained by the ubiquitin antibody and the Amytracker dye (Figs. 10C and S17). However, p62 antibody detected all of the seeded-aggregates but not the granular-like aggregates formed in cortical neurons. This is in line with our findings in the hippocampal neurons (Figs. 9C, S14Ag-i, Dg-i, S14Dg-i and S14Fg-i), suggesting that some of the pS129-positive species formed by the human PFFs (WT or E83Q) may represent non-fibrillar aggregated forms of aSyn. Furthermore, the LB-like inclusions formed in the cortical neurons upon hWT PFF treatment were also positive for all the LB markers. This confirms that these inclusions (Figs. 10 and S17) shared the same properties as those formed in the E83Q PFF-treated cortical (Figs. 10 and S17) or hippocampal neurons (Figs. 9 and S13-S16).

Interestingly, we observed distinct patterns of seeding activity in cortical vs. hippocampal neurons. Firstly, the extent of seeding and LB-like inclusions formation after the addition of the E83Q PFF was significantly lower in the cortical neurons (∼15%) (Fig. 10B) than in the hippocampal neurons (∼30%) (Fig. 9B). Such differences might be due to the reduced aSyn levels and seeding activity of the aSyn PFFs in cortical neurons (*46–48*) (Fig. S12). Secondly, although hE83Q PFFs induced the formation of predominantly ring-like inclusions (∼70%, Fig. 9) in hippocampal neurons, the dense-LB-like inclusions were the dominant in cortical neurons (∼70%, Fig. 10C). Finally, both WT and E83Q PFFs induced the formation of nuclear pS129-positive aSyn aggregates in cortical with more nuclear aggregates detected in E83Q-PFF cortical neurons (∼8%) compared to WT-PFF treated cortical neurons (∼2%) (Figs. 10A-B and S17). These nuclear aggregates were not observed in the PFF-treated hippocampal neurons (Figs. 9 and S13-16).

Finally, we observed that compared to the control primary cortical cultures treated with PBS, the total number of cells (DAPI ^+^ cells) was significantly reduced by ∼ 40% in those treated with hWT or E83Q PFF (Fig. 10E). At D14, both the neuronal (NeuN ^+^ cells) and the glial (NeuN^−^ cells) populations were dramatically affected with ∼40% of cellular loss in the primary cortical cultures treated with hWT PFF (Fig. 10F-G). Neuronal cells loss was even significantly higher (∼60%) in the E83Q PFF-treated primary cortical neurons (Fig. 10F). No further increase in cell death was quantified between D14 and D21 in the glial or neuronal cell population (Fig. 10E-G). Furthermore, although the cortical neurons bearing LB-like inclusions and especially those with a ring-like organization showed, by confocal imaging, signs of advanced cell death (i.e., cell bodies swelling and cytoplasm vacuolization; Fig. S17Af, Bf, Cb-c, f, Db-c, Eb-d), barely any DNA fragmentation was identified in these neurons by the TUNEL cell death assay (TUNEL ^+^ NeuN ^+^ and pS129 ^+^ neurons, Fig. S18). This suggests that the neurodegeneration associated with the LB formation and/or maturation in cortical neurons might involve alternative non-apoptotic cell death processes.

Altogether, these findings suggest that the differences in aSyn levels and cellular environment are key determinants of LB formation and maturation.

## Discussion

### The E83Q mutation dramatically accelerates aSyn aggregation and forms fibrils of distinct morphology, stability, and conformation

Given that the NAC domain is essential for the aggregation of aSyn, and that the E83Q mutation results in a reduction in charge within an already highly hydrophobic domain, it is not surprising that this mutant exhibited a ∼10-fold faster aggregation than WT aSyn (over the concentration range of 10-50 μM) (Fig. 3A-B). Furthermore, previous studies have shown that substitution of E83 by alanine enhances aSyn aggregation (*83*) and interferes with the ability of dopamine to inhibit aSyn aggregation *in vitro* (*84*). Interestingly, a comparison of the lag phase of E46K (∼40 h at 7.5 μM to ∼15 h at 100 μM) (*85, 86*) and H50Q aggregation kinetics (∼20 h at 50 μM to ∼25 h at 70 μM) (*87, 88*) with E83Q lag-phase (∼3 h at 10 μM to ∼0.9 h at 50 μM) suggests that the E83Q has the highest aggregation propensity among all the known aSyn mutants.

Our comparison of the biophysical properties of the WT and E83Q fibrils shows that that the E83Q mutant forms fibrils with different core structures and dynamic properties. This is reflected in our findings, where the E83Q protein formed shorter fibrils (average length: ∼180 nm) that exhibit increased stability, slower monomer release and distinct morphological, structural and ProK-resistance properties compared to WT fibrils. Furthermore, our ssNMR comparative studies provided residue-specific insights suggesting major differences in the molecular structures of fibrils formed by human E83Q and WT aSyn (Fig. 5A and B). These observations are consistent with recent cryo-EM studies demonstrating that other familial mutants (H50Q (*88*), A53T (*89*), and E46K (*86*)) form fibril structures/folds that differ from the various polymorphs observed for WT aSyn (*90, 91*). Interestingly, in all the cryo-EM structures of WT aSyn fibrils, the E83 residue is exposed on the fibril surface, except for polymorphs 2a and 2b (Fig. 1C). In most fibril structures, the E83 residue is not engaged inter-filament interactions, suggesting that the acidic group of glutamic acid interacts with the solvent. However, in the case of the E83Q mutation, removing the negative charge from E83 due to the glutamine substitution may disrupt the long-range hydrophilic interactions with the solvent along the fibril axis.

It was recently reported that hE46K fibrils possessed the cross-seeding ability that templated the fibrillation of hWT monomers to form fibrils that inherited the structural and pathological features of the hE46K fibril strain (*92*). Here, we also showed that E83Q PFFs have the ability to pass their morphological properties to hWT monomers, although they showed preferential seeding of E83Q monomers.

### The E83Q mutation promotes the accumulation and oligomerization of aSyn

Despite the fact that we did not see fibrils-like inclusions or LB-like pathology in mammalian cell lines, our data showed that overexpression of E83Q aSyn induced the formation of small, dot-like aggregates in a fraction of the transfected cells (Figs. 6A-E and S6). Using the cross-linking approach, we established that the formation of these puncta structures correlates with the accumulation of multimeric/oligomeric aSyn species in the soluble fractions (Fig. 6F-G) and increased toxicity (Figs. 6H and S6). These findings demonstrate that the E83Q mutation promotes the formation of aSyn oligomers. However, the fact that E83Q mutant is expressed at higher levels than WT aSyn makes it difficult to determine if the increased oligomerization is simply due to the differences in expression levels or the inherent aggregation properties of the E83 mutant.

As observed with many of the aSyn PD-linked mutations (E46K, H50Q, and A53T) (*49, 93–99*), the overexpression of E83Q in mammalian cell lines (HEK, HeLa, or M17 neuroblastoma cells) was not sufficient to induce the formation of pS129-positive aggregates or inclusions that share the morphological and biochemical features of bona fide LBs (Figs. 6A, D and S6). The lack of a simple correlation between the *in vitro* aggregation propensities of aSyn PD-linked mutations (e.g., E83Q aSyn; Fig. 3) and the extent of LB pathology formation in the brain or inclusion formation in cells and neurons suggest that additional cellular factors or stressors play important roles in regulating the intrinsic aggregation propensity of aSyn in cells and LB pathology in neurons. These findings also suggest that the different synucleinopathy-related mutations may exert their actions *via* distinct mechanisms.

### The E83Q-mutant aSyn PFFs induce the formation of LB-like inclusions in primary neurons

Several studies have consistently shown that WT human aSyn PFFs exhibit significantly reduced seeding activity than WT mouse aSyn PFFs in primary neurons. Previous studies have also demonstrated the existence of a species barrier that renders human aSyn fibrils less efficient at seeding endogenous aSyn in primary mouse neurons (*53, 54*) and rodents (*53*). This explains why the vast majority of neuronal seeding models are based on the use of mouse aSyn PFFs (Figs. S19-20). It is noteworthy that the primary structure of mouse and human aSyn sequences differ by seven amino acids. Herein, we showed that in both hippocampal and cortical neurons, the E83Q human aSyn PFFs not only exhibit a much higher seeding activity than WT human aSyn PFFs at D14 and D21 (Fig. 8B-C) but also form even higher levels of pS129 pathologies than mouse aSyn PFFs at D21 (Figs. S12D and S20). In our hands, the differences in seeding are not due to differences in the internalization pathway or processing of the fibrils (Figure S11).

Although human aSyn PFFs have been shown to induce the formation of filamentous aggregates in cells, there are no reports in the literature to date demonstrating the induction of LB-like inclusions by human aSyn PFFs in primary neurons. Herein, we showed that in primary hippocampal neurons, hE83Q PFFs, but not hWT PFFs, induce the *de novo* formation of fibrils that can mature into pathological LB-like inclusions as observed in neurons treated with mouse PFFs. These observations suggest that the E83Q mutation induces the formation of fibrils with conformational properties that render them more efficient at recruiting and seeding the aggregation of endogenous mouse aSyn monomers, leading to the accelerated formation of LB-like inclusions in the hippocampal neurons. Conversely, in cortical neurons, the formation and maturation of LB-like inclusions were not limited to the neurons treated with hE83Q PFFs, but also occurred, albeit at a much lower level, in neurons treated with hWT PFFs. Consistent with previous findings, in cortical neurons, both endogenous aSyn and seeding activity levels are much lower than in hippocampal neurons. Therefore, our data suggest that intrinsic cellular properties, other than the endogenous level of aSyn or its misfolded properties, contribute to the cellular vulnerability to LB pathology formation and maturation, as also suggested in human synucleinopathies (*100, 101*). Altogether, our findings demonstrate that in the primary neuronal seeding model, human PFFs generated from E83Q aSyn could: 1) overcome the reported species barrier (*53, 54*); and 2) enhance the capacity of the human PFFs to trigger the *de novo* formation of pS129-positive fibrils capable of converting into LB-like inclusions

### The E83Q mutation induced the formation of LB-inclusions that resemble many of the features of some *bona fide* Lewy body pathologies

To further assess the effect of the E83Q mutation on the morphological diversity and biochemical composition of the LB-like inclusions, we assessed their immunoreactivity to well-known LB markers. In both cortical and hippocampal neurons, the LB-like inclusions formed upon hE83Q PFF treatment were positively stained for the classical markers used to define *bona fide* LBs in human pathology, including pS129 aSyn, p62, ubiquitin, and the amyloid tracer dye (Amytracker). In addition, other LB markers, such as cytoskeletal proteins (*102*) (e.g., MAP2 and NFL) and mitochondria (*71, 103*) (e.g., Tom 20), were also found to colocalize with the LB-like inclusions formed in hE83Q PFF-treated neurons. Strikingly, the distribution of these markers within the LB-like inclusions greatly resembled their distribution in LBs in the brain tissues of PD patients (*22, 69, 71, 72, 74, 78, 103-108*).

Interestingly, the addition of E83Q PFFs induced the formation and maturation of seeded-aggregates with distinct morphological and organizational features (Figs. 8-9 and S13-S16) in cortical and hippocampal neurons. Morphological classification based on pS129 staining revealed that at D21 the E83Q PFF-seeded neurons were populated either by filamentous- and/or granular-like inclusions (∼70% in hippocampal neurons; ∼85% in cortical neurons), or by LB-like inclusions (∼30% in hippocampal neurons; ∼15% in cortical neurons). Interestingly, in both cortical and hippocampal E83Q PFF-treated neurons, we were able to detect LB-like inclusions with the characteristic ring-like appearance of brainstem/nigral LBs (*25, 30-32, 69, 71, 109*) in addition to the LB-like inclusion with a dense core resembling cortical LBs (*25, 30, 105, 110-112*). Intriguingly, the relative distribution of the ring- and dense core-like inclusions depended on the type of neurons, with the ring-like inclusions mainly detected in hippocampal neurons (∼70%) and the dense core-like inclusions in cortical neurons (∼80%). In comparison, the LB-like inclusions observed in mouse PFF-treated neurons presented mostly with a dense core (∼80–90% at D14 and D21) (Figs. 9 and S13C-D), and only a few presented with a ring-like organization. This suggests that the remodeling of the newly seeded fibrils into LB-like inclusions of distinct morphologies is driven not only by the structure of the PFF seeds but also by neuronal-cell type-dependent properties. These findings are consistent with the immunohistochemical-based studies that revealed that the morphology of the LBs varies according to their location in human brain tissues from PD and synucleinopathies (*26, 69, 100, 101, 108, 113*). Finally, at D21, the level of LN-like pathology detected in E83Q PFFs-treated neurons was higher than that of the mouse PFF-treated neurons (Fig. 8A and S13F). This demonstrates that the use of PFFs generated from aSyn carrying the E83Q mutation in the neuronal seeding model allows the reproduction of some of the aSyn pathological diversity that may be relevant to the aSyn pathological heterogeneity observed in PD and synucleinopathies (*26*), including the formation of the halo-like LBs, the diffuse cortical LBs and the Lewy neurites-like aggregates in the neuritic extensions.

Recently, the organizational complexity of ring-like brainstem LBs was further characterized *via* microscopy (*71, 109*). Using aSyn antibodies specifically against pS129, full-length aSyn, or C-terminal truncated aSyn, these studies revealed a distribution of concentric layers of aSyn species in LBs, with pS129 detected mainly at the periphery. In contrast, the C-terminal truncated aSyn species accumulated in the center core of the inclusions. The NFL cytoskeletal proteins and mitochondria, on the other hand, accumulated at the outer shell (*71*). To determine to what extent the E83Q inclusions resemble the polymorphism of LB pathology in human brains, we examined the distribution of aSyn species and various known components of LBs using ICC (Figs. 9 and S13–S16, S21). The LB-like inclusions formed in the hE83Q or mWT PFF-treated neurons as dense core inclusions showed a homogenous and even distribution pattern of the pS129 aSyn, p62, ubiquitin, and the Amytracker signals across the inclusion as previously reported for the cortical LB (*105, 111, 112*). Conversely, in the LB-like inclusions with a ring-like appearance, pS129 aSyn (Figs. 9 and S13–S16) and the Amytracker signal are preferentially localized at the periphery of these inclusions (Fig. S21). In contrast, p62 and ubiquitin were detected at both the periphery and the core of the ring-like inclusions as NFL cytoskeletal proteins and mitochondria (Fig. S21). Such organization resembles the brainstem LB inclusions observed in human brain tissues from stage 3–5 PD patients (*25, 26, 31, 32, 69, 71, 109*). These findings further illustrate the power of using the neuronal seeding model as a platform to investigate the sequence (e.g., PD-linked mutations), molecular and cellular determinants of LB formation and to screen for modifiers of aSyn pathology formation and toxicity.

In this study, we investigated the effect of the first NAC-domain DLB-linked mutation on the structure of aSyn (monomers and fibrils) its aggregation (oligomerization and fibril formation) *in vitro* and in cells, and fibril seeding activity in primary neurons. Our results show that the E83Q mutation: 1) perturbs the conformational ensembles of free aSyn without impairing its ability to bind membranes *in vitro* and intracellularly; 2) dramatically accelerates the aggregation kinetics of aSyn *in vitro*; and 3) alters the dynamics and core structure of aSyn fibrils. The E83Q fibrils are shorter in length, more stable and exhibit a ProK digestion profile distinct from that of WT aSyn. In the cell-based overexpression models, the E83Q mutation did not affect the cellular distribution of aSyn, but it enhanced aSyn oligomerization and promoted the formation of pS129-negative puncta structures. Finally, studies in our neuronal seeding models showed that: 1) the E83Q mutation dramatically increases the seeding activity of human PFFs; and 2) the E83Q mutation can restore the capacity of PFFs to induce the formation of LB-like inclusions in mouse hippocampal neurons. Most notable is the fact that the E83Q fibrils induced the formation of aSyn-aggregate structures with a spectrum of morphologies similar to some of the Lewy pathologies observed in human PD brains, including the classical brainstem LB structures. In both cellular models, overexpression of E83Q or seeding with E83Q PFFs was associated with neurodegeneration and apoptosis. Altogether, our findings suggest that (E83Q) unlocks the pathogenicity of human aSyn fibrils and recapitulates some of its pathological diversity by promoting its oligomerization, increasing its toxicity and inducing the formation of more pathogenic fibrillar structures. Thereofore, exploring the effects of this mutation in various models of synucleinopathies could yield novel insights into the molecular mechanisms underpinning aSyn-induced neurodegeneration and aSyn pathology formation and spreading within the brain.

## Materials and Methods

### Recombinant overexpression and purification of human WT and E83Q aSyn

Recombinant overexpression and purification of human WT aSyn were performed as described previously (*114*). pT7-7 plasmids encoding human WT aSyn were used for transformation in BL21 (DE3) *E. coli* cells on an ampicillin agar plate. A single colony was transferred to 200 mL of Luria broth (LB) medium containing 100 μg/mL ampicillin (AppliChem, A0839) (small-scale culture) and incubated overnight at 180 rpm at 37 °C. The following day, the pre-culture was used to inoculate 6 L of LB medium containing 100 μg/mL ampicillin (large-scale culture) with the starting absorbance at 600nm of big culture between 0.05–0.1. Upon the absorbance at 600nm approaching 0.4 to 0.6, aSyn protein expression was induced by the addition of 1 mM 1-thio-β-d-galactopyranoside (AppliChem, A1008), and the cells were further incubated at 180 rpm at 37 °C for 4–5 h. Cells were harvested by centrifugation at 4000 rpm using JLA 8.1000 rotor (Beckman Coulter, Bear, CA) for 30 min at 5 °C. The harvested pellets were stored at −20 °C until use. Cell lysis was performed by dissolving the bacterial pellet in 100 mL of buffer A (40 mM Tris–HCl, pH 7.5) containing protease inhibitors [1 mM EDTA (Sigma-Aldrich, cat# 11873580001] and 1 mM PMSF (Applichem, A0999), followed by ultrasonication (VibraCell VCX130, Sonics, Newtown, CT) at the following parameters: 8 min; cycle: 30 sec ON, 30 sec OFF; amplitude 70%. After lysis, centrifugation at 12,000 rpm at 4 °C for 30 min was performed to collect the supernatant. This supernatant was collected in 50 mL Falcon® tubes and placed in boiling water (100 °C) ∼15 min. This solution was subjected to another round of centrifugation at 12,000 rpm at 4 °C for 30 min. The supernatant obtained at this step was filtered through 0.45-μm filters and injected into a sample loop connected to HiPrep Q Fast Flow 16/10 (Sigma-Aldrich, GE28-936-543). The supernatant was injected at 2 mL/min and eluted using buffer B (40 mM Tris-HCl, 1 M NaCl, pH 7.5) from gradient 0 to 70% at 3 mL/min. All fractions were analyzed by SDS-PAGE, and the fractions containing pure aSyn were pooled, flash-frozen, and stored at –20 °C until use. Fractions containing the purified aSyn were loaded on a reverse-phase HPLC C4 column (PROTO 300 C4 10 μm, Higgins Analytical; buffer A, 0.1% TFA in water; buffer B, 0.1% TFA in acetonitrile), and the protein was eluted using a gradient from 35 to 45% buffer B over 40 min (15 mL/min). The purity of the elution from HPLC was analyzed by UPLC and ESI-MS. Fractions containing highly purified aSyn were pooled, snap-frozen and lyophilised.

### Preparation of WT and E83Q aSyn monomers

The lyophilized aSyn of required quantity was resuspended in PBS buffer and followed by the filtration protocol as described before (*38*). Briefly, the lyophilized aSyn following resuspension was adjusted to pH ∼7.2–7.4 and filtered using 100 kDa spin filters. The filtrate containing monomers was estimated for the concentration by measuring the absorbance at 280 nm by a Nanodrop 2000 Spectrophotometer (ThermoFisher), and using an extinction coefficient of 5960 M−1 cm−1 predicted using the sequence (ProtParam, ExPASy). Monomeric aSyn (both WT and E83Q) was prepared as described above and followed any kind of further experiments.

### Preparation of WT and E83Q aSyn preformed fibrils seeds for cell studies

WT and E83Q aSyn fibrils were prepared by dissolving 4 mg of lyophilized recombinant aSyn in 600 μL of 1x PBS and the pH was adjusted to ∼7.2–7.4. The solution was filtered through 0.2 μm filters (MERCK, SLGP033RS), and the filtrate is transferred to black screw cap tubes. This tube was incubated in a shaking incubator at 37 °C and 1000 rpm for five days. After five days, the formation of fibrils was assessed by TEM and Coomassie staining as described previously (*38*). For the preparation of sonicated seeds, WT and E83Q aSyn fibrils were subjected to sonication for 20 sec, at 20 % amplitude 1 sec pulse on and 1 sec pulse off (Sonic Vibra Cell, Blanc Labo, Switzerland) only in mentioned experiments. The amount of monomers and oligomers release from sonicated fibrils was quantified by filtration following the published protocol (*38*). The structure of the fibrils was monitored by TEM and the amount of released monomers and oligomers after sonication by Coomassie staining analysis.

### ThT fluorescence based aggregation kinetics

For the setup of ThT aggregation kinetics, a 400 μL master-mix solution (3x 110 μL) containing five different final concentrations of WT or E83Q aSyn monomers (10, 20, 30, 40 and 50 μM) was prepared in 1X PBS (gibco), respectively, to which a final ThT concentration of 20 μM was added.

For the assessment of aggregation of co-presence of the mixture of WT and E83Q monomers, a 400 μL master-mix solution (3x 110 μL) encompassing a ratio of WT:E83Q monomers concentration (40:10, 25:25, 10:40 μM) was prepared in 1X PBS (gibco), respectively, to which a final ThT concentration of 20 μM was added.

For the cross-seeding aggregation experiments, 2 % WT or E83Q PFFs (w/v, relative to the monomer concentration) was added to the 400 μL master-mix solution containing the final concentration (50 μM) of WT or E83Q aSyn monomers in 1X PBS, respectively, to which a final ThT concentration of 20 μM was added.

For all the ThT based aggregation assays, a 110 uL of replicates from each master-mix solution was added to the three individual wells in a black 96-well optimal bottom plate (Costar), with each well pre-added with six SiLibeads® ceramic beads (Sigmund Lindner) with a diameter of 1.0–1.2 mm. The plate was sealed with Corning® microplate tape and transferred to a Fluostar Optima plate reader (BMG labtech). The parameters used were: continuous orbital shaking at 600 rpm at 37 °C, ThT fluorescence was monitored every 300 sec using excitation at 450 ± 10 nm and emission at 480 ± 10 nm. At the endpoint of the assays, the samples were analyzed by electron microscopy.

### Solubility assay

As described for the ThT aggregation kinetics experiments, a 800 μL master-mix solution (5x 150 μL) containing three different final concentrations of WT or E83Q aSyn (10, 30, and 50 μM) was prepared, respectively. Three individual samples for each concentration was set as replicates. A volume of 150 μL each from the master-mix solution was added to the 4 individual wells, respectively, onto a black 96-well optimal bottom plate (Costar), with each well pre-added with six SiLibeads® ceramic beads (Sigmund Lindner) with diameter of 1.0–1.2 mm. The plate was sealed with Corning® microplate tape and transferred to a Fluostar Optima plate-reader (BMG labtech). Aggregation was induced on the 96-well plate by using the parameter used for ThT aggregation kinetics such as continuous orbital shaking at 600 rpm at 37 °C. As shown in the Fig. 3D and E, at every timepoints (0, 3, 6, 18, 24, 30, 42, 48, 66, and 90 h) nearly 70 μL sample was taken from one well of the plate after pausing the plate reader. Out of 70 μL drawn, 25 μL used for the EM/AFM imaging analysis. 40 μL from the remaining was centrifuged at 100,000 *g* for 30 min at 4 °C. After carefully removing the supernatant, one part of it used for the UPLC analysis and another part for the SDS-PAGE analysis.

### Proteinase K digestion of WT and E83Q fibrils

The fibrils used for the proteinase K (ProK) digestion were prepared as above and not subjected to sonication. Briefly, 10 μg of fibrils was taken in 20 μL of 1X PBS, added with increasing concentrations (0.0, 0.5, 1.0, 1.5, and 2.0 μg/mL) of ProK and incubated for 37 °C for 30 min. Following incubation, the reaction was quenched by directly adding 5 μL of 5X Laemmli buffer to the reaction mixture and boiling the samples for 8 min at 95 °C. The digested fibrillar products were separated by SDS-PAGE using precast 12% Bis-Tris gels (Bio-Rad, Criterion XT) with MES running buffer (Bio-Rad). Bands were visualized using the Coomassie blue staining.

### Negative staining: transmission electron microscopy

Electron microscopy analyses of protein samples were performed as described previously (*114*). Briefly, 5 μL protein samples were placed on glow discharged Formvar and carbon-coated 200 mesh-containing copper EM grids. After about a minute, the samples were carefully blotted using filter paper and air-dried for 30 sec. These grids were washed three times with water and followed by staining with 0.7% (w/v) uranyl formate solution. TEM images were acquired by Tecnai Spirit BioTWIN electron microscope. Widths of different fibril morphologies (single, twisted and stacked) and periodicity lengths of twisted fibrils were measured using the image processing package Fiji(*115*). The relative frequency of each fibril morphology was estimated by examining and counting randomly chosen all visible fibrils within the TEM images. The number of fibrils quantified for WT is 164 for WT and 176 for E83Q.

### Mass spectrometry analysis

Mass spectrometry (MS) analyses of proteins were performed by liquid chromatography-mass spectrometry (LC-MS) on the LTQ system (Thermo Scientific, San Jose, CA). Before analysis, proteins were desalted online by reversed-phase chromatography on a Poroshell 300SB C3 column (1.0 × 75 mm, 5 μm, Agilent Technologies, Santa Clara, CA, on the LTQ system). 10 uL protein samples, roughly at a concentration of 0.5–1 μg, were injected on the column at a flow rate of 300 μ L/min and were eluted from 5% to 95% of solvent B against solvent A, linear gradient. The solvent composition was Solvent A: 0.1% formic acid in ultra-pure water; solvent B: 0.1% formic acid in acetonitrile. MagTran software (Amgen Inc., Thousand Oaks, CA) was used for charge state deconvolution and MS analysis.

### Native nESI- ion mobility- MS experiments

For the monomer experiments, WT and E83Q aSyn were redissolved in 20 mM aqueous ammonium acetate (Sigma-Aldrich, St. Louis, Missouri, USA) solution at pH 6.8 with a final protein concentration of 20 µM. Native IM-MS experiments were performed on a Synapt G2 HD mass spectrometer (Waters Corporation, Wilmslow, UK) equipped with a direct infusion nESI source. In-house generated gold-coated borosilicate capillaries were used to introduce a few µL of sample into the instrument. The most important experimental parameters were: spray capillary 1.5-1.7 kV; source temperature 30 °C; sample cone 25 V; extraction cone 1 V; trap collision energy 5 V; transfer collision energy 0 V; trap bias 40 V; IM wave velocity 300 m/s; IM wave height 35 V. Gas pressures in the instrument were: source 2.6 mbar; trap cell 0.024 mbar; IM cell 3.0 mbar; transfer cell 0.025 mbar. For collision induced unfolding experiments (CIU) identical instrument setting were used, but the trap collision energy (the acceleration voltage into the trap cell) was increased in a stepwise manner in 5 V increments. Mass spectra were analysed and visualised using MassLynx V4.1 (Waters Corporation, Wilmslow, UK) and Origin 8 (OriginLab Corporation, Northampton, USA). Drift time profiles were generated by selecting the upper half of mass spectral peaks, i.e. the m/z range from half height to half height of each peak, to avoid overlap with adduct peaks. Drift times from a travelling wave ion mobility cell can be converted into collision cross section (CCS) values, the rotationally averaged projection area of the protein, by calibration. Denatured cytochrome C from equine heart, denatured ubiquitin from bovine erythrocytes and denatured myoglobin from equine skeletal muscle (all purchased from Sigma-Aldrich, St. Louis, Missouri, USA) were used to calibrate IM measurements of α-syn based on literature CCS values (*116*) at a final concentration of 10 µM in 50/49/1 H2O/acetonitrile/formic acid. Acetonitrile (≥99.8%) and formic acid (99%) were purchased from Thermo Fisher Scientific (Waltham, Massachusetts, USA).

### nESI-MS analysis of fibril samples

For the fibril experiments, WT and E83Q fibrils were separated from eventual remaining monomer using centrifugal filters with a cut-off of 100 kDa (MilliporeSigma, Burlington, Massachusetts, USA). Samples were incubated with 0.5 µg/mL proteinase K for 5 minutes at 37 °C. After incubation the samples were frozen at -80 °C and thawed immediately before the measurement. Control samples were measured directly, while an additional C18 ZipTip (MilliporeSigma, Burlington, Massachusetts, USA ) purification step was necessary for the digested fibril samples. Measurements were performed with identical experimental parameters as for the monomer samples. Mass spectra were analysed and visualised as described above and peptide masses were calculated and assigned to peptide fragments using PAWS (Protein Analysis Work Sheet, Genomic Solutions Ltd, Cambridgeshire, UK), with a mass tolerance of ± 250 ppm.

### Preparation of small unilamellar vesicles

The preparation of small unilamellar vesicles (SUVs) was carried out as previously described (*97*). Briefly, the negatively charged phospholipid 1-hexadecanoyl-2-(9Z-octadecenoyl)-sn-glycero-3-phospho-(1′-racemic glycerol), denoted 16:0/18:1 phosphatidylglycerol and hereafter referred to as POPG, was purchased from Avanti Polar Lipids, Inc. (Alabaster, AL). A stock of POPG at 10 mg/mL in chloroform was prepared in a brown glass vial and stored at –20 °C. The SUVs were prepared using the extrusion method. Briefly, 100 μL of POPG from the stock was added to 900 μL of chloroform in a new glass vial to have a solution of 1 mg/mL POPG. The solution mixture was dried gently using a nitrogen stream to form a thin film on the wall of the glass vial. Any remaining chloroform was removed by placing the vial under a chemical hood overnight with the glass vial left opened. The phospholipids were then resolubilized in 1X PBS to reach their final concentrations of1.2 mM and followed by water bath sonication for 30 min. The solution was then extruded through Avestin LiposoFast™ (Avestin Inc., Ottawa, ON) using a membrane pore size of 100 nm. The size and homogeneity of the resulting vesicles were assessed by dynamic light scattering measurements (Stunner).

For another liposome preparation, 5 mg of 1,2-fioleoyl-sn-glycero-3-phosphoethanolamine (DOPE):1,2-dioleoyl-sn-glycero-3-phospho-L-serine (DOPS):1,2-dioleoyl-sn-glycero-3-phosphocholine (DOPC) 5:3:2 w/w (Avanti Polar Lipids) were resuspended in 0.8 mL methanol:chloroform 1:1, evaporated under a nitrogen stream, lyophilized o/n and resuspended in 0.52 mL of HEPES buffer (50 mM HEPES, 100 mM NaCl, pH 7.4, 0.02% NaN_3_). The resulting turbid sample was sonicated until transparency (15 minutes, 10’’ on, 20’’ off).

### Far-ultraviolet circular dichroism (CD) spectroscopy

Protein samples (10 μM) in 1X PBS loaded onto a 1-mm path length quartz cuvette were analyzed alone or mixed with POPG vesicles using the following protein:lipid molar ratios: 1:0.5, 1:1, 1:10, 1:25, 1:50, and 1:100. CD spectra were recorded at room temperature (RT) using a Chirascan spectropolarimeter (Applied Photophysics) with the following parameters: temperature 20 °C; wavelength range 198–250 nm; data pitch 0.2 nm; bandwidth 1 nm; scanning speed 50 nm/min; digital integration time 2 sec. The final CD spectra was a binomial approximation on an average of three repeat measurements. For the measurement of aSyn in the presence of another liposome mixture [DOPE:DOPS:DOPC (5:3:2)], individual WT and E83Q aSyn-to-liposome ratios were pipetted (ratio: 1:10, 1:20, 1:40, 1:100 and 1:200) at a protein concentration of 50 µM and the respective liposome concentrations from a 12.5 mM liposome stock in HEPES buffer (50 mM HEPES, 100 mM NaCl, pH 7.4, 0.02% NaN_3_) to a total volume of 12 µL. For CD measurement the sample was diluted in 48 µL deionized water and transferred to a 0.02 cm pathlength FireflySci cuvette for a final protein concentration of 10 µM. CD data were collected from 190 to 260 nm by using a Chirascan-plus qCD spectrometer (Applied Photophysics, Randalls Rd, Leatherhead, UK) at 20 °C, 1.5 time-per-point (s) in 1 nm steps. The datasets were averaged from three repeats. All spectra were baseline corrected against buffer in deionized water and smoothened (window size: 8).

### NMR spectroscopy

Isotopically labeled WT and E83Q aSyn were purified following the same purification protocol as described above in the section “ Recombinant overexpression and purification of human WT and E83Q aSyn” except the usage of the minimal medium instead of LB medium. Briefly, *E*-*coli* BL21(DE3) cells expressing the aSyn E83Q and WT were first grown in LB medium and then transferred to minimal medium containing ^13^C-labeled glucose and ^15^N-labeled ammonium chloride for production of ^13^C ^15^N-labeled E83Q and WT protein before induction. After protein expression, bacteria were lysed by sonication, and the lysate was subjected to 100 °C boiling for ∼15 min and further followed by the next steps of purification protocol such as anion-exchange chromatography, reversed-phase HPLC, and lyophilization. The purity of the lyophilized ^13^C ^15^N-labeled E83Q and WT protein was analyzed by UPLC and ESI-MS.

NMR experiments were measured on a Bruker 800 MHz spectrometer equipped with a 5 mm triple-resonance, pulsed-field z-gradient cryoprobe using two-dimensional ^1^H,^15^N SOFAST heteronuclear multiple quantum coherence (HMQC)^2^ and ^1^H,^13^C heteronuclear single quantum coherence (HSQC)^3^ pulse sequences for monomer characterization at 15 °C.

All experiments were performed in HEPES buffer (50 mM HEPES, 100 mM NaCl, pH 7.4, 0.02% NaN3) with 5 % (v/v) D2O. The sample concentration for natural abundance ^1^H,^13^C-HSQC experiments was 300 µM for both WT and E83Q aSyn. Spectra were processed with TopSpin 3.6.1 (Bruker) and analyzed using Sparky 3.115 (T. D. Goddard and D. G. Kneller, SPARKY 3, University of California, San Francisco). The combined ^1^H/^15^N chemical shift perturbation was calculated according to (((δ_H_)^2^+ (δ_N_/10)^2^)/2)^1/2^.

### Solid-state NMR spectroscopy

For the analysis of fibrils for ssNMR studies, ^13^C ^15^N-labeled E83Q and WT fibrils were prepared by dissolving ∼20 mg of lyophilized recombinant aSyn in 4 mL of 1x PBS (gibco), and the pH was adjusted to ∼7.2–7.4. The solutions were filtered through 0.2 μm filters (MERCK, SLGP033RS), and the filtrates were transferred to a 15 mL falcon tube. These tubes were incubated in a shaking incubator at 37 °C and 1000 rpm for five days. After six days, the formation of fibrils was assessed by TEM and Coomassie staining. NMR samples contained ∼6 mg of U-[^13^C,^15^N]-labeled protein packed into a 3.2 mm rotor. Solid-state NMR experiments were recorded on a 950 MHz Bruker Avance III HD standard-bore spectrometer equipped with a 3.2 mm (^1^H, ^13^C, ^15^N) Efree triple-resonance probe (Bruker Biospin). 13C–13C dipolar-assisted rotational resonance experiments (*117, 118*) with a mixing time of 50 ms and hNCA experiments (*119, 120*) were recorded at 265 K for WT and E83Q aSyn fibrils. The following 90° pulse widths were used: 2 – 3.75 μs for 1H, and 4.5 μs for 13C. 1H decoupling strengths were 80–100 kHz. Measurements were performed with a magic angle spinning frequency of 17 kHz. Spectra were processed using Topspin (Bruker) and analyzed with CcpNmr-Analysis (*121*). 13C–13C correlation DARR and 2D hNCA experiments were processed with a sine bell shift of 2 and 4, respectively.

### Atomic force microscopy imaging

AFM was performed on freshly cleaved mica discs that are positively functionalized with 1% (3-aminopropyl)triethoxysilane (APTS) in an aqueous solution for 3 min at RT. For the fibril deposition onto the substrates, 20 µL aliquots each containing a 50-µM protein solution was loaded on the surface of mica discs at predefined time-points. Deposition lasted 3 min and was followed by a gentle drying by nitrogen flow. The mica discs were stored in a desiccator for one day prior to imaging to avoid prolonged exposure to atmospheric moisture. AFM imaging was performed at RT by a Park NX10 operating in true non-contact mode that was equipped with a super sharp tip (SSS-NCHR) Park System cantilever. The length and height quantifications were performed manually using XEI software developed by Park Systems Corp.

### Cell culture and transfections

HEK293 HeLa and M17 cells were cultured at 95% air and 5% CO_2_ in DMEM (for HEK293 and HeLa) or DMEM-F12 (1:1; for M17) supplemented with 10% fetal bovine serum (Gibco) and penicillin-streptomycin (Thermo Fisher). According to the manufacturer’s protocol, transfections were performed with Effectene Transfection Reagent (Qiagen).

### Western blotting of HEK293, HeLa, and M17 cells

HEK293, HeLa, or M17 cells were lysed in RIPA buffer [150 mM sodium chloride, 50 mM Tris (pH 8.0), 1% NP-40, 0.5% deoxycholate, 0.1% SDS] supplemented with protease inhibitor cocktail, 1 mM PMSF, and phosphatase inhibitor cocktail 2 and 3 (Sigma-Aldrich). Cell lysates were cleared by centrifugation at 4 °C for 15 min at 13,000 rpm. The pellet (insoluble fraction) was resuspended in SDS/TBS supplemented with protease inhibitor cocktail, 1 mM PMSF, and phosphatase inhibitor cocktail 2 and 3 (Sigma-Aldrich) and sonicated using a fine probe (0.5-sec pulse at amplitude of 20%, 15 iterations). BCA protein assay was performed to quantify the protein concentration in the soluble and insoluble fractions before addition of 4X Laemmli buffer. Proteins from the soluble and the insoluble fractions were then separated on a 16.5% or a 18% SDS-polyacrylamide gel, transferred onto a nitrocellulose membrane (Thermo Fisher) with a semi-dry system (Bio-Rad), and immunostained.

### Subcellular fractionation of mammalian cell lines

Cells from a 10-cm plate were collected by scrapping in 500 μL fractionation buffer [HEPES (pH 7.4) 20 mM, KCI 10 mM, MgCl_2_ 2 mM, EDTA 1 mM, EGTA 1 mM, DTT 1 mM, protease inhibitor cocktail, and phosphatase inhibitor cocktails 2 and 3]. Cell suspensions were incubated for 15 min on ice before being mechanically lysed using a Dounce homogenizer (with type B pestle). 20 strokes (or until all cells are lysed as verified by microscopy) were performed. Cell extracts were then left on ice for 20 min before being centrifuged at 720 × *g* (3,000 rpm) for 5 min. The pellet contained nuclei, and the supernatant contained cytoplasm, membrane, and mitochondria fractions. The supernatant (A) was transferred into a fresh tube and kept on ice. The nuclear pellet was washed with 500 μL fresh fractionation buffer before being mechanically lysed using a Dounce homogenizer (with type B pestle) with a minimum of 20 strokes. Lysates were centrifuged at 720 × *g* (3,000 rpm) for 10 min. Supernatant (B) was discarded, and the pellet that contained nuclei was resuspended in TBS (pH 7.4, Tris(hydroxymethyl)aminomethane 50 mM and NaCl 150 mM) with 0.1% SDS. Sonication was then performed to shear genomic DNA and homogenize the lysate (3 sec on ice at a power setting of 60%). The supernatant (A) that contains the membrane fractions was centrifuged at 40,000 rpm (100,000 × *g*) for 1 h. The supernatant (C) that corresponds to the cytosolic fraction was collected in a fresh tube. The pellet was washed with 400 μL of fractionation buffer and resuspended using a Dounce homogenizer (with type B pestle) with a minimum of 20 strokes. Lysates were centrifuged for 45 min (100,000 × *g*), and the pellet was resuspended as membrane fraction in the same buffer used for the nuclei. Protein estimation was carried out using a BCA assay kit, and 30 μg of total protein was loaded onto 16% tricine gels, electrophoresed, and immunostained.

### Immunocytochemistry

Immortal cells or primary hippocampal neurons were washed twice with PBS, fixed in 4% PFA for 20 min at RT, and immunostained as previously described (*122*). The antibodies used are indicated in the corresponding legend of each figure. The source and dilution of each antibody can be found in Fig. S4. The cells plated on CS were then examined with a confocal laser scanning microscope (LSM 700, Carl Zeiss Microscopy) with a ×40 objective and analyzed using Zen software (RRID:SCR_013672). The cells plated in black, clear-bottom, 96-well plates were imaged using the IN Cell Analyzer 2200 (with a ×10 objective). For each independent experiment, two wells were acquired per tested condition; in each well, nine fields of view were imaged.

### Size-exclusion chromatography (SEC) and cross-linking assay

After the transfection of the HEK and HeLa cells for 72 h (i.e., with EV, WT aSyn, and E83Q aSyn), SEC and cross-linking experiments were conducted. Briefly, the media of both cell cultures was removed, and the cells were washed twice with PBS (pH 7.4). To detach the cells, 2 mL of trypsin was added to each plate, followed by incubation at 37 °C for 5–7 min. Subsequently, 8 mL of fresh media was added to quench the reaction, and the cells were transferred into a conical Falcon® tube and centrifuged at RT for 2 min at 300 × *g*. The supernatant was discarded, the pellet was resuspended in fresh media, and the cells were counted. Cells were spun down once more under the aforementioned conditions, the supernatant was discarded, and the cells were resuspended to a final concentration of approximately 4*10^6^ cells/180 μL of PBS supplemented with phosphatase inhibitor.

To evaluate the major aSyn species after overexpression in mammalian cells (HEK and HeLa), cell extracts were analyzed by SEC. Briefly, 160 μL of cell extracts combined with 1.6 μL of DMSO was injected onto a Superdex 200 10/300 GL column and equilibrated with 1X PBS (pH 7.4). Proteins were eluted at 0.5 mL/min, and 500-μL fractions were collected using the ÄKTA FPLC System (GE Healthcare). The SEC fractions were analyzed by WB, and the SEC chromatograms were replotted with OriginPro software.

The native oligomeric state of E83Q was studied through cross-linking experiments, as follows. DSG was used as the chemical crosslinking reagent and was solubilized in DMSO to an initial concentration of 100 mM. The DSG was then added to the cells for a final concentration of 1 mM; as a control, the same volume of DMSO was added to the cells. After a 30-min incubation at 37 °C at 500 rpm, the reaction was quenched by the addition of 1 M Tris pH 7.6, for a final concentration of 50 mM. Following the cross-linking reaction, the HEK and HeLa cells were lysed by a sonication step (amp: Amp 40%; iterations: 2 × 15 sec), after which the lysed cells were centrifuged for 30 min at 20,000 × *g* at 4 °C. The supernatant was collected and the pellet was snap-frozen. The protein concentration was assessed by BCA assay kit prior to Western blotting of the soluble fraction. All WB analyses related to cross-linking experiments were conducted using NuPAGE 4–12% Bis-Tris gels (Invitrogen). To further assess the size distribution of the native aSyn species upon cross-linking, 160 μL of HEK and HeLa cells resultant from the cross-linking experiments (+1 mM DSG) were analyzed by SEC in similar conditions as those mentioned above.

### Primary neuronal culture

Primary hippocampal and cortical neurons were prepared from P0 pups of WT mice (C57BL/6JRccHsd, Harlan) or aSyn KO mice (C57BL/6J OlaHsd, Harlan) and cultured as previously described (*22, 46, 123*). All procedures were approved by the Swiss Federal Veterinary Office (authorization number VD 3392). Briefly, the cerebral hippocampi and the cortexes were isolated stereoscopically in Hank’s Balanced Salt Solution (HBSS) and digested by papain (20 U/mL, Sigma-Aldrich) for 30 min at 37 °C. After inhibiting the papain activity using a trypsin inhibitor (Sigma-Aldrich), the tissues were dissociated by mechanical trituration. The cells were finally resuspended in adhesion media (MEM, 10% horse serum, 30% glucose, L-glutamine, and penicillin/streptomycin; Life Technologies) and plated in 24- or 6-well plates previously treated with poly-L-lysine 0.1% w/v in water (Brunschwig, Switzerland) at a density of 300,000 cells/mL (for biochemistry analysis) or 250,000 cells/mL (for ICC/confocal microscopy analysis). After 3 h, the adhesion media was removed and replaced with Neurobasal medium (Life Technologies) containing B27 supplement (Life Technologies), L-glutamine, and penicillin/streptomycin (100 U/mL, Life Technologies). Neurons were plated directly in Neurobasal medium in the black, clear-bottom, 96-well plates at a concentration of 200,000 cells/mL.

### Primary culture treatment with mouse aSyn fibrils or PBS

As morphological (*124–126*), physiological (*127–129*), genetic (*130*), and proteomic (*131*) changes are observed during the different developmental stages of primary neurons in culture (days *in vitro*, DIV), we adapted the time-course of the PFF treatment in primary neurons to be able to analyze the spatio-temporal effects of the seeding mechanism uncoupled from the intrinsic changes related to neuronal development and maturation (see also (*22*)). PFFs were added to the neuronal cell culture media at different DIV, and all PFF-treated neurons were harvested at DIV 26 (Fig. S10), which ensures that the developmental stage of primary neurons in culture was similar at harvest time. For control neurons, PBS was added at DIV 5.

The day of the treatment, human WT or E83Q PFF or mouse WT PFF were thawed at RT and diluted to a final concentration of 70 nM using the Neurobasal media collected from wells containing the plated neurons. The leftover Neurobasal media in the wells was then aspirated and replaced by the media containing the diluted PFFs. Treated neurons were cultured for up to 21 days without any further media changes until the end of the treatment (*22*).

### Quantitative high-content analysis (HCA)

After aSyn PFF treatment, primary hippocampal or cortical neurons plated in black, clear-bottom, 96-well plates (BD) were washed twice with PBS, fixed in 4% PFA for 20 min at RT, and immunostained as described above. Images were acquired using the Nikon 10×/0.45 Plan Apo, CFI/60 of the IN Cell Analyzer 2200 (GE Healthcare), a high-throughput imaging system equipped with a high-resolution 16-bit sCMOS camera (2048 × 2048 pixels), using a binning of 2 × 2. For each independent experiment, duplicated wells were acquired per condition, and nine fields of view were imaged for each well. Images were analyzed using CellProfiler 3.0.0 software (RRID:SCR_007358) to identify and quantify the level of LB-like inclusions (stained with pS129 antibody) formed in MAP2-positive neurons, the number of neuronal cell bodies (co-stained with MAP2 and DAPI), or the number of neurites (stained with MAP2).

### Preparation of soluble and insoluble fractions of primary hippocampal neurons

After aSyn PFF treatment, primary hippocampal neurons were lysed as previously described (*22–24*). Briefly, treated neurons were lysed in 1% Triton X-100/TBS (50 mM Tris, 150 mM NaCl, pH 7.5) supplemented with protease inhibitor cocktail, 1 mM PMSF, and phosphatase inhibitor cocktail 2 and 3 (Sigma-Aldrich). After sonication using a fine probe [(0.5-sec pulse at an amplitude of 20%, 10 iterations (Sonic Vibra Cell, Blanc Labo], cell lysates were incubated on ice for 30 min and centrifuged at 100,000 × *g* for 30 min at 4 °C. The supernatant (soluble fraction) was collected, and the pellet was washed in 1% Triton X-100/TBS, sonicated as described above, and centrifuged for another 30 min at 100,000 × *g*. The supernatant was discarded, whereas the pellet (insoluble fraction) was resuspended in 2% SDS/TBS supplemented with protease inhibitor cocktail, 1 mM PMSF, and phosphatase inhibitor cocktail 2 and 3 (Sigma-Aldrich), and sonicated using a fine probe (0.5-sec pulse at an amplitude of 20%, 15 iterations).

The BCA protein assay was performed to quantify the protein concentration in the total, soluble, and insoluble fractions before the addition of 4X Laemmli buffer (10% SDS, 50% glycerol, 0.05% bromophenol blue, 0.3 M Tris-HCl pH 6.8, and 20% *β*-mercaptoethanol). Proteins from the total, soluble, and insoluble fractions were then separated on 16% tricine gels, transferred onto a nitrocellulose membrane (Thermo Fisher) with a semi-dry system (Bio-Rad), and immunostained as previously described (*22*).

### Cell death quantification in mammalian cell lines and primary neurons

Mammalian cell lines (HEK293, HeLa, and M17) overexpressing WT or E83Q aSyn for up to 96 h or primary hippocampal neurons were plated in 96-well plates and treated with aSyn PFFs (70 nM or up to 500 nM). After 7, 14, and 21 days of treatment, cell death was quantified using complementary cell death assays. Using a CytoTox 96® Non-Radioactive Cytotoxicity Assay (Promega, Switzerland), the lactic acid dehydrogenase (LDH) released into culture supernatants was measured according to the manufacturer’s instructions. After a 30-min coupled enzymatic reaction, which results in the conversion of a tetrazolium salt into a red formazan product, the amount of color formed (proportional to the number of damaged cells) was measured using an Infinite M200 Pro plate reader (Tecan) at a wavelength of 490 nm.

In primary cortical neurons, cell death was quantified by the total count of cells and by the Terminal dUTP nick end-labeling (TUNEL) cell death assay. PFFs- or PBS- treated neurons were fixed at the indicated times in 4% PFA for 15 min at RT. Cells were permeabilized in a solution composed of 0.1% Triton X-100 in 0.1% citrate buffer, pH 6.0, and then washed in PBS buffer before incubation with terminal deoxynucleotide transferase (In Situ Cell Death Detection kit; Roche, Switzerland) for 1 h at 37 °C in a solution containing TMR red dUTP. The neurons were stained using an antibody against the NeuN protein, a nuclear neuronal marker. The neurons bearing seeded aggregates were identified using pS129 antibody (MJFR13). The nucleus was counterstained with DAPI at 1/5000 (Sigma-Aldrich, Switzerland). The cells were washed five times in PBS before mounting in polyvinyl alcohol (PVA) mounting medium with the anti-fading DABCO reagent (Sigma-Aldrich, Switzerland). The cells plated on CS were then examined with a microscope (EVOS, Life Technology) with a 10 × objective and analyzed using ImageJ (US National Institutes of Health, Bethesda, MD, USA). A minimum of 4000 cells was counting for each condition and for each independent experiment.

### Western blotting quantification

The level of total aSyn (15 kDa, 12 kDa, or HMW) or pS129-aSyn were estimated by measuring the WB band intensity using Image J software (U.S. National Institutes of Health, Maryland, USA; RRID:SCR_001935) and normalized to the relative protein levels of actin; in the case of the cross-linking experiments, the level was normalized to the relative protein levels of GAPDH.

### Statistical analyses

Independent experiments were performed to verify the findings (minimum 3 independent experiments) with all attempts at replication successful. No data were excluded from the analysis. The statistical analyses were performed using Student’s t-test, one-way ANOVA test followed by a Tukey-Kramer or HSD post-hoc tests or two-way ANOVA followed by multiple pairwise comparisons with Sidak’s correction post-hoc test using GraphPad Prism 9.1.1 software (RRID:SCR_002798). The data were regarded as statistically significant at p<0.05.

## Supporting information

Kumar and Mahul et al Supplementary Info

## Acknowledgments

We thank the CIME facility (EPFL) for the use of their electron microscopy facility and PTPSP (EPFL) for their tremendous support for the use of their CD instrument. We thank Dr. Fabien Kuttler (PTCB, EPFL) for helping optimize the HCA pipeline. We also thank Dr. Rajasekhar Kolla, Somanath Jagannath and Galina Limorenko for their critical review of the first versions of this manuscript and valuable feedback. We are grateful to Dr. Sergey Nazarov and Ahmed Sadek for their help with the generation of Figure 1C. We also thank Lorène Aeschbach and Yllza Jasiqi for their help with the generation of plasmids and proteins and the primary neuronal culture related to this work. We would like to express our gratitude to Dr. Janna Hastings and Nathan Riguet for their help with the statistical analysis.

## Author contributions

Conceptualization: HAL

Experimental design: HAL, STK, ALMM

Methodology: STK, ALMM, RNH, PM, RM, GR, AIO, IR, SD

Investigation: STK, ALMM, RNH, PM, RM, AIO, MZ, FS, HAL

Visualization: STK, ALMM, RNH, PM, RM, SD, AIO

Supervision: HAL

Writing—original draft: HAL, STK, ALMM, PM, RNH, RM

Writing—review & editing: HAL, STK, ALMM, RNH, PM, RM, SD, AIO, MZ, FS

## Data and materials availability

All data are available in the main text or supplementary materials.

## Supplementary Materials

Supplementary file is enclosed.

