## Supplementary material for "A novel mutation (E83Q) unlocks the pathogenicity of human alpha-synuclein fibrils and recapitulates its pathological diversity": Kumar and Mahul et al Supplementary Info

**This PDF file includes:**

Figs. S1 to S21
References (1 to 9)

**Figure S1**

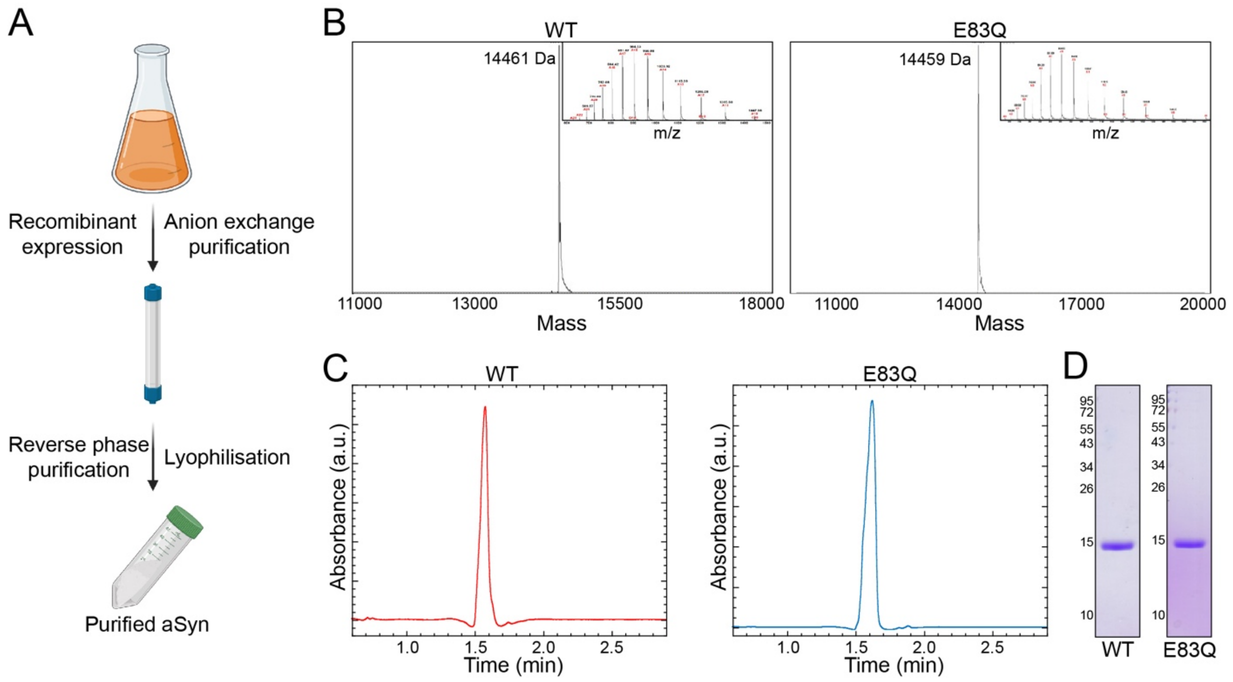


**Fig. S1. Purification strategy of aSyn proteins and their purity analysis**

**(A)** A scheme illustrating the recombinant expression and purification of both WT and E83Q aSyn used in this study. **(B)** ESI-MS spectra of WT (left, observed=14,461 Da; theoretical=14,459 Da) and E83Q (right, observed=14,459 Da; theoretical=14,458 Da) aSyn following purification. **(C-D)** Reverse phase UPLC spectra and SDS-PAGE analysis of WT (left) and E83Q (right) aSyn.

**Figure S2**

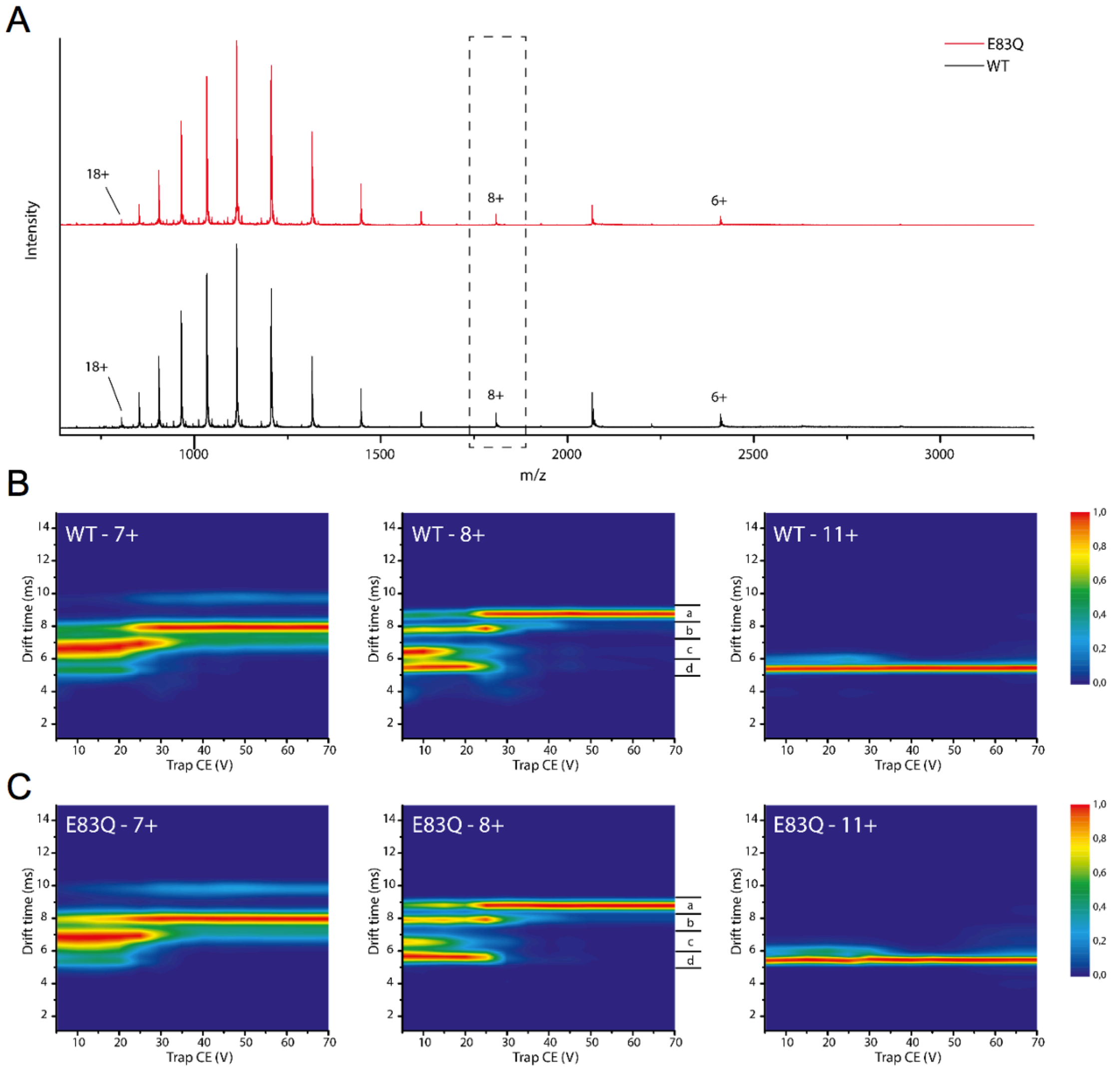


**Fig. S2. nESI-MS experiments of WT and E83Q aSyn. (A)** Mass spectra of WT (black) and E83Q (red) aSyn monomers. **(B-C)** CIU plots of the 7+, 8+, and 11+ charge state of WT (B) and E83Q **(C)** aSyn monomers, respectively. During the ESI process, a protein picks up a number of charges that are directly related to its solvent accessible surface area (SASA), and therefore to its conformation. Low charge states represent more compact conformations and high charge states indicate more extended protein states. Natively unfolded proteins, such as aSyn, display a broad range of charge states. Fig. S2A shows the native mass spectrum of aSyn WT and the E83Q mutant, and in both cases a multimodal intensity distribution ranging from 6+ to the 18+ charge state can be seen. No significant charge state intensity differences can be detected between the two proteins. To investigate if certain conformations are stabilized or destabilized by the mutation, collision induced unfolding (CIU) experiments are performed, which determine the amount of collisional activation (trap CE/voltage) leading to stepwise, gas-phase unfolding. CIU plots for the 7+, 8+, and 11+ charge state are shown in Fig. S2B and S2C, for the WT and E83Q mutant, respectively. For the 7+ charge state the most extended conformation is already present at lower voltages for the E83Q mutant compared to the WT, indicating reduced stability of the conformational family with a drift time around 6.5 ms for the mutant. For the 8+ charge state the two most extended conformational families, a and b, show similar patterns while the 'd' conformational family appears slightly more stable for the E83Q mutant compared to the WT as it is detected until a voltage of 25V and 20V is applied, respectively. The second most compact conformation is, as mentioned before, almost absent for the E83Q mutant. For the 11+ charge state no conformational changes are detected when increasing the energy, indicating that Coulombic forces here become the main determinant of the protein structure.

**Figure S3**

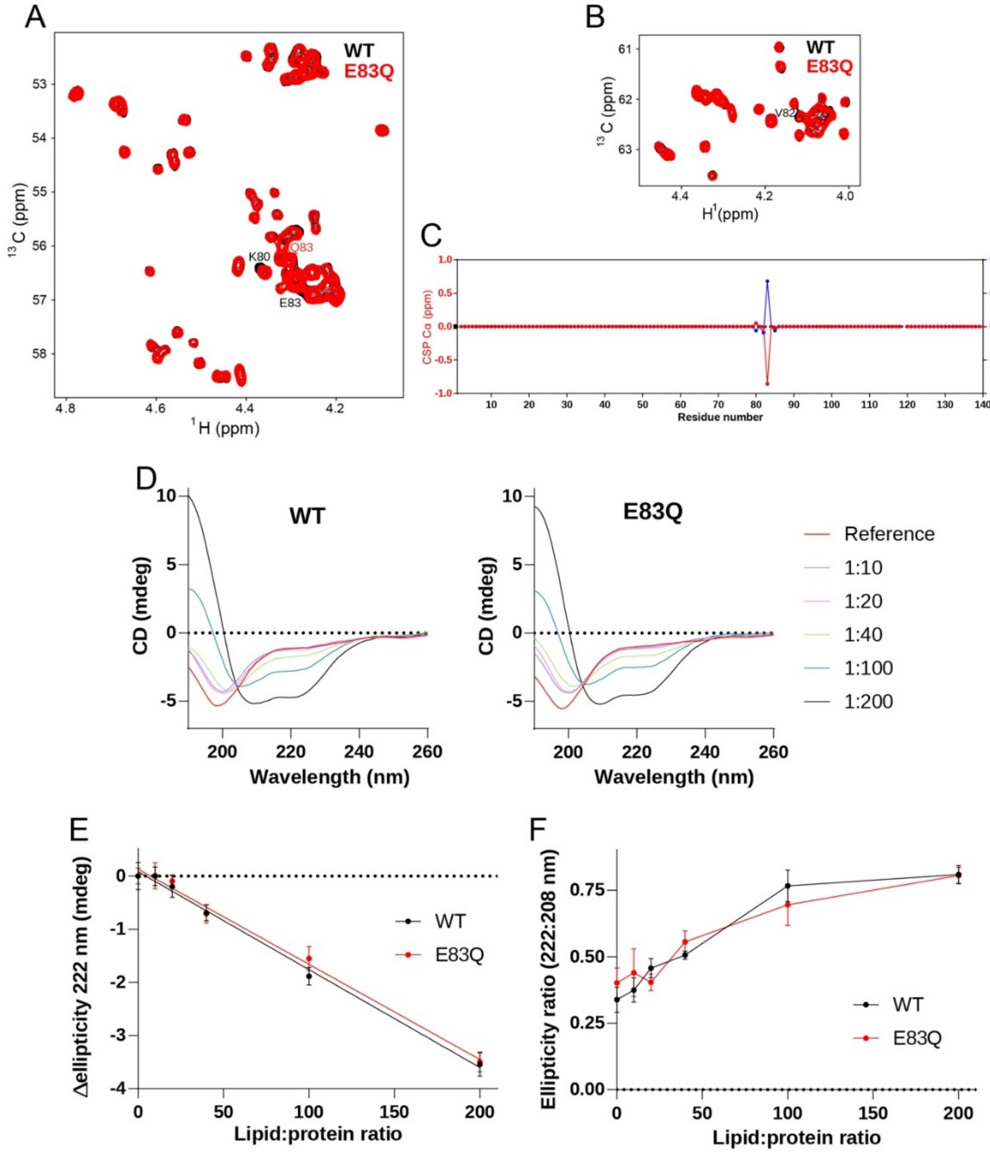


**Fig. S3. Secondary structure analysis of WT and E83Q only or in the presence of lipid vesicles. (A-B)** Selected regions of the $^{1}$H/$^{13}$C HSQC of WT (black) and E83Q (red) αSyn. The affected residues are labeled. E83 disappearing is not visible because of peak overlapping. **(C)** Cα (red) and Hα (blue) chemical shift perturbations between WT and E83Q αSyn based on the $^{1}$H/$^{13}$C HSQC spectra. **(D)** CD experiments of WT and E83Q αSyn at different protein:lipid ratios. Lipids used here are DOPE:DOPS:DOPC (5:3:2). **(E)** Change in ellipticity at 222 nm upon increasing concentrations of lipid for WT (black) and E83Q (red)indicating a similar increase of α-helical structure. **(F)** Change in the ellipticity ratio between 222 and 208 nm upon increasing concentrations of lipid for WT (black) and E83Q (red).

**Figure S4**

# A

| Primary Antibody | Catalog # | Company | Clone | RRID | Host | Concentration | WB dilution | ICC dilution | Epitope |
|---|---|---|---|---|---|---|---|---|---|
| anti-aSyn **total** | LASHUEL | - | LASH-EGT1-20 | - | Rabbit | Not provided | 1:1000 | 1:1000 | 1-20 |
| anti-aSyn **total** | LASHUEL | - | LASH-BL-A15110B | - | Mouse | 1.2 mg/ml | - | 1:500 | 34-45 |
| anti-aSyn **total** | 610787 | BD | SYN-1 | RRID:AB_398108 | Mouse | 0.25 mg/ml | 1:1000 | 1:1000 | 91-99 |
| anti-aSyn **total** | 4179 | Cell signalling | D37A6 | RRID:AB_1904156 | Rabbit | 1 mg/ml | 1:1000 | - | Near Glu105 |
| anti-aSyn **total** | 120101 | Biolegend | 4B12 | RRID:AB_389229 | Mouse | 0.5 mg/ml | 1:1000 | - | 103-108 |
| anti-aSyn **total** | Sc-12767 | Santa Cruz | Clone 211 | RRID:AB_628318 | Mouse | 0.2 mg/ml | 1:500 | - | 121-125 |
| anti-aSyn **total** | ab131508 | Abcam | - | RRID:AB_11155736 | Rabbit | 1 mg/ml | 1:1000 | 1:500 | 134-138 |

# B

| Primary Antibody | Catalog # | Company | Clone | RRID | Host | Concentration | WB dilution | ICC dilution | Epitope |
|---|---|---|---|---|---|---|---|---|---|
| anti-**pS129**-aSyn | ab168381 | Abcam | MJF-R13 | RRID:AB_2728613 | Rabbit | 4.229 mg/ml | 1:10 000 | 1:1000 | Not provided |
| anti-**pS129**-aSyn | 825701 | BioLegend | p-syn/81A | RRID:AB_2564891 | Mouse | 1.0 mg/ml | 1:1000 | 1:1000 | Peptide (residues 124-134) including phosphorylated Ser129 of human aSyn |
| anti-**pS129**-aSyn | 010-26481 | Wako | Biotin-pSyn #64 | RRID:AB_2537218 | Mouse | 1.0 mg/ml | 1:1000 | 1:1000 | residues 124-134 |

# C

| Primary Antibody | Catalog # | Company | Clone | RRID | Host | Concentration | WB dilution | ICC dilution | Epitope |
|---|---|---|---|---|---|---|---|---|---|
| anti-**actin** | ab6276 | Abcam | AC-15 | RRID:AB_2223210 | Mouse | 2.2 mg/ml | 1:5000 | Not tested | DDDIAALVIDNGSGK |
| anti-**MAP2** | ab92434 | Abcam | - | RRID:AB_2138147 | Chicken | Not provided | Not tested | 1:2000 | Recombinant full length protein |
| anti-**p62** | H00008878 | Abnova | 2C11 | RRID:AB_437085 | Mouse | 1 mg/ml | 1:1000 | 1:500 | Raised against a full length recombinant SQSTM1 |
| anti-**ubiquitin** | Sc-8017 | Santa-Cruz | P4D1 | RRID:AB_628423 | Mouse | 0.2 mg/ml | 1:500 | 1:500 | 1-76 |
| anti-**Tom 20** | sc-17764 | Santa-Cruz | F-10 | RRID:AB_628381 | Mouse | 0.2 mg/ml | 1:500 | 1:200 | Raised against amino acids 1-145 |
| anti-**NFL** | 837801 | Biolegend | SMI 311 | RRID:AB_2565383 | Mouse | 1mg/ml | - | 1:500 | Not provided |

# D

| Secondary Antibody | Catalog # | Company | RRID | Concentration | WB dilution | ICC dilution |
|---|---|---|---|---|---|---|
| Goat anti-**mouse Alexa Fluor 680** | A21058 | Invitrogen | RRID:AB_2535724 | 2 mg/ml | 1:5000 | - |
| Goat anti-**rabbit Alexa Fluor 680** | A21109 | Invitrogen | RRID:AB_2535758 | 2 mg/ml | 1:5000 | - |
| Goat anti-**mouse Alexa Fluor 800** | 926-32210 | Li-Cor | RRID:AB_621842 | 1 mg/ml | 1:5000 | - |
| Goat anti-**rabbit Alexa Fluor 800** | 926-32211 | Li-Cor | RRID:AB_621843 | 1 mg/ml | 1:5000 | - |
| Donkey anti-**mouse Alexa Fluor 568** | A10037 | Invitrogen | RRID:AB_2534013 | 2 mg/ml | - | 1:800 |
| Donkey anti-**rabbit Alexa Fluor 568** | A10042 | Invitrogen | RRID:AB_2534017 | 2 mg/ml | - | 1:800 |
| Donkey anti-**rabbit Alexa Fluor 647** | A31573 | Invitrogen | RRID:AB_2536183 | 2 mg/ml | - | 1:800 |
| Donkey anti-**mouse Alexa Fluor 647** | A31571 | Invitrogen | RRID:AB_162542 | 2 mg/ml | - | 1:800 |
| Donkey anti-**chicken Alexa Fluor 488** | 703-545-155 | Jackson Immunoresearch | RRID:AB_2340375 | 1 mg/ml | - | 1:400 |
| Goat anti-**chicken Alexa Fluor 568** | A11041 | Invitrogen | RRID:AB_2534098 | 2 mg/ml | - | 1:500 |

# E

| Dyes | Catalog # | Company | RRID | Concentration | WB dilution | ICC dilution |
|---|---|---|---|---|---|---|
| Amytracker**680** | Amytracker™680 | Ebba Biotech | - | | - | 1:500 |

**Fig. S4. List of the antibodies used in this study**
**(A)** Antibodies used for the detection of total aSyn. **(B)** Antibodies used for the detection of aSyn phosphorylated on S129 residue. **(C)** Other antibodies used in the study. **(D)** Secondary antibodies or dyes used for immunoblotting or confocal imaging.

**Figure S5**

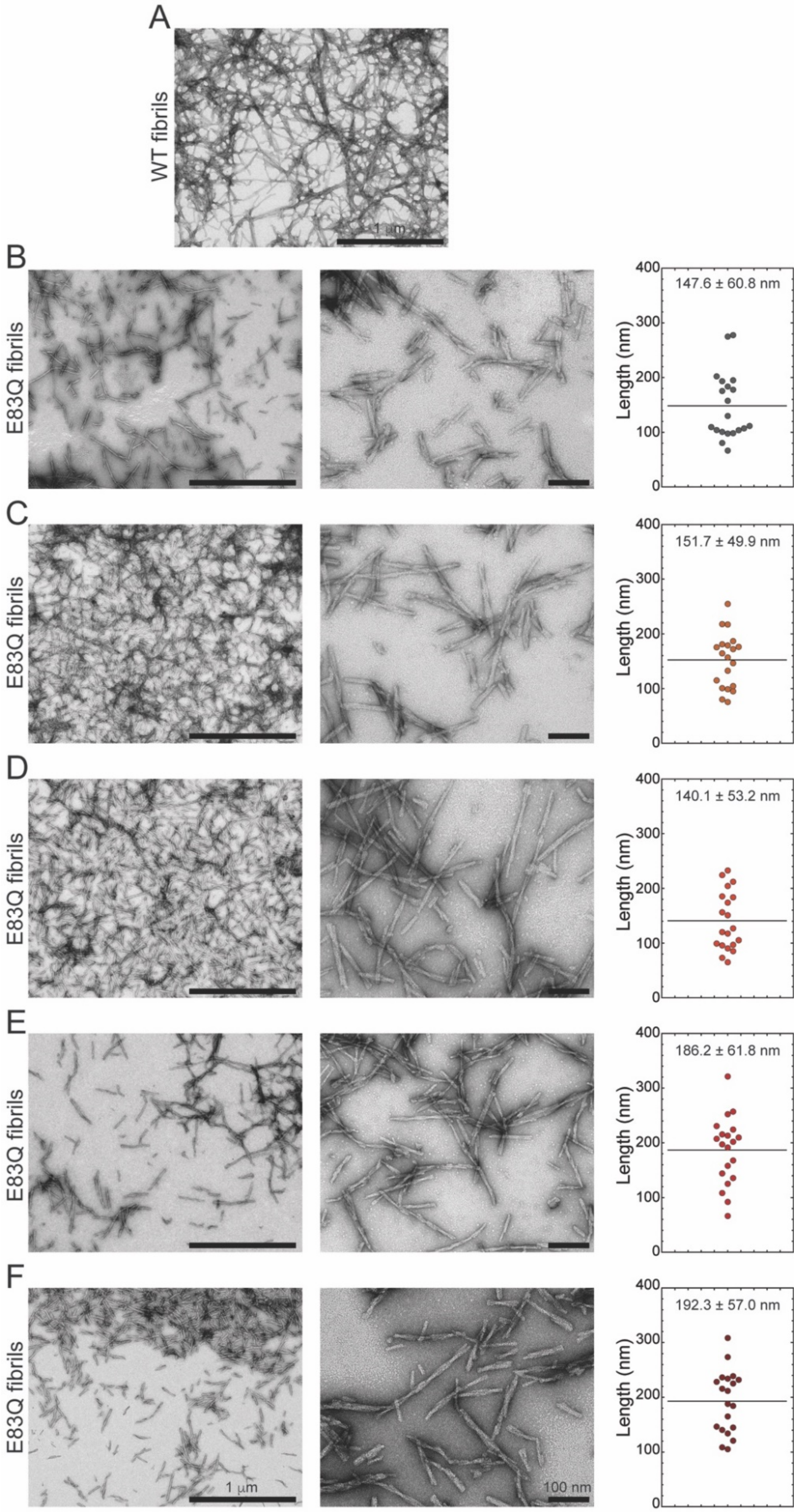

A
WT fibrils
1 μm
B
E83Q fibrils
147.6 ± 60.8 nm
Length (nm)
C
E83Q fibrils
151.7 ± 49.9 nm
Length (nm)
D
E83Q fibrils
140.1 ± 53.2 nm
Length (nm)
E
E83Q fibrils
186.2 ± 61.8 nm
Length (nm)
F
E83Q fibrils
192.3 ± 57.0 nm
Length (nm)
1 μm
100 nm
0
100
200
300
400

**Figure S5**

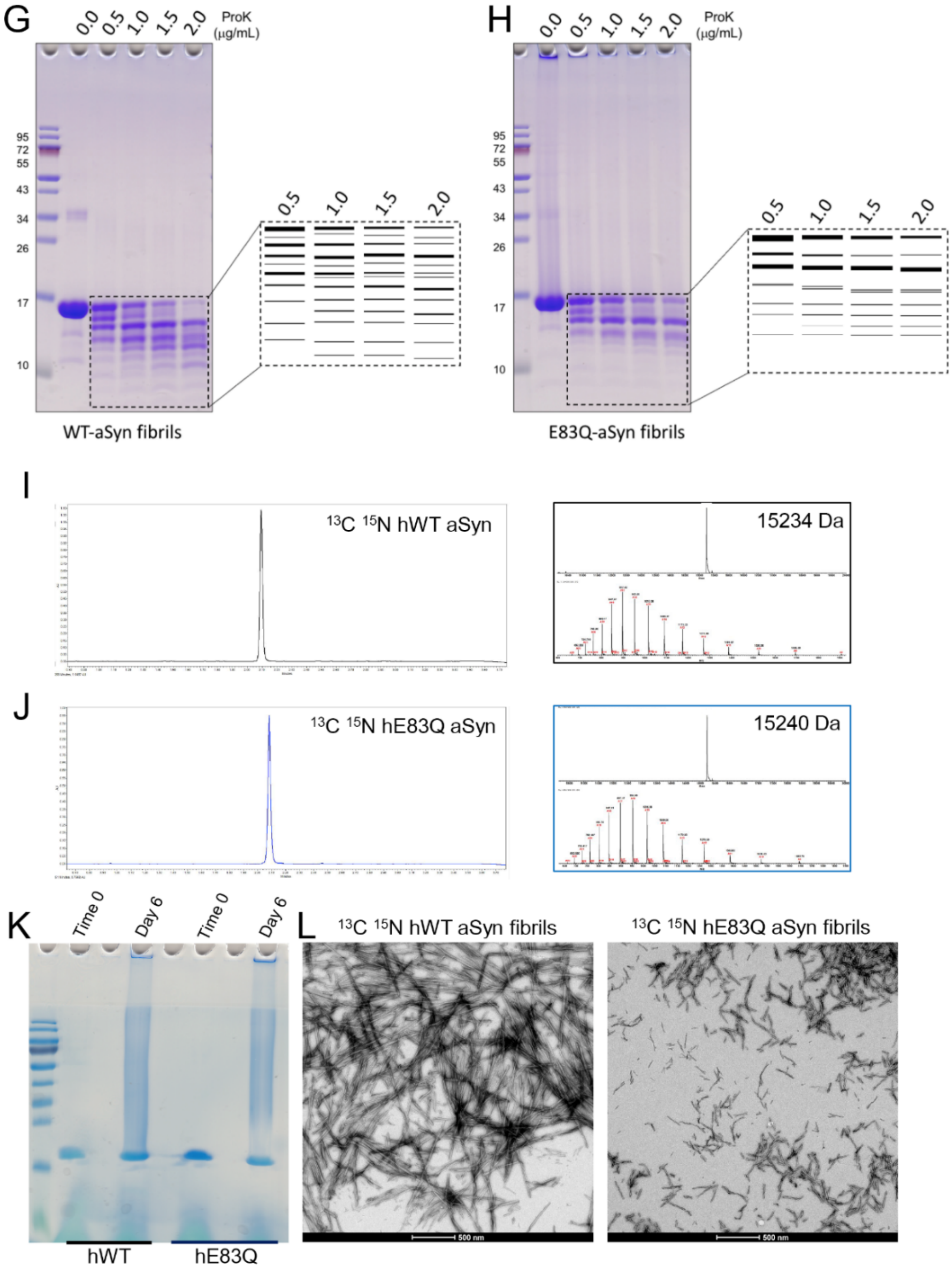


**Fig. S5. EM analysis and proteinase K analysis of WT and E83Q fibrils**. **(A)** EM image showing the long structures of WT fibrils. WT fibrils are longer than 1 µm in length consistently, proving harder for the length analysis. **(B-F)** EM images showing the highly reproducible shorter morphologies of E83Q fibrils, irrespective of varying aggregation conditions. Aggregation conditions include **B**) 20 µM E83Q in PBS (added 20 µM ThT), shaking at 600 rpm for 3 days in a 96 well plate, **C**) 50 µM E83Q in PBS (added 20 µM ThT), shaking at 600 rpm for 3 days in a 96 well plate, **D**) 50 µM E83Q in PBS, shaking at 600 rpm for 3 days in an Eppendorf tube, **E**) ~6.6 mg/mL of E83Q in PBS, shaking at 1000 rpm for 5 days in an Eppendorf tube, and **F**) ~5 mg/mL of E83Q in PBS, shaking at 1000 rpm for 5 days in a 15 mL falcon tube. **(G-H)** SDS-PAGE analysis of ProK digestion of WT **(G)** and E83Q **(H)** fibrils and illustration of the digested band patterns. **(I-J)** Reverse phase UPLC spectra and ESI-MS spectra of $^{13}$C $^{15}$N-labeled WT **(I)** and $^{13}$C $^{15}$N-labeled E83Q (J) aSyn following purification. **(K)** SDS-PAGE analysis of $^{13}$C $^{15}$N-labeled WT and $^{13}$C $^{15}$N-labeled E83Q aSyn before and after fibrillation. **(L)** TEM analysis of $^{13}$C $^{15}$N-labeled WT and $^{13}$C $^{15}$N-labeled E83Q fibrils. Scale bars = 500 nm.

**Figure S6**

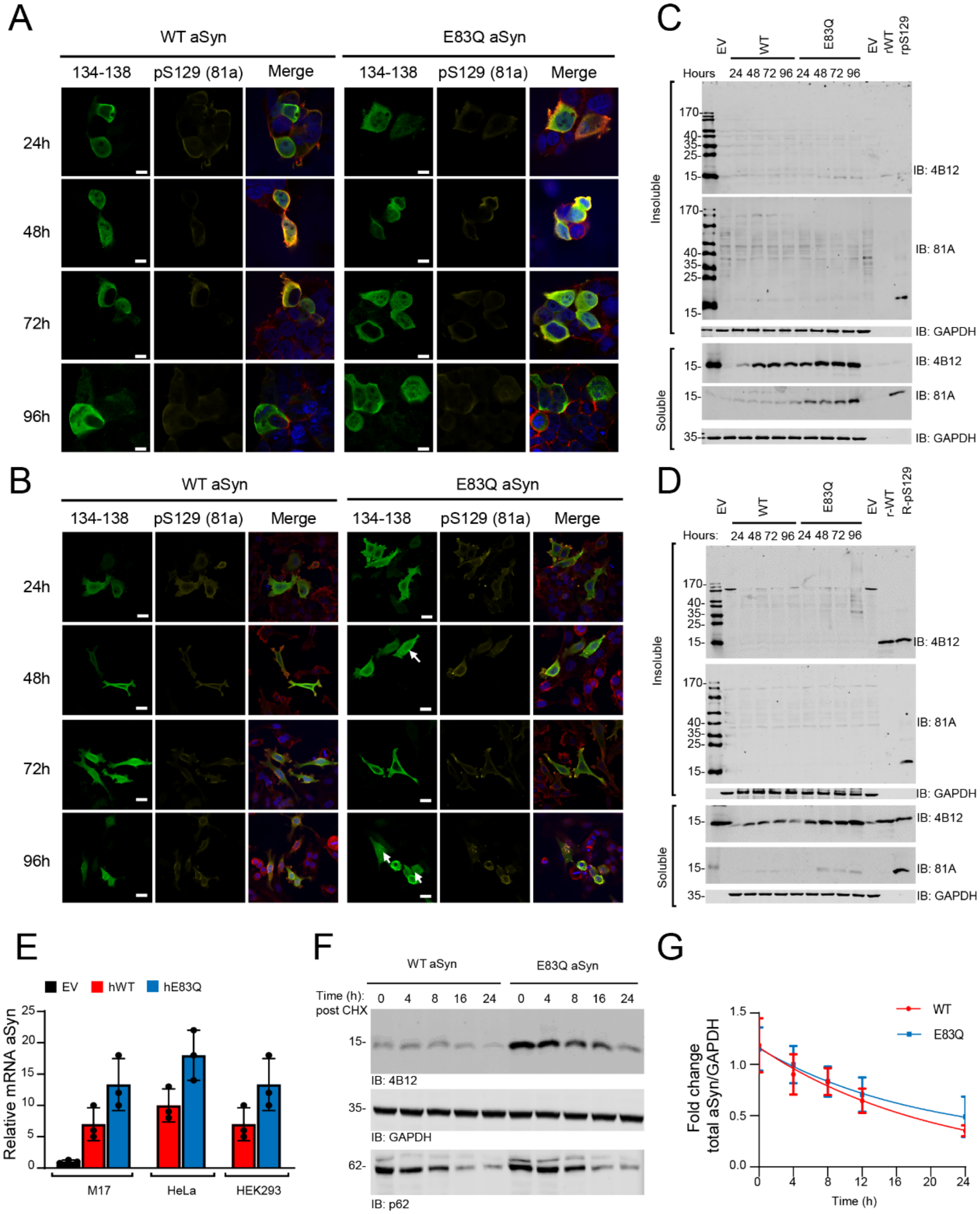
A
WT aSyn
E83Q aSyn
134-138
pS129 (81a)
Merge
24h
48h
72h
96h
B
WT aSyn
E83Q aSyn
134-138
pS129 (81a)
Merge
24h
48h
72h
96h
C
EV
WT
E83Q
EV
rWT
rpS129
Hours
24 48 72 96 24 48 72 96
Insoluble
Soluble
IB: 4B12
IB: 81A
IB: GAPDH
D
EV
WT
E83Q
EV
r-WT
R-pS129
Hours:
Insoluble
Soluble
IB: 4B12
IB: 81A
IB: GAPDH
E
EV
hWT
hE83Q
Relative mRNA aSyn
M17
HeLa
HEK293
F
WT aSyn
E83Q aSyn
Time (h): post CHX
0 4 8 16 24
IB: 4B12
IB: GAPDH
IB: p62
G
WT
E83Q
Fold change total aSyn/GAPDH
Time (h)

**Figure S6**

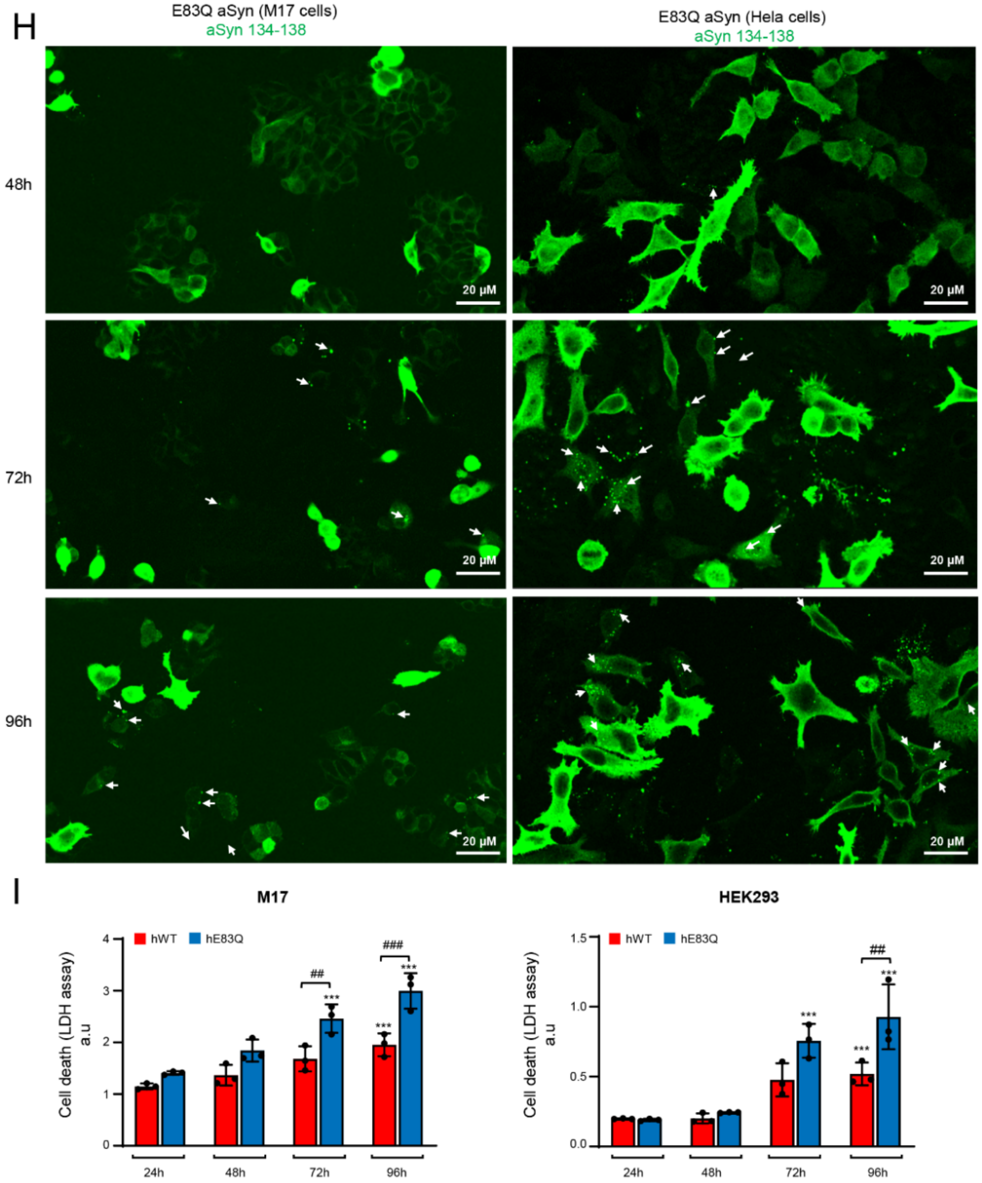


**Fig. S6. Overexpression of E83Q aSyn in immortal mammalian cells does not change the cellular properties compared to WT aSyn. (A-B)** Immunofluorescence (ICC) image of HEK293 cells (**A**) or HeLa cells (**B**) transfected with either WT or E83Q aSyn plasmids for indicated time. **(C-D)** WB of detergent soluble and insoluble fractions of lysed HEK293 cells **(C)** or HeLa cells **(D)** transfected with either WT or E83Q aSyn plasmids for the indicated time. **(E)** aSyn mRNA levels in the indicated cells. The graphs represent the mean +/- SD of a minimum of 3 independent experiments. No significant differences were measured. (**F**) WB of the detergent soluble fraction of lysed HEK293 cells transfected with either WT or E83Q aSyn plasmids for 48 hours at this time 100ug/ml of cycloheximide was added to cells for cycloheximide pulse-chase analysis and then cells were lysed at indicated time-point. (**G**) Fold change of aSyn quantified using densitometry after cycloheximide treatment for the indicated time estimated from the WB like in D. The graphs represent the mean +/- SD of a minimum of 3 independent experiments. No significant differences were measured. **(H)** Large field of view of HeLa cells overexpressing aSyn E83Q for 48h, 72h and 96h showing the formation overtime of dot-like species in the cytosol of these cells. **(I)** Cell death was measured by quantifying the levels of LHD released to the medium at indicated time and cell types. The graphs represent the mean +/- SD of a minimum of 3 independent experiments. $p<0.0005$=*** (ANOVA followed by Tukey HSD *post-hoc* test, 24h vs. other time-points). $p<0.005$=##, $p<0.0005$=### (ANOVA followed by Tukey HSD *post-hoc* test, WT vs. E83Q overexpressing cells).

**Figure S7**

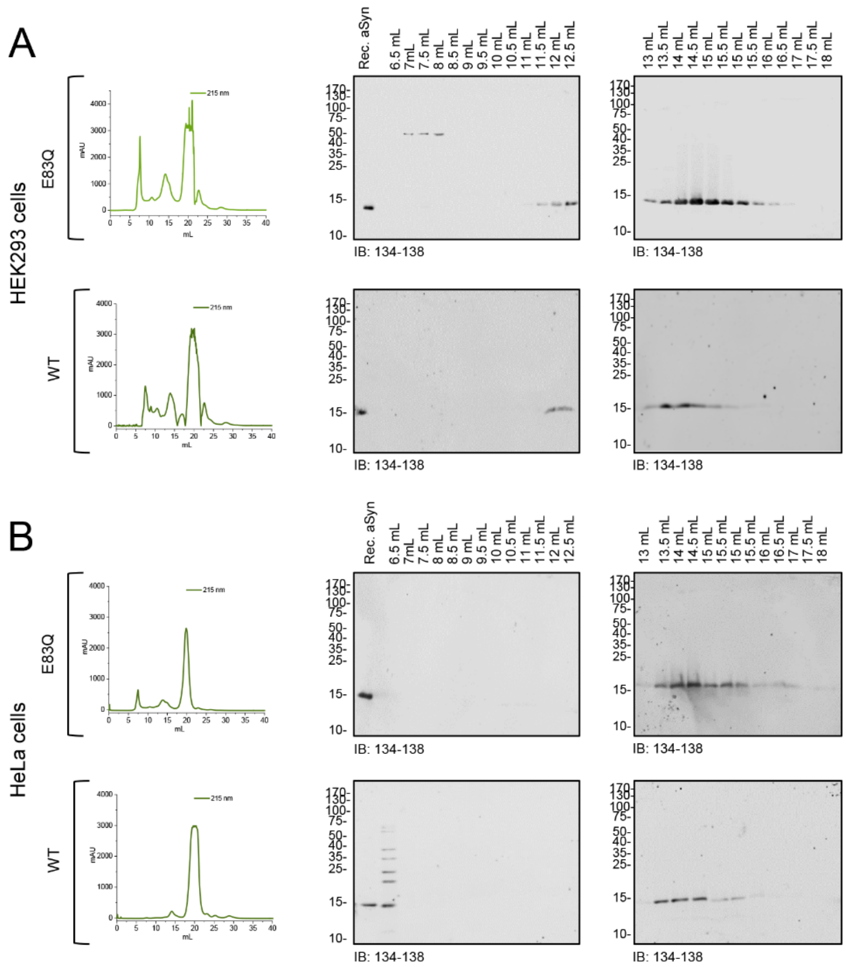


**Fig. S7. SEC analysis of cells extracts overexpressing WT and E83Q aSyn. (A)** HEK293 cells and **(B)** HeLa cells.

**Figure S8**

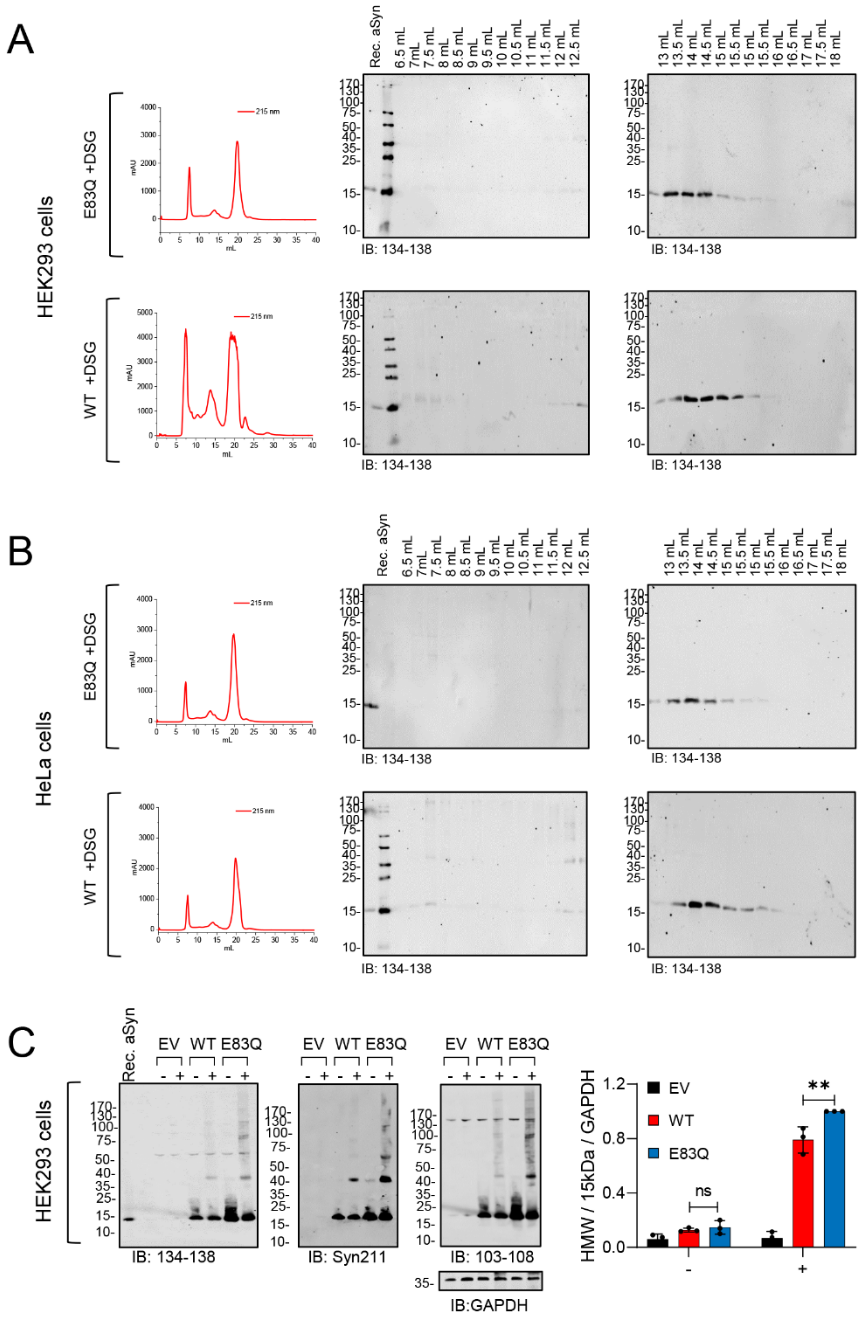

**Fig. S8. SEC analysis of the cross-linked samples (+1mM). (A)** HEK293 cells and **(B)** HeLa cells. aSyn monomer is the most predominant aSyn form observed by WB after SEC analysis, indicating that aSyn monomer remains the predominant species upon cross-linking as well as HMW species represent the minor species and thus are not present in sufficient quantities to be purified and isolated. **(C)** WB analysis following the cross-linking (DSG) analysis of HEK cells, overexpressing either an empty vector (EV), WT or E83Q constructs. HMW species of aSyn are observed in the overexpression of aSyn WT and E83Q in the presence of a cross-linker. Graphs on the right-hand side show the quantification of HMWs aSyn bands detected from and above 25 kDa to the top of the gel, HEK. Data represent the mean +/- SD of 3 independent experiments. aSyn 103-108 antibody was used as a reference to perform the densitometry. Non-specific (ns), $p<0.005$=** (ANOVA followed by Tukey HSD *post-hoc* test, comparing non-treated cells [(WT vs. E83Q) and treated cells with DSG (WT+ vs. E83Q +)] in both cell lines. "+" cells treated with 1mM DSG; "-" cells treated with DMSO.

**Figure S9**

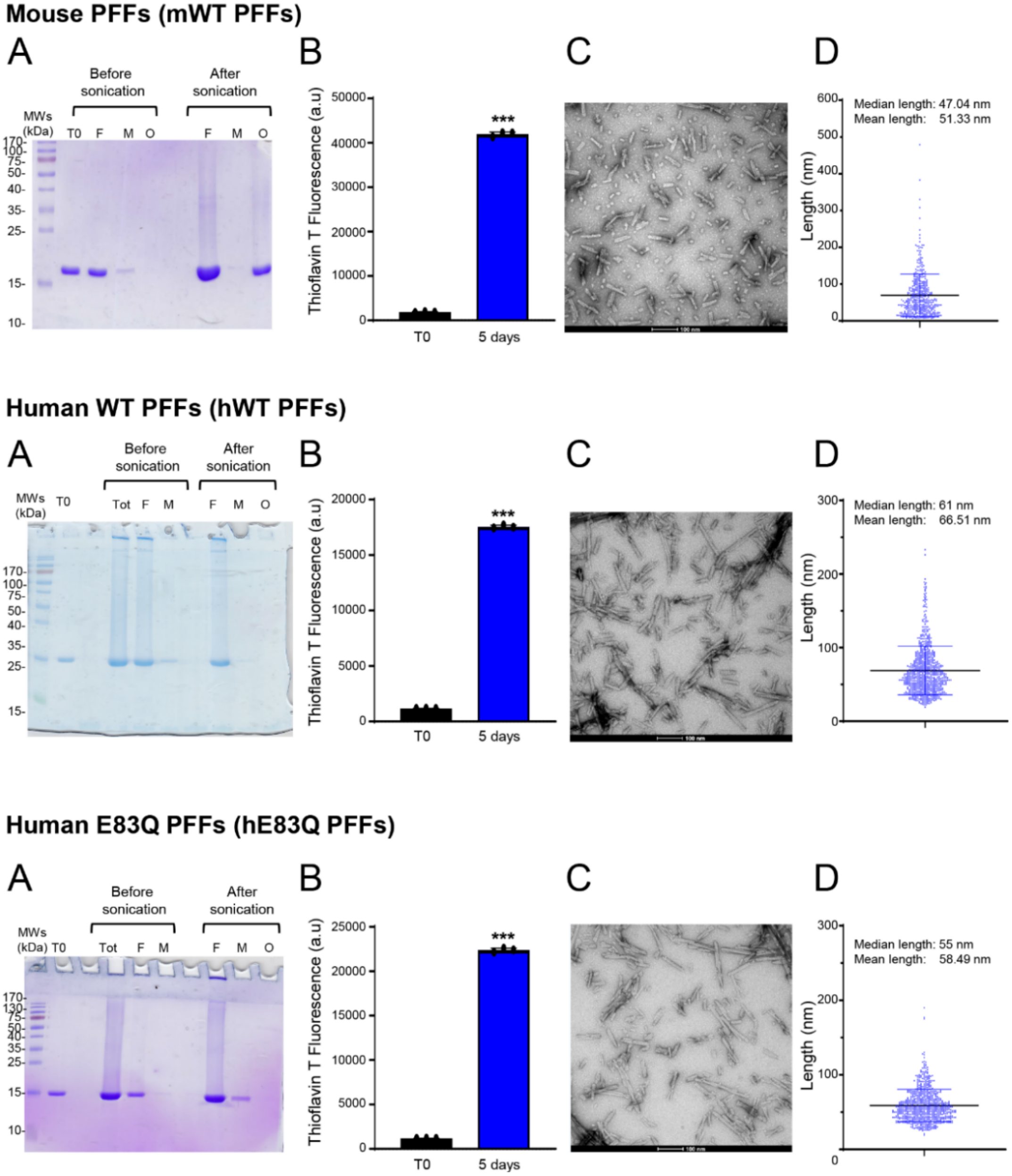


**Fig. S9. Preparation and characterization of recombinant monomeric and PFFs aSyn species used in the primary neuronal model (related to Figures 7-9)A)** Purity and characterization of aSyn monomers. Recombinant WT mouse aSyn or WT human aSyn or E83Q human aSyn was produced in *E. coli* and purified by anion exchange chromatography and size-exclusion chromatography, followed by a final chromatographic step using reverse-phase HPLC, as previously described [1]. The purity of recombinant monomeric aSyn after purification was assessed by ESI-LC/MS, which showed the expected mass. **(B-E)** Purity and characterization of aSyn fibrils. aSyn fibrils were formed by incubation of monomeric aSyn for 5 days at 37°C under constant agitation at 1000 rpm. **(B)** After sonication, fibril formation was assessed by ThT fluorometry. All data represent the average ± SD (n=3). **(C)** Purity of aSyn fibrils was verified by SDS-PAGE and Coomassie blue staining. After sonication, fibril preparations were centrifuged, and the presence of the fibrils was verified in the pellet fraction, while the absence of monomer release after the sonication step was assessed in the supernatant fraction or after filtration through a 100 kDa filter (filtration). **(D-E)** aSyn fibrils were characterized by transmission electron microscopy (TEM) imaging. **(D)** Representative images of negatively stained aSyn fibrils before and after sonication. All aSyn fibrils showed the characteristic rigid non-branched fibrillar morphology. Scale bars = 100 nm. **(E)** Average length of the fibrils after sonication.

**Figure S10**

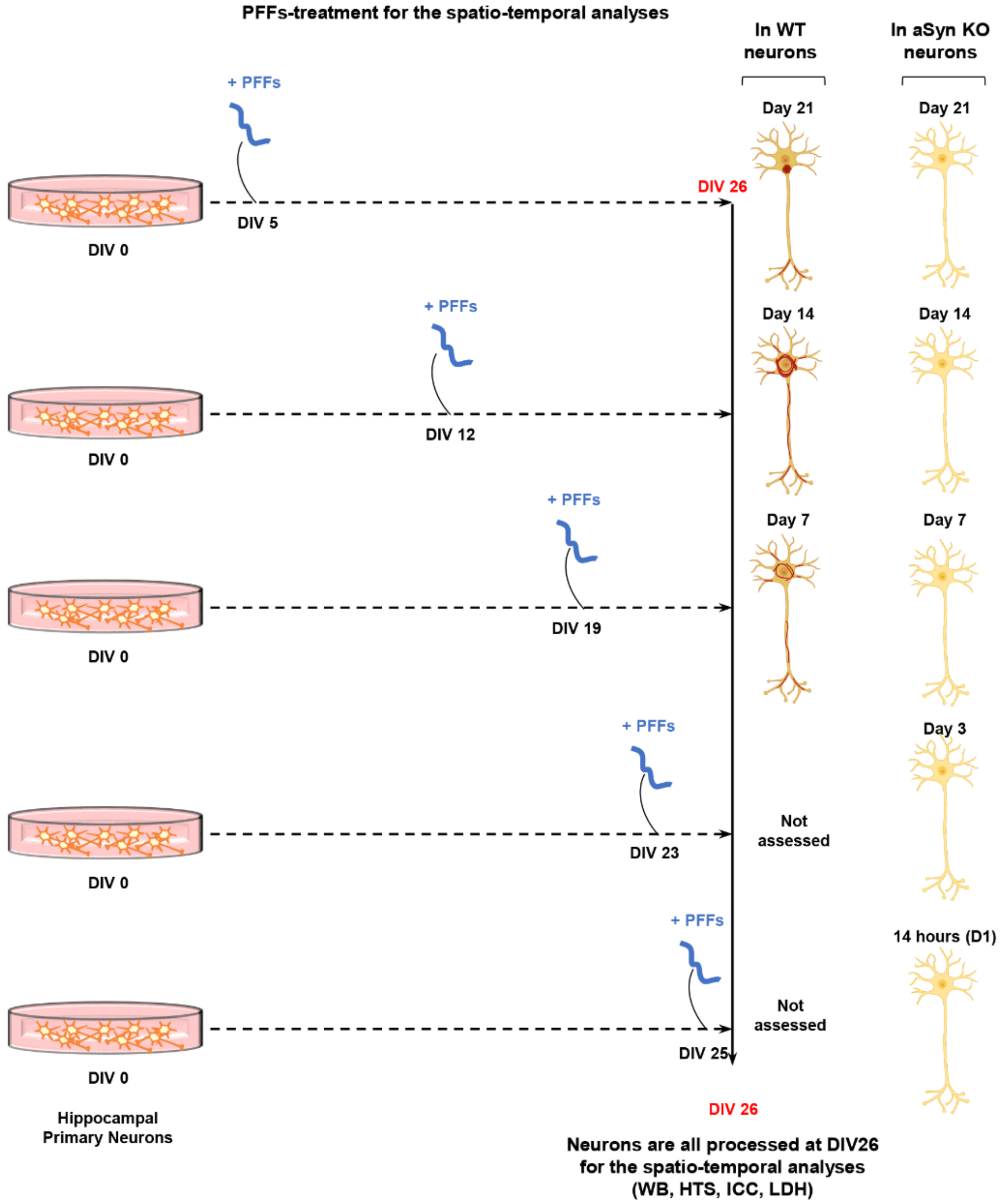


**Fig. S10. PFFs treatment in WT or aSyn KO neuronal primary culture.** PFFs were added to the neuronal cell culture media (WT neurons or aSyn KO neurons) at different DIV, and PFFs-treated neurons were all harvested at the DIV 26. This ensures that the developmental stage of primary neurons in culture was similar at the harvest time.

Figure S11

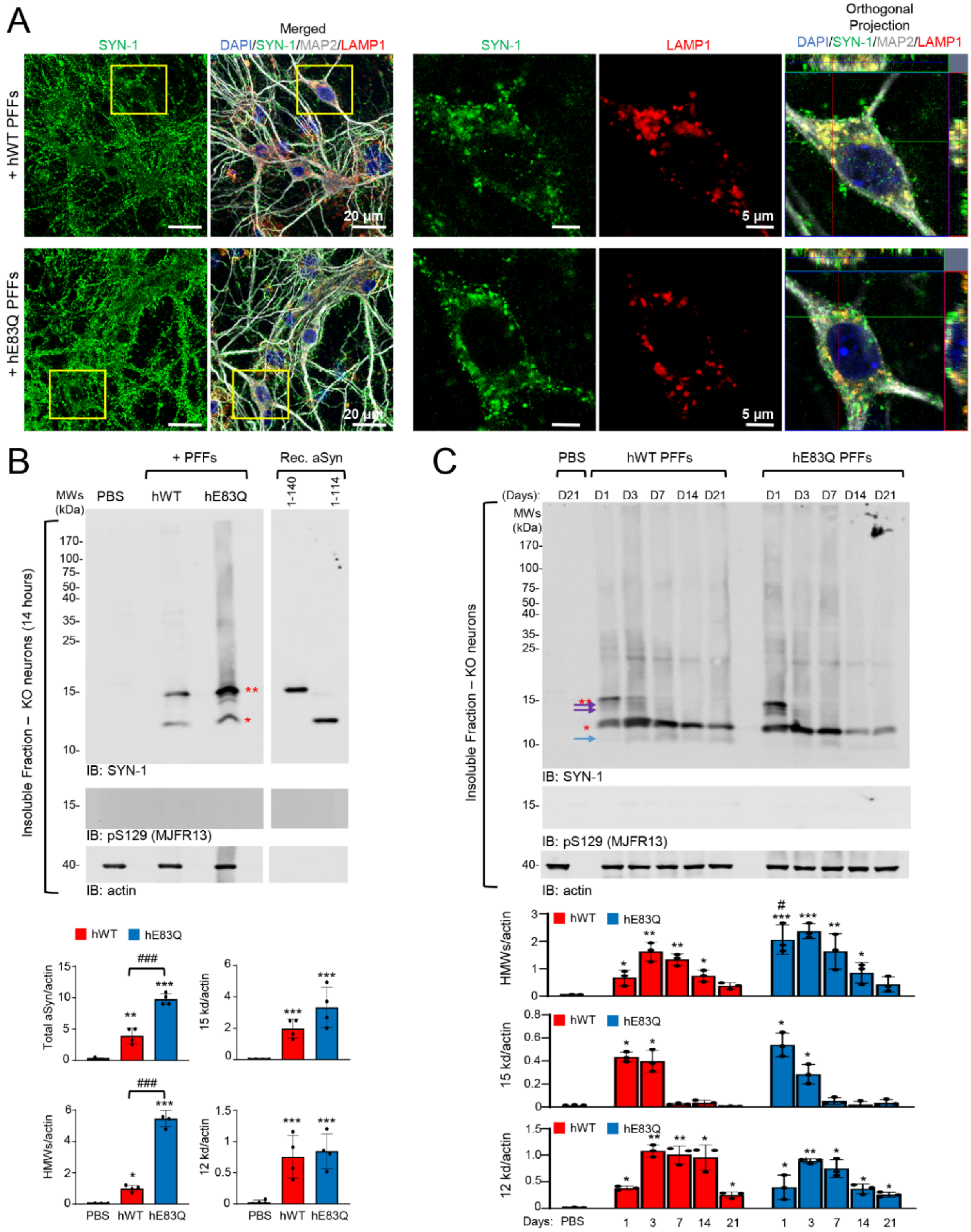


**Fig. S11. E83Q PFFs are efficiently internalized and processed in primary neurons.** 70 nM of human (h) WT (hWT) or E83Q (hE83Q) sonicated PFFs or PBS (negative control) were added to aSyn KO neurons at DIV 5 (days *in vitro*) for 14 hours **(B)**, 24 hours **(A)** and up to D21 **(C)**. **(A)** ICC confirmed the internalization of PFFs inside the neurons using a total aSyn antibody (SYN-1; epitope: 91-99). Neurons were counterstained with microtubule-associated protein (MAP2) antibody and the late lysosomes compartment with the LAMP1 antibody. The nucleus was counterstained with DAPI staining. Scale bars = 20 µm and 5 µm. Neurons imaged at higher magnification (right-hand side panel) are shown in the yellow square in the left-hand panel. **(B)** WB analysis of internalized hWT and hE83Q PFFs in aSyn KO primary neurons. After sequential extractions of nααeuronal cell lysates, the insoluble fractions were analyzed by immunoblotting. Total aSyn, pS129, and actin were detected by SYN-1, pS129 (MJFR13), and actin antibodies, respectively. Monomeric ayn (15 kDa) is indicated by a double asterisk; C-terminal truncated aSyn (12 kDa) is indicated by a single asterisk; the higher molecular weights (HMWs) corresponding to the newly formed fibrils are detected from 25 kDa to the top of the gel. 20 ng of recombinant monomeric aSyn full-length or C-terminally truncated at residue 114 (1-114) were loaded to assess the size of the truncated PFFs. **(C)** Fate and clearance of hWT and hE83Q PFFs after internalization was followed by WB analyses of the insoluble fraction of the PFF-treated aSyn KO neurons up to 21 days (D21). Control neurons were treated with PBS buffer (PBS). Total aSyn and pS129 were detected by SYN-1, pS129 (MJFR13). The total loading level was assessed using the actin antibody. Full-length aSyn (15 kDa) is indicated by a double red asterisk, the C-terminal truncated species (12 kDa) by a single red asterisk. The purple arrows indicate the intermediate aSyn-truncated fragments and the blue arrow the 10 kDa species. **(B-C)** The graphs represent the mean +/- SD of a minimum of 3 independent experiments. $p<0.05$=*, $p<0.005$=**, $p<0.0005$=*** (ANOVA followed by Tukey HSD *post-hoc* test, PBS vs. PFF-treated neurons). $p<0.05$=#, $p<0.0005$=### (ANOVA followed by Tukey HSD *post-hoc* test, hWT PFF-treated neurons vs. hE83Q PFF-treated neurons).

**Figure S12**

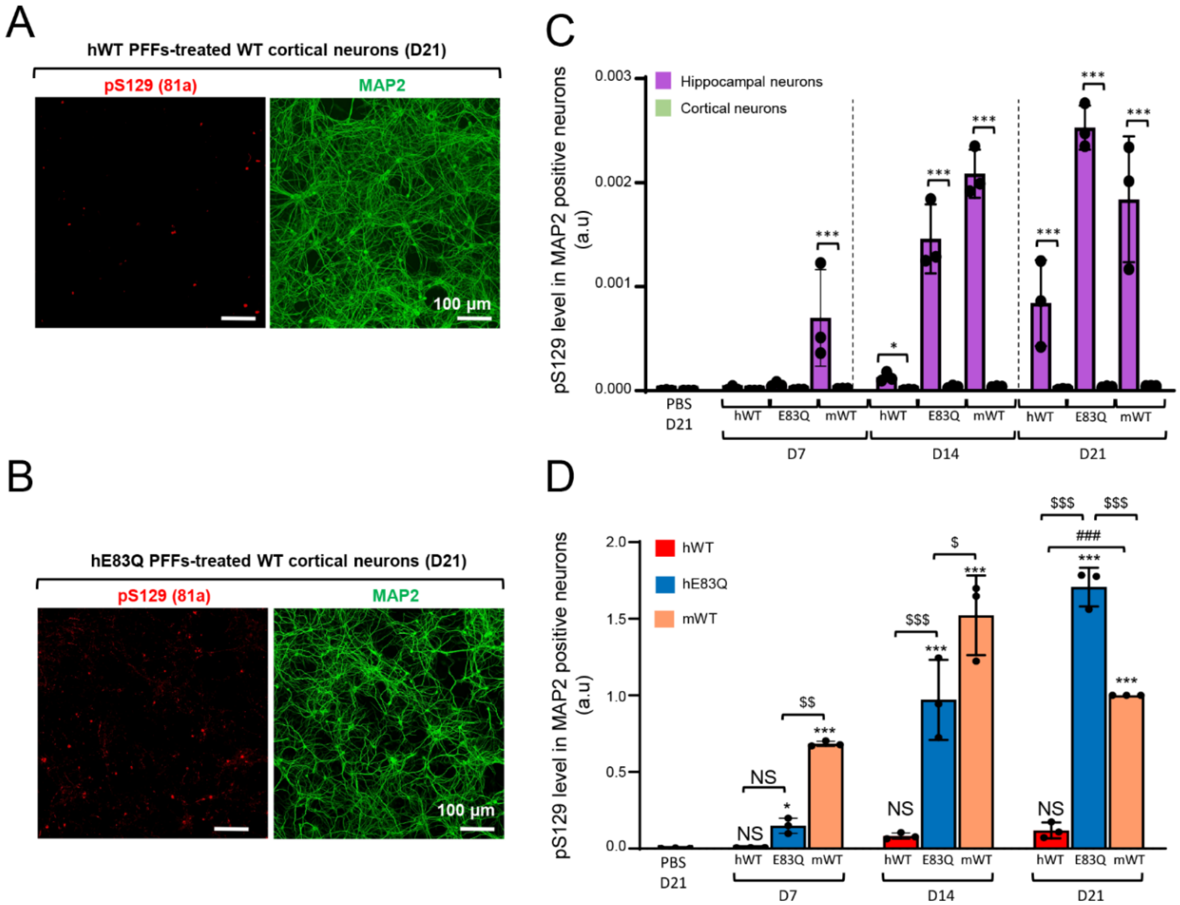


**Fig. S12. E83Q mutation is enough to restore the capacity of human PFFs to seed as efficiently as mouse PFFs in primary cortical neurons.** Temporal analysis of the level of aSyn seeded-aggregates formed in WT cortical **(A-B, D)** or hippocampal **(C)** neurons after the addition of 70 nM of hWT, hE83Q or mWT aSyn PFFs to the primary culture (at DIV 5) for 7 days (D7), 14 days (D14) or 21 days (D21). Control neurons were treated with PBS. **(A-D)** The level of seeded-aggregates was measured overtime in PBS- or PFF-treated neurons by high content analysis (HCA). Seeded-aggregates were detected by ICC using pS129 (81a) antibody and neurons were counterstained with microtubule-associated protein (MAP2) antibody, and the nucleus was counterstained with DAPI staining. For each independent experiment, a minimum of two wells was acquired per condition, and nine fields of view were imaged for each well. 3 independent experiments were performed. Representative images of seeded-aggregates formed at D14 **(A)** and at D21 **(B)** in primary cortical neurons. Scale bars = 100 µm. **(C-D)** Images were then analyzed using Cell Profiler software to quantify the total level of pS129 in MAP2 positive neurons as previously described [2]. **(C-D)** The graphs represent the mean +/- SD of a minimum of 3 independent experiments. **(C)** p<0.05=*, p<0.005=**, p<0.0005=*** (ANOVA followed by Tukey HSD *post-hoc* test, hippocampal vs. cortical neurons). (**D**) p<0.05=*, p<0.0005=*** (ANOVA followed by Tukey HSD *post-hoc* test, PBS vs. PFF-treated neurons). p<0.05=$, p<0.005=$$, p<0.0005=$$$ (ANOVA followed by Tukey HSD *post-hoc* test, E83Q PFF-treated neurons vs. hWT or mWT PFF-treated neurons). N.S (non-specific, D14 vs D21). p<0.0005=### (ANOVA followed by Tukey HSD *post-hoc* test, hWT PFF-treated neurons vs. mWT PFF-treated neurons).

**Figure S13**

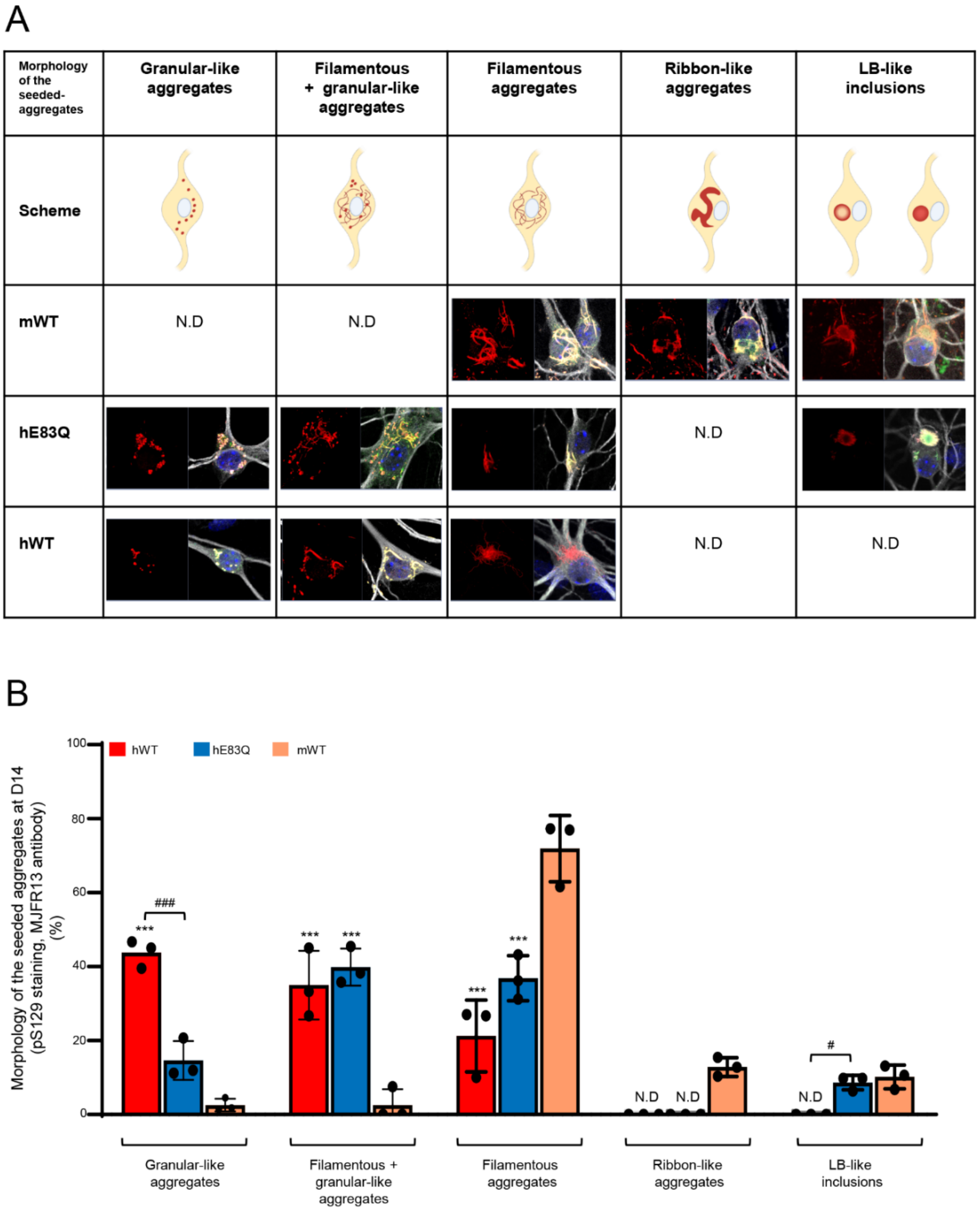

A
Morphology of the seeded-aggregates
Granular-like aggregates
Filamentous + granular-like aggregates
Filamentous aggregates
Ribbon-like aggregates
LB-like inclusions
Scheme
mWT
N.D
N.D
hE83Q
N.D
hWT
N.D
N.D
B
hWT
hE83Q
mWT
100
80
60
40
20
0
Morphology of the seeded aggregates at D14 (pS129 staining, MJFR13 antibody) (%)
###
***
***
***
***
***
#
N.D
N.D
N.D
Granular-like aggregates
Filamentous + granular-like aggregates
Filamentous aggregates
Ribbon-like aggregates
LB-like inclusions

**Figure S13**

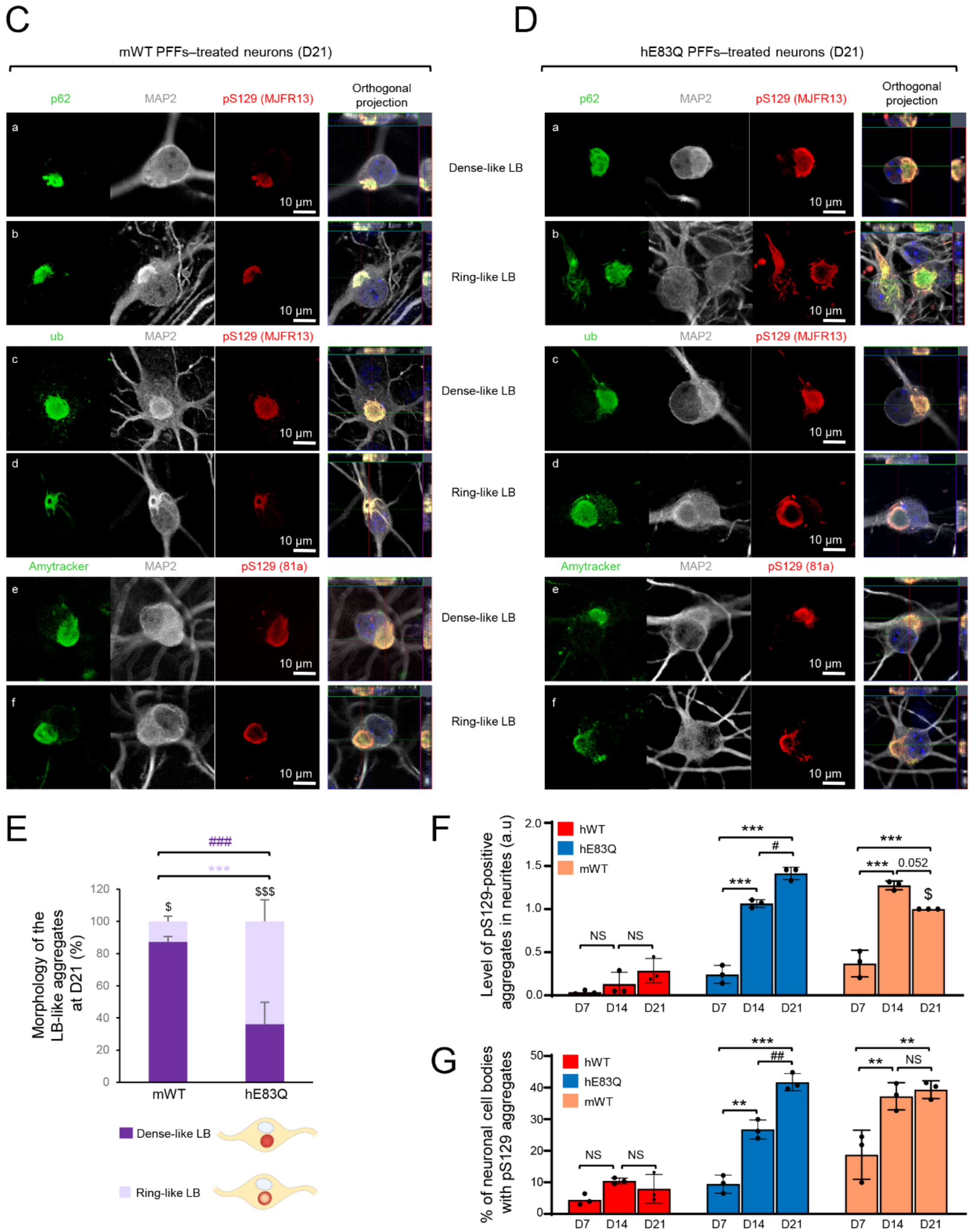


**Fig. S13**. **Quantification of the different types of morphologies of the aSyn seeded-aggregates formed in PFF-treated neurons at D14 and D21. (A)** Classification of the different types of morphologies of the aSyn seeded-aggregates formed in primary neurons and representative scheme for each type of morphology. **(B)** Quantification of the different types of morphologies of the aSyn seeded-aggregates observed by ICC in primary neurons at D14. A minimum of 100 neurons was counted in each experiment. The graphs represent the mean +/- SD of a minimum of three independent experiments. p<0.0005=*** (ANOVA followed by Tukey HSD *post-hoc* test, mWT PFF-treated neurons vs. human PFFs (hWT or E83Q)-treated neurons). p<0.05=#, p<0.0005=### (ANOVA followed by Tukey HSD *post-hoc* test, hE83Q PFF-treated neurons vs. hWT PFF-treated neurons). **(C, D)** LB-like inclusions were detected by ICC using pS129 (MJFR13) in combination with p62 (a–d), ubiquitin (e–h), Amytracker dye (i–l) in mWT PFF **(C)** (a–b, e–f, i–j) or hE83Q PFF **(D)** (c–d, g–h, k–l)-treated neurons. Neurons were counterstained with MAP2 antibody, and the nuclei were counterstained with DAPI. Scale bars = 10 μm. **(E)** Quantification of the morphologies (LB-like inclusions with a dense core or LB-like inclusions with a ring-like structure) of the LB-like inclusions observed by ICC in primary neurons at D21. A minimum of 50 neurons was counted in each experiment (n=3). The graphs represent the mean +/- SD of a minimum of three independent experiments. p<0.0005=*** (ANOVA followed by Tukey HSD *post-hoc* test, ring-like structures counted in mWT PFF-treated neurons vs. E83Q PFF-treated neurons). p<0.0005=### (ANOVA followed by Tukey HSD *post-hoc* test, dense core-like structures counted in mWT PFF-treated neurons vs. E83Q PFF-treated neurons). p<0.05=$, p<0.0005=$$$ (ANOVA followed by Tukey HSD *post-hoc* test, ring-like vs. dense core-like LB in hE83Q PFF-treated neurons or mWT PFF-treated neurons). N.D for not detected. **(F-G)** The level of seeded-aggregates in neurites **(F)** and in the neuronal cell bodies **(G)** was measured overtime in PBS- or PFF-treated neurons by high content analysis (HCA). Seeded-aggregates were detected by ICC using pS129 (81a) antibody and neurons were counterstained with microtubule-associated protein (MAP2) antibody, and the nucleus was counterstained with DAPI staining. Neurites and neuronal cell bodies segmentations were performed as previously described[2]. For each independent experiment, a minimum of two wells was acquired per condition, and nine fields of view were imaged for each well. 3 independent experiments were performed. The graphs represent the mean +/- SD of a minimum of three independent experiments. p<0.0005=***, p<0.005=** (ANOVA followed by Tukey HSD *post-hoc* test, D7 vs. D14 or D21). p<0.0005=# (ANOVA followed by Tukey HSD *post-hoc* test, D14 vs. D21). NS for non-specific.

**Figure S14**

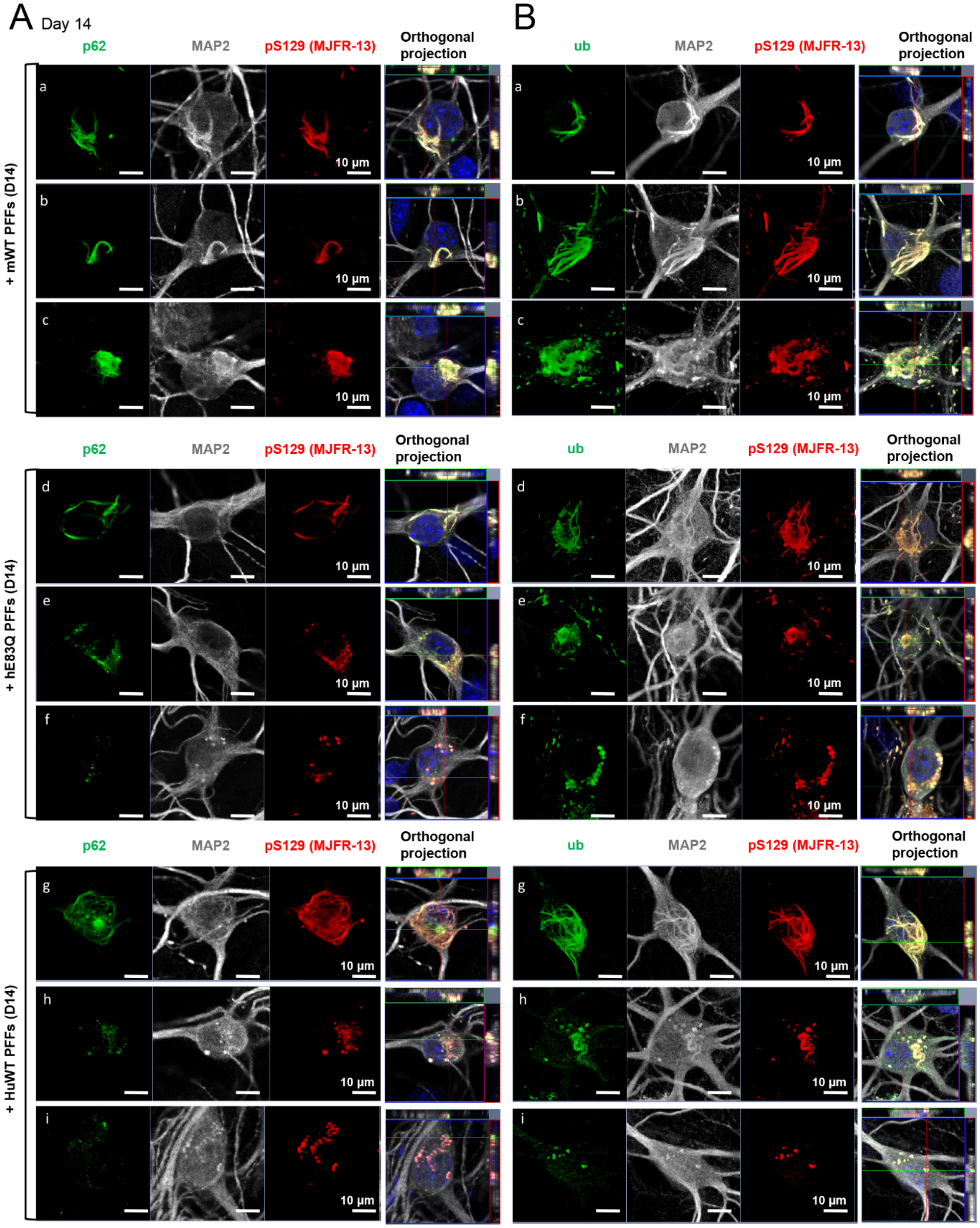

A
Day 14
p62
MAP2
pS129 (MJFR-13)
Orthogonal projection
+ mWT PFFs (D14)
+ hE83Q PFFs (D14)
+ HuWT PFFs (D14)
a
b
c
d
e
f
g
h
i
10 µm
B
ub
MAP2
pS129 (MJFR-13)
Orthogonal projection

**Figure S14**

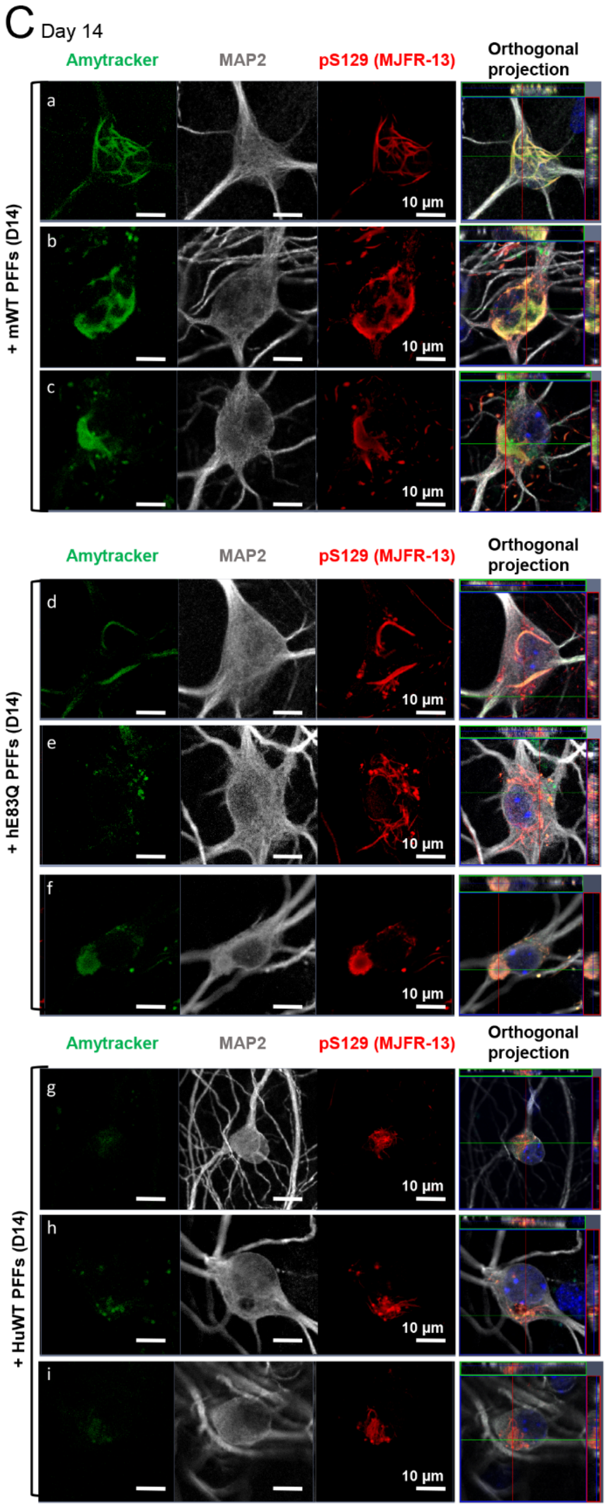

C Day 14
Amytracker
MAP2
pS129 (MJFR-13)
Orthogonal projection
+ mWT PFFs (D14)
a
b
c
10 µm
+ hE83Q PFFs (D14)
d
e
f
+ HuWT PFFs (D14)
g
h
i

Figure S14

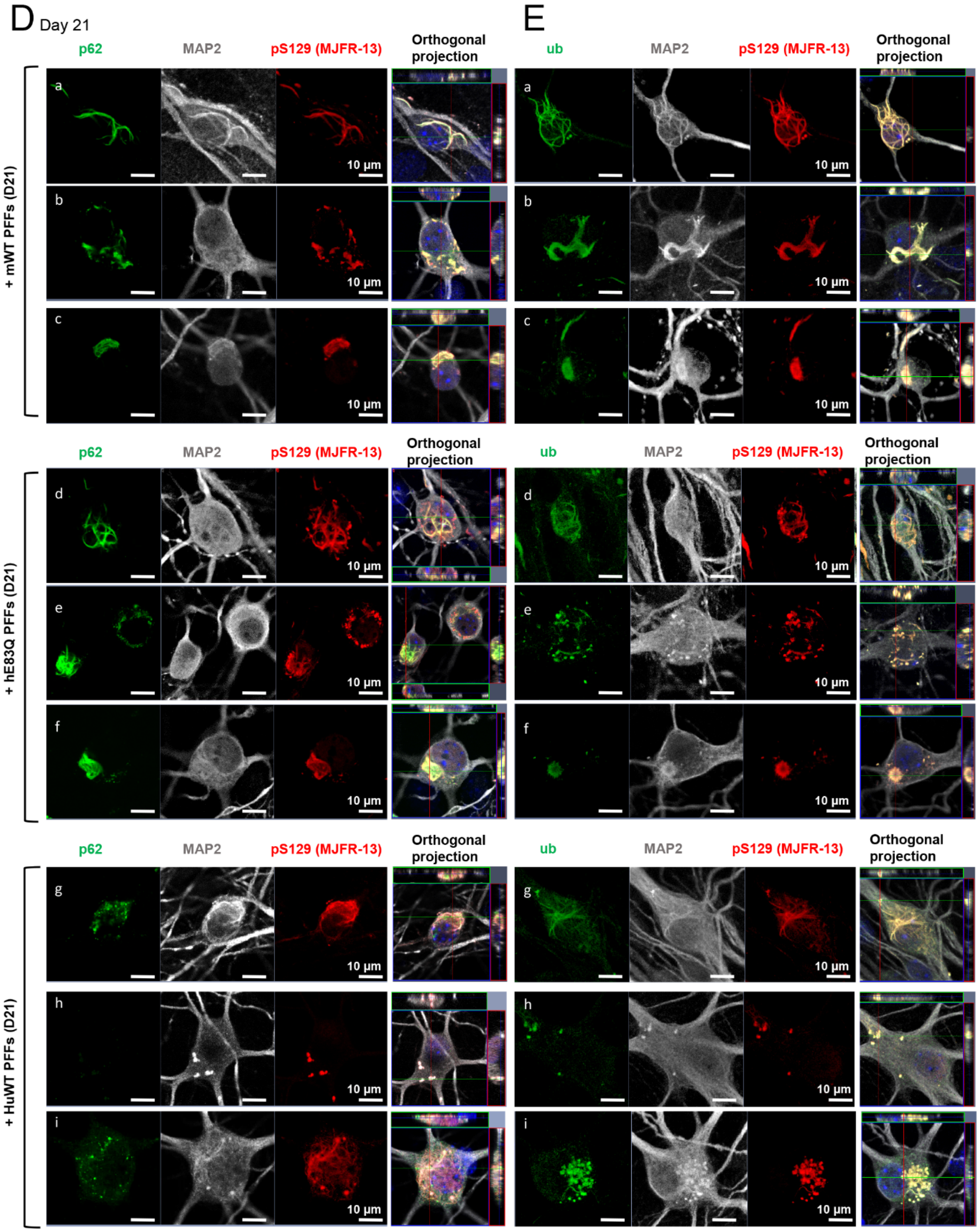

D Day 21
p62
MAP2
pS129 (MJFR-13)
Orthogonal projection
+ mWT PFFs (D21)
a
b
c
d
e
f
g
h
i
10 µm
+ hE83Q PFFs (D21)
+ HuWT PFFs (D21)
E
ub

**Figure S14**

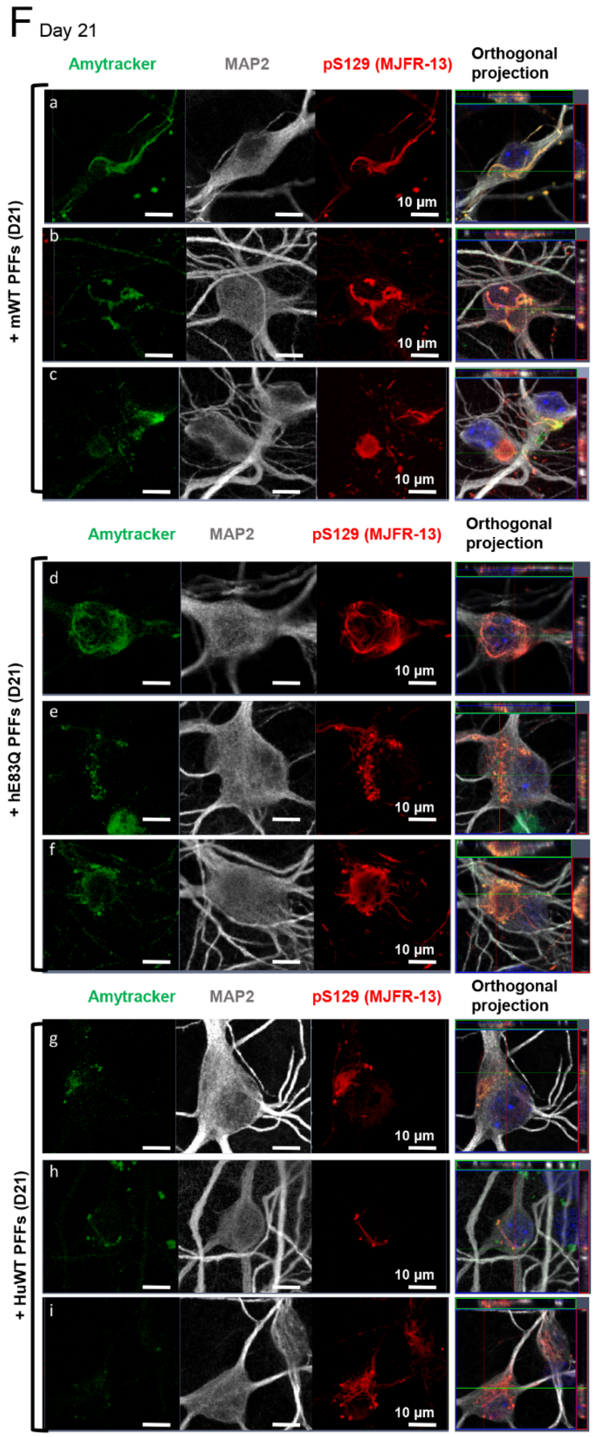


**Fig. S14. Temporal characterization of the seeded-aggregates formed in PFF-treated neurons using pathological markers of the LB. (A-F)** Temporal analysis of the aSyn seeded-aggregates formed in WT neurons after the addition of 70 nM of mouse (m) WT or hE83Q or hWT aSyn PFFs to the primary culture (at DIV 5) for 14 days (D14, **A-C**) and 21 days (D21, **D-F**). Aggregates were detected by ICC using pS129 (MJFR13 or 81a) in combination with p62 **(A, D)**, ubiquitin **(B, E)** or Amytracker dye **(C, F)** in mWT (**a-c**), hE83Q (**d-f**), hWT (**g-i**) PFF-treated neurons. Neurons were counterstained with microtubule-associated protein (MAP2) antibody, and the nucleus was counterstained with DAPI staining. Scale bars = 10 µm.

**Figure S15**

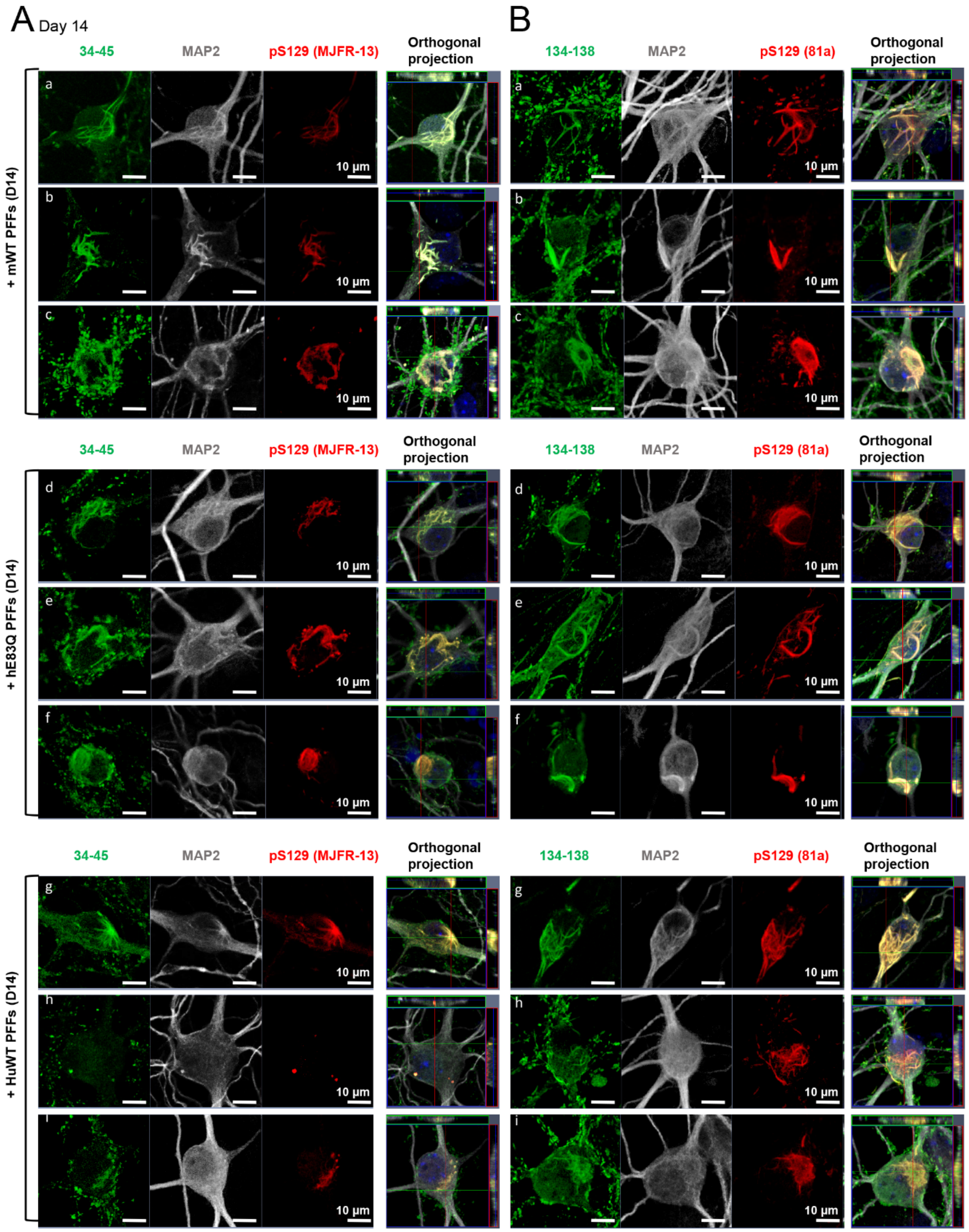

A Day 14
B
34-45
MAP2
pS129 (MJFR-13)
Orthogonal projection
134-138
MAP2
pS129 (81a)
Orthogonal projection
+ mWT PFFs (D14)
+ hE83Q PFFs (D14)
+ HuWT PFFs (D14)
a
b
c
d
e
f
g
h
i
10 µm

**Figure S15**

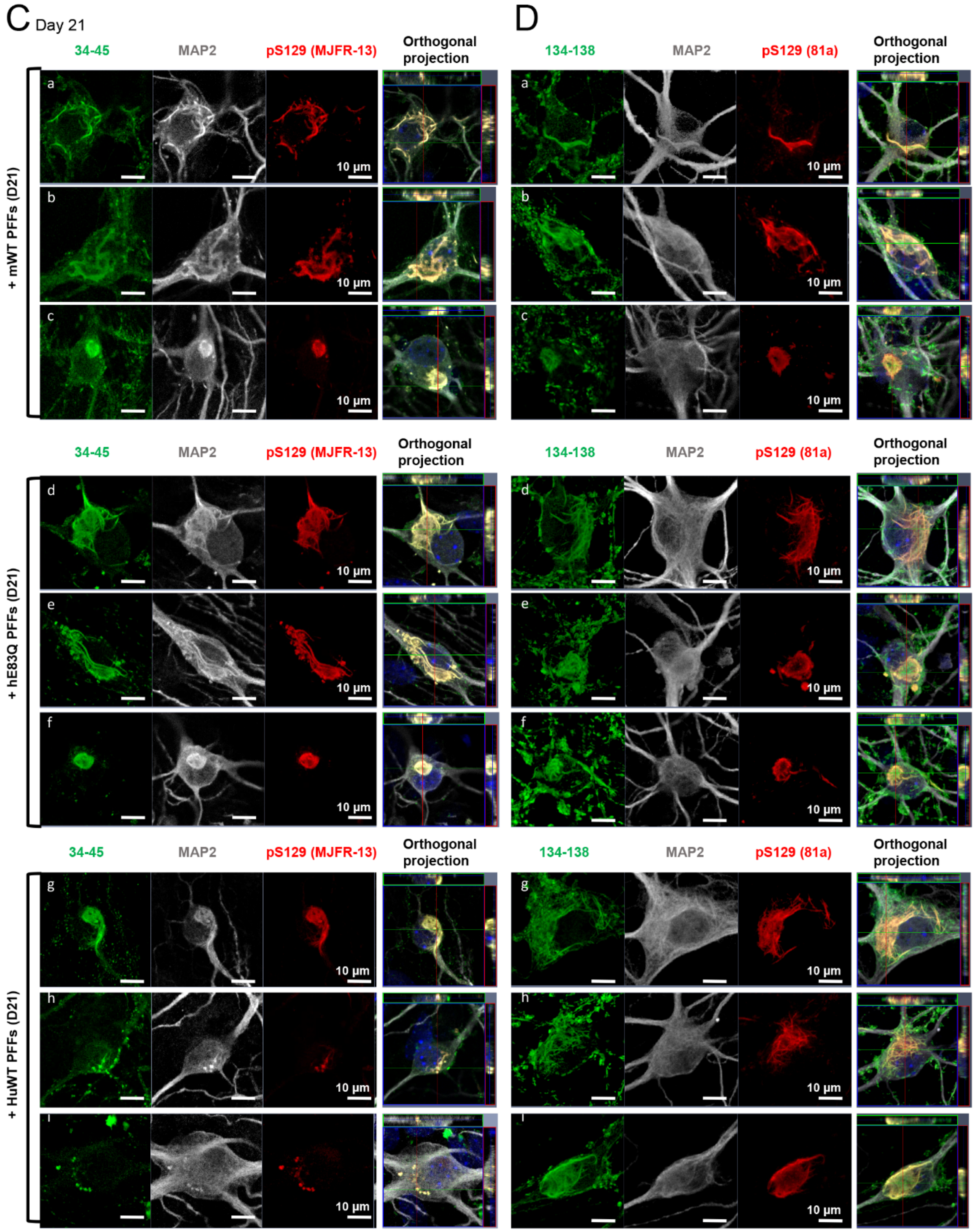


**Fig. S15. Temporal characterization of the seeded-aggregates formed in PFF-treated neurons with N-ter and C-ter total aSyn**. **(A-D)** Temporal analysis of the aSyn seeded-aggregates formed in WT neurons after the addition of 70 nM of hWT or hE83Q or mouse (m) WT aSyn PFFs to the primary culture (at DIV 5) for 14 days (D14, **A-B**) and 21 days (D21, **C-D**). Aggregates were detected by ICC using pS129 (MJFR13 or 81a) in combination with total N-ter aSyn (epitope: 34-45) (**A, C**), or total C-ter (epitope: 134-138) (**B, D**) aSyn antibodies in mWT (**a-c**), hE83Q (**d-f**), hWT (**g-i**) PFF-treated neurons. Neurons were counterstained with microtubule-associated protein (MAP2) antibody, and the nucleus was counterstained with DAPI staining. Scale bars = 10 µm.

**Figure S16**

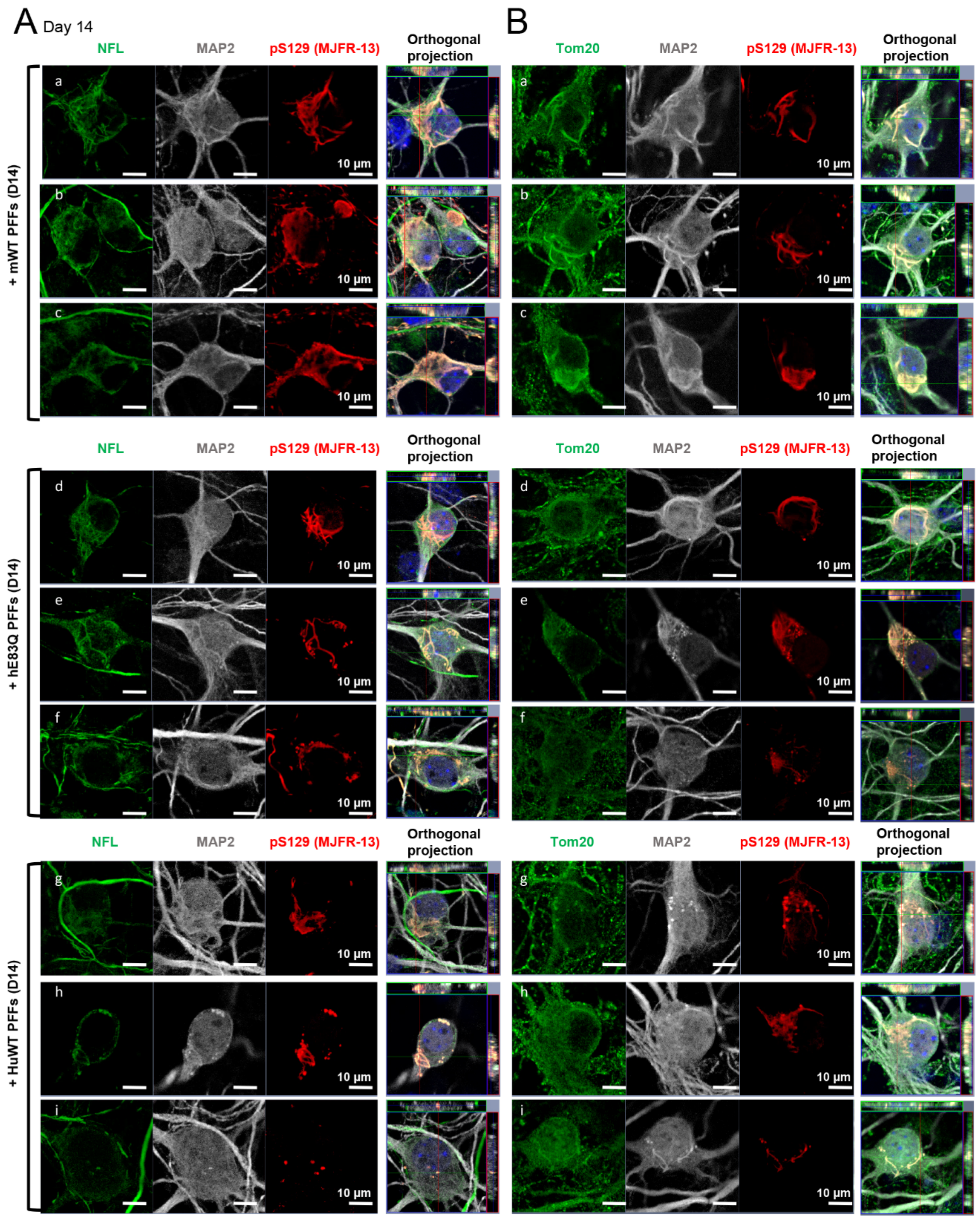

A Day 14
B
NFL
MAP2
pS129 (MJFR-13)
Orthogonal projection
Tom20
+ mWT PFFs (D14)
+ hE83Q PFFs (D14)
+ HuWT PFFs (D14)
a
b
c
d
e
f
g
h
i
10 µm

**Figure S16**

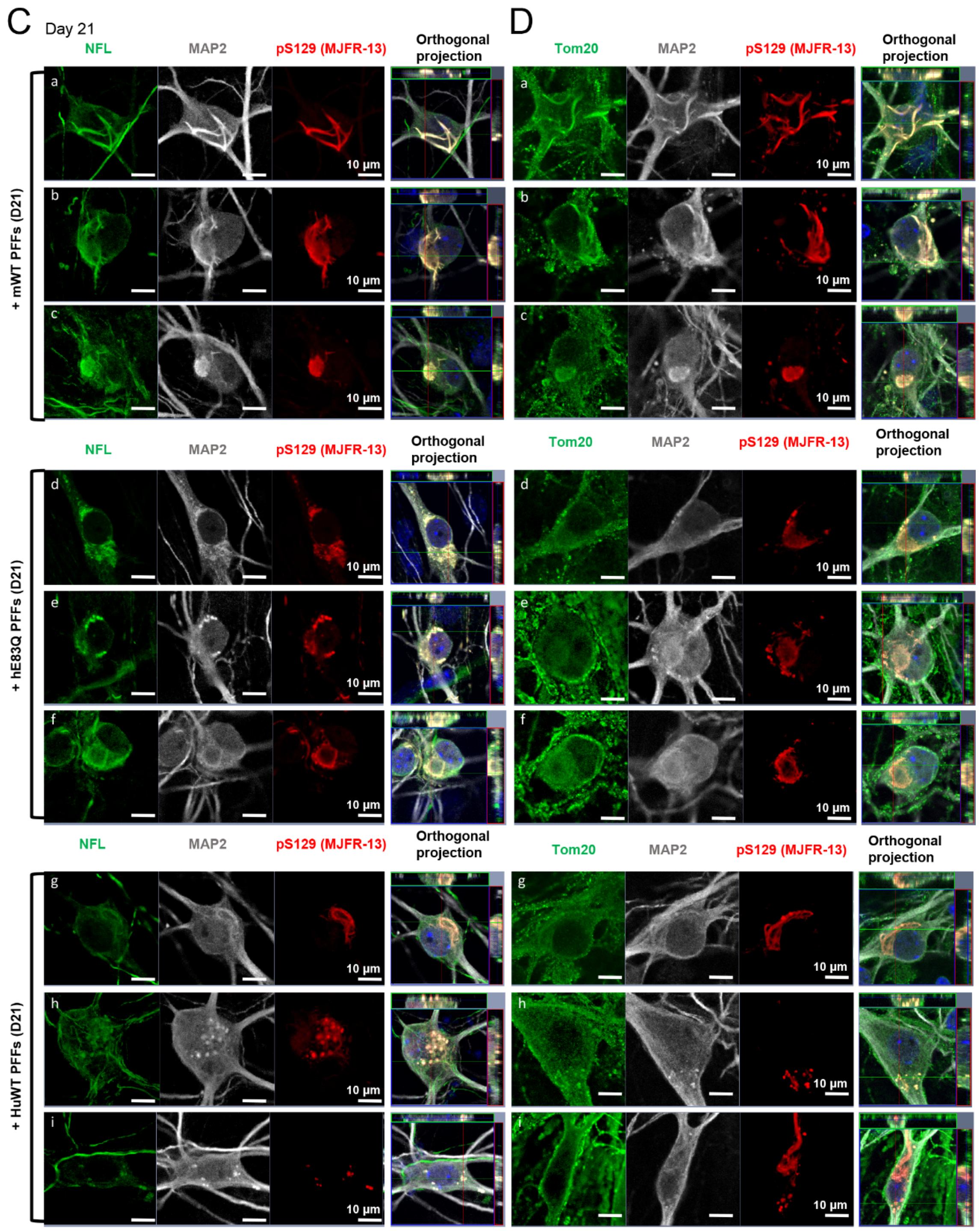


**Fig. S16. Temporal characterization of the seeded-aggregates formed in PFF-treated hippocampal neurons with neurofilaments and mitochondria markers. (A-D)** Temporal analysis of the aSyn seeded-aggregates formed in WT neurons after the addition of 70 nM of hWT or hE83Q or mouse (m) WT aSyn PFFs to the primary culture (at DIV 5) for 14 days (D14, **A-B**) and 21 days (D21, **C-D**). Aggregates were detected by ICC using pS129 (MJFR13 or 81a) in combination with NFL **(A, C)**, or with the mitochondrial marker Tom20 **(B, D)** aSyn antibodies in mWT (**a-c**), hE83Q (**d-f**), hWT (**g-i**) PFF-treated neurons. Neurons were counterstained with microtubule-associated protein (MAP2) antibody, and the nucleus was counterstained with DAPI staining. Scale bars = 10 µm.

**Figure S17**

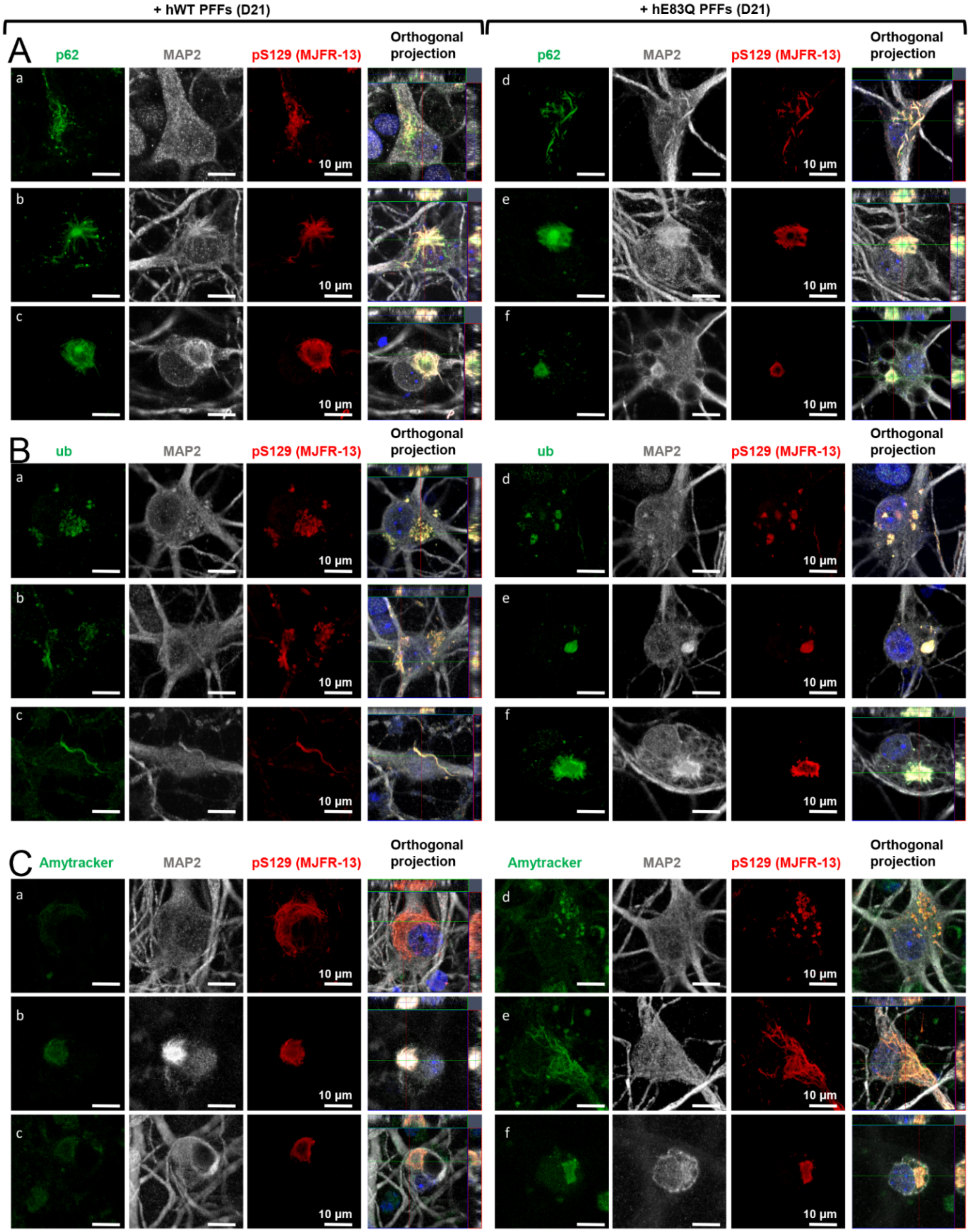
+ hWT PFFs (D21)
+ hE83Q PFFs (D21)
A
p62
MAP2
pS129 (MJFR-13)
Orthogonal projection
a
b
c
d
e
f
10 µm
B
ub
MAP2
pS129 (MJFR-13)
Orthogonal projection
C
Amytracker
MAP2
pS129 (MJFR-13)
Orthogonal projection

**Figure S17**

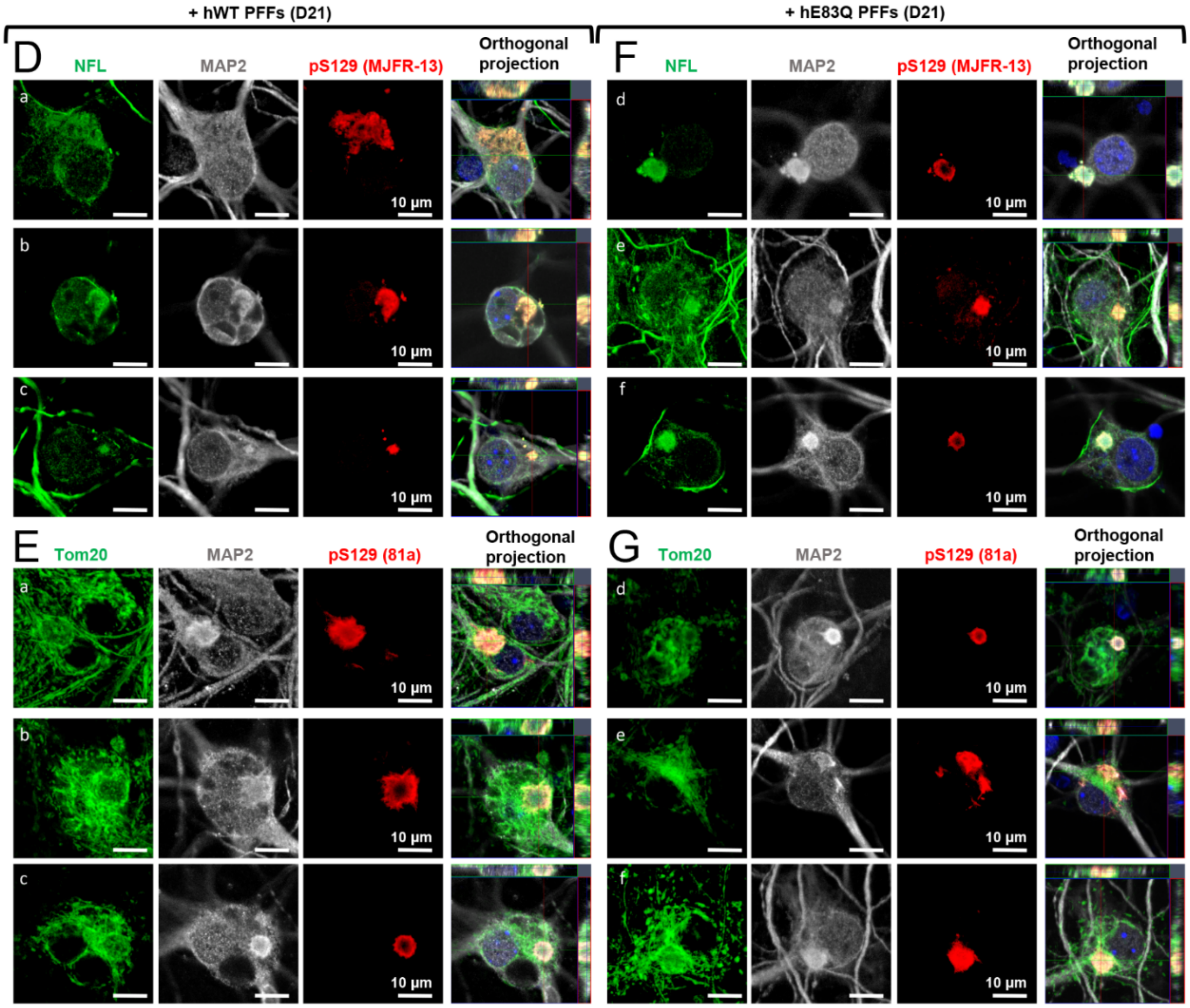


**Fig S17. Morphological characterization of the seeded-aggregates formed in PFF-treated cortical neurons using pathological markers of the LB.** 70 nM of hE83Q or hWT aSyn PFFs was added to the cortical primary culture (at DIV 5) for 21 days. Aggregates were detected by ICC using pS129 (MJFR13 or 81a) in combination with p62 **(A)**, ubiquitin **(B)**, Amytracker dye **(C)**, NFL **(D)** or with the mitochondrial marker Tom20 **(E)** in hWT (**a-c**) or E83Q (**d-f**) PFF-treated cortical neurons. Neurons were counterstained with MAP2 antibody, and the nucleus with DAPI staining. Scale bars = 10 µm.

**Figure S18**

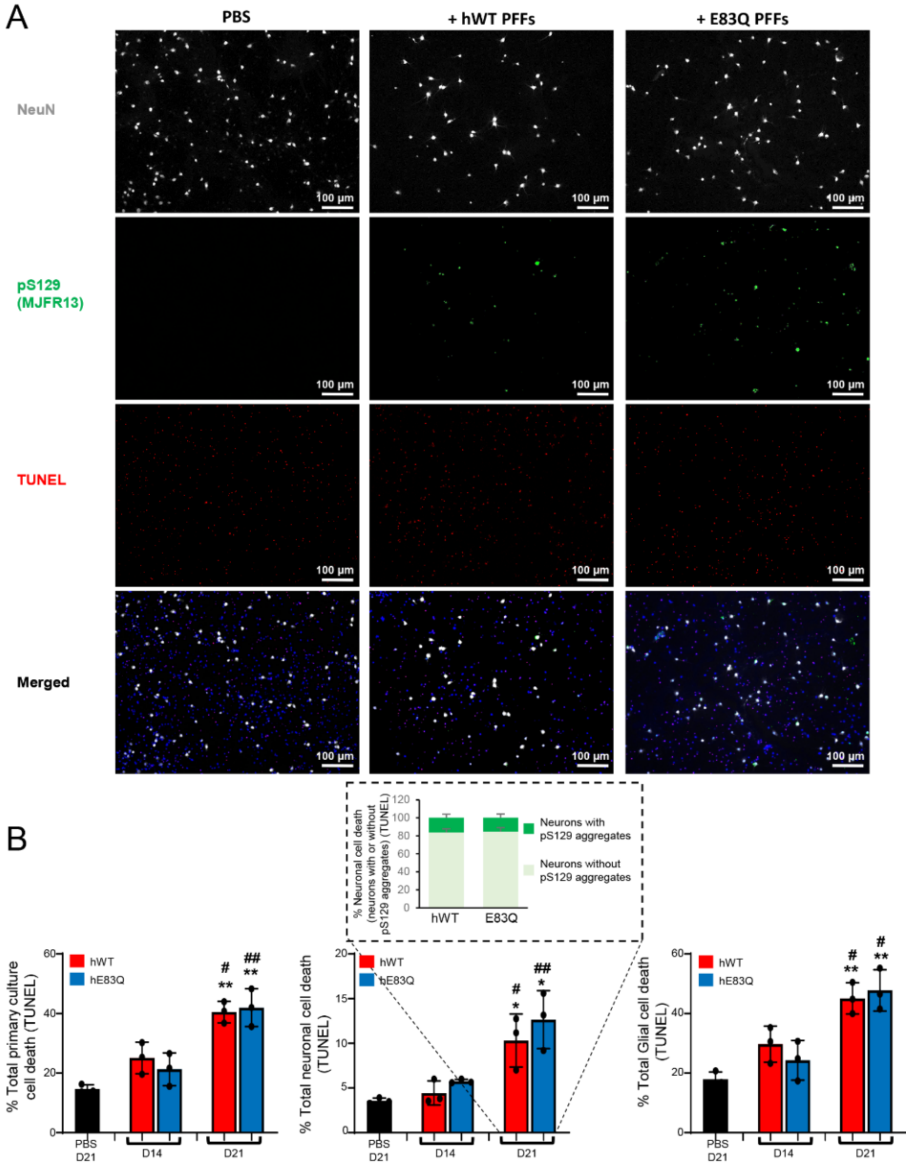


**Fig S18. Cell death quantification in PFF-treated cortical neurons.** 70 nM of hE83Q or hWT aSyn PFFs was added to the cortical primary culture (at DIV 5) for 14 and 21 days. PBS was used as a control to treat cortical neurons 21 days. **(A)** At the indicated time, PBS- and PFF-treated cortical neurons were fixed and ICC was performed. The neuronal population was specifically stained with the specific neuronal marker NeuN, neurons with seeded-aggregates were detected after pS129 staining (MJFR13 antibody), and the nuclei of the total cell population (neuron and glial cells) were counterstained with DAPI staining. In addition, the Terminal dUTP nick end-labeling (TUNEL) apoptotic cell death assay was performed for each condition. Scale bars = 100 µm. In the PBS or PFF-treated primary culture, the percentage of total cell death was quantified as follows: (TUNEL-positive nuclei/ DAPI-positive nuclei), the percentage of apoptotic neurons was quantified as follows: [(TUNEL-positive and NeuN-positive cells)/total NeuN-positive cells] and the percentage of glial cells as: [(TUNEL-positive and NeuN-negative cells)/(DAPI positive and NeuN-negative cells)]. Among the neuronal population, those in which seeded-aggregates were formed were not affected by cell death (TUNEL+ NeuN+ pS129+). **(B)** The graphs represent the mean +/- SD of three independent experiments. For each independent experiment, 3 fields of view (FOV) per condition were acquired at a 10x magnification (~400 to 1'000 cells per FOV were counted). $p<0.005$=** (ANOVA followed by Tukey HSD *post-hoc* test, PBS vs. PFF-treated neurons). $p<0.05$=# (ANOVA followed by Tukey HSD *post-hoc* test, hWT PFF-treated neurons vs. E83Q PFF-treated neurons).

**Figure S19**

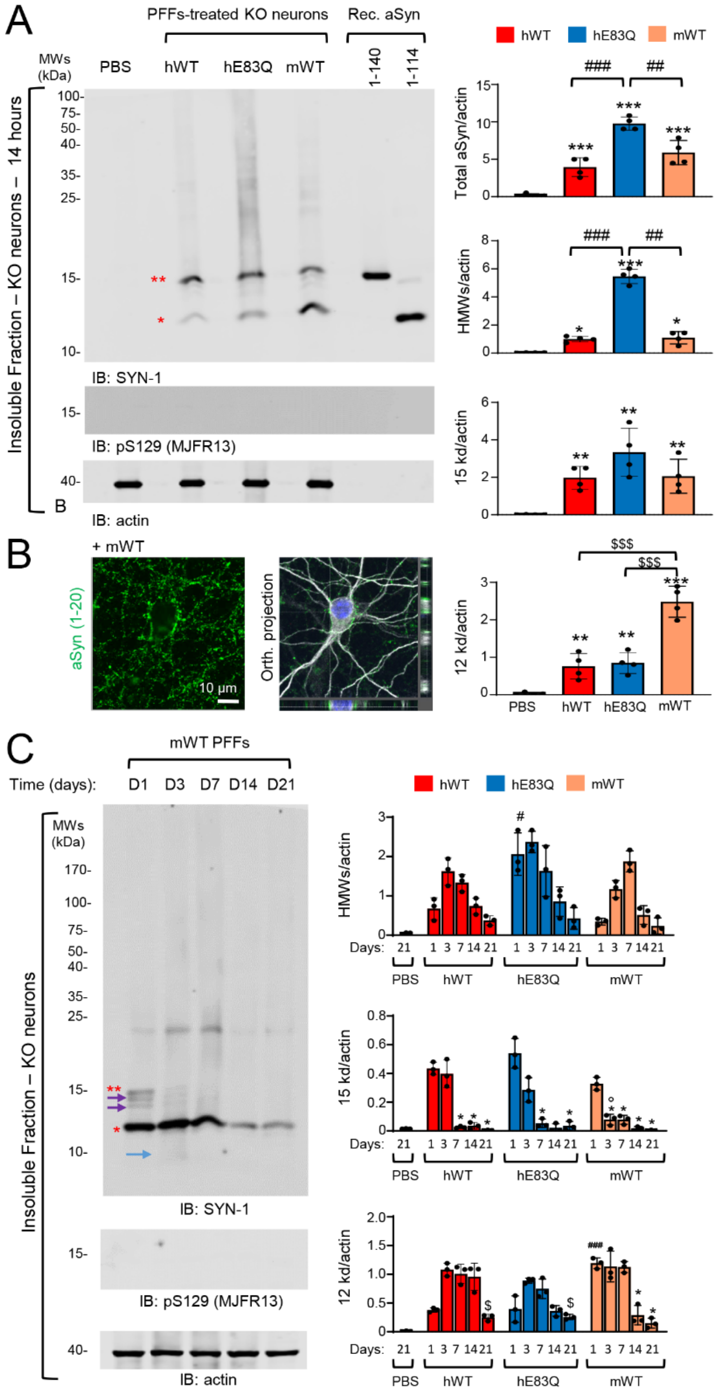


**Fig. S19. E83Q PFFs are efficiently internalized and processed in primary neurons as the mouse WT PFFs.** 70 nM of human (h) WT (hWT), hE83Q (hE83Q) or mouse WT (mWT) sonicated PFFs or PBS (negative control) were added to aSyn KO neurons at DIV 5 for 14 hours. **(A)** WB analysis of internalized hWT, hE83Q or mWT PFFs in aSyn KO primary neurons. After sequential extractions of neuronal cell lysates, the insoluble fractions were analyzed by immunoblotting. Total aSyn, pS129, and actin were detected by SYN-1, pS129 (MJFR13), and actin antibodies, respectively. Monomeric aSyn (15 kDa) is indicated by a double asterisk; C-terminal truncated aSyn (12 kDa) is indicated by a single asterisk; the higher molecular weights (HMWs) corresponding to the newly formed fibrils are detected from 25 kDa to the top of the gel. 20 ng of recombinant monomeric aSyn full-length or C-terminally truncated at residue 114 (1-114) were loaded to assess the size of the truncated PFFs. The graphs represent the mean +/- SD of a minimum of 3 independent experiments. $p<0.05$=*, $p<0.005$=**, $p<0.0005$=*** (ANOVA followed by Tukey HSD *post-hoc* test, PBS vs. PFF-treated neurons), $p<0.005$=##, $p<0.0005$=### (ANOVA followed by Tukey HSD *post-hoc* test, hE83Q PFF-treated neurons vs. hWT or mWT PFF-treated neurons), $p<0.0005$=$$$ (ANOVA followed by Tukey HSD *post-hoc* test, mWT PFF-treated neurons vs. hWT or hE83Q PFF-treated neurons). **(B)** Internalization of PFFs inside the neurons was confirmed by ICC using a total aSyn (epitope: 1-20) antibody. Neurons were counterstained with microtubule-associated protein (MAP2) antibody, and the nucleus was counterstained with DAPI staining. Scale bars = 10 μm. **(C)** Fate and clearance of mWT PFFs after internalization was followed by WB analyses of the insoluble fraction of the PFF-treated aSyn KO neurons up to 21 days (D21). Control neurons were treated with PBS buffer (PBS). Total aSyn and pS129 were detected by SYN-1, pS129 (MJFR13). The total loading level was assessed using the actin antibody. Full-length aSyn (15 kDa) is indicated by a double red asterisk, the C-terminal truncated species (12 kDa) by a single red asterisk. The purple arrows indicate the intermediate aSyn-truncated fragments and the blue arrow the 10 kDa species. **(B-C)** The graphs represent the mean +/- SD of a minimum of 3 independent experiments. $p<0.05$=* (ANOVA followed by Tukey HSD *post-hoc* test, D1 vs. other time-point), $p<0.0005$=### (ANOVA followed by Tukey HSD *post-hoc* test, hE83Q PFF-treated neurons vs. hWT or mWT PFF-treated neurons at D1), $p<0.0005$=£££ (ANOVA followed by Tukey HSD *post-hoc* test, mWT PFF-treated neurons vs. hWT or hE83Q PFF-treated neurons at D1), $p<0.05$=° (ANOVA followed by Tukey HSD *post-hoc* test, mWT PFF-treated neurons vs. hWT or hE83Q PFF-treated neurons at D3), $p<0.05$=$ (D3 vs. D21).

**Figure S20**

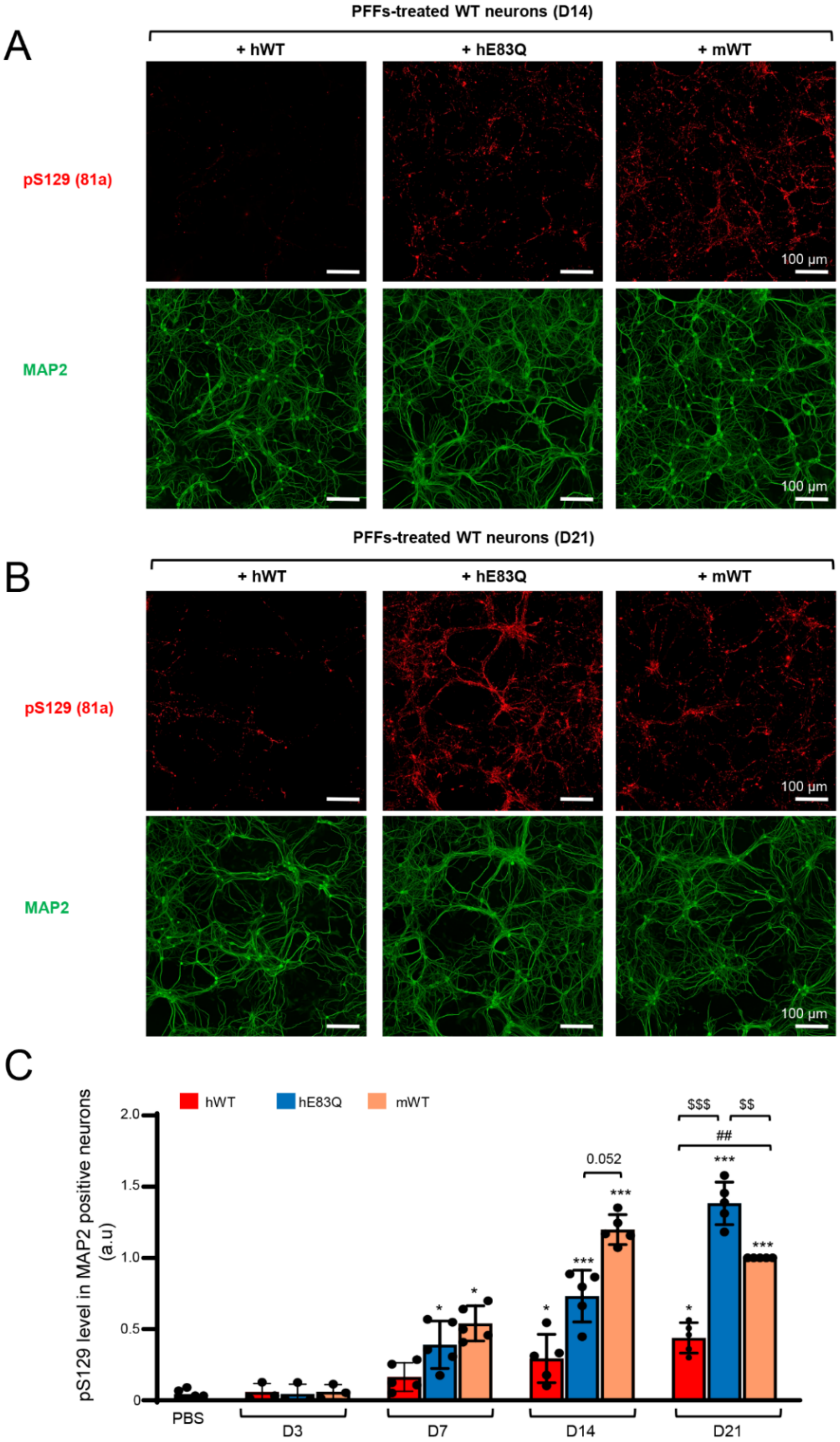
A
PFFs-treated WT neurons (D14)
+ hWT
+ hE83Q
+ mWT
pS129 (81a)
MAP2
100 μm
B
PFFs-treated WT neurons (D21)
+ hWT
+ hE83Q
+ mWT
pS129 (81a)
MAP2
100 μm
C
hWT
hE83Q
mWT
pS129 level in MAP2 positive neurons (a.u)
2.0
1.5
1.0
0.5
0
PBS
D3
D7
D14
D21
0.052
$$$
$$
##
*
***

**Figure S20**

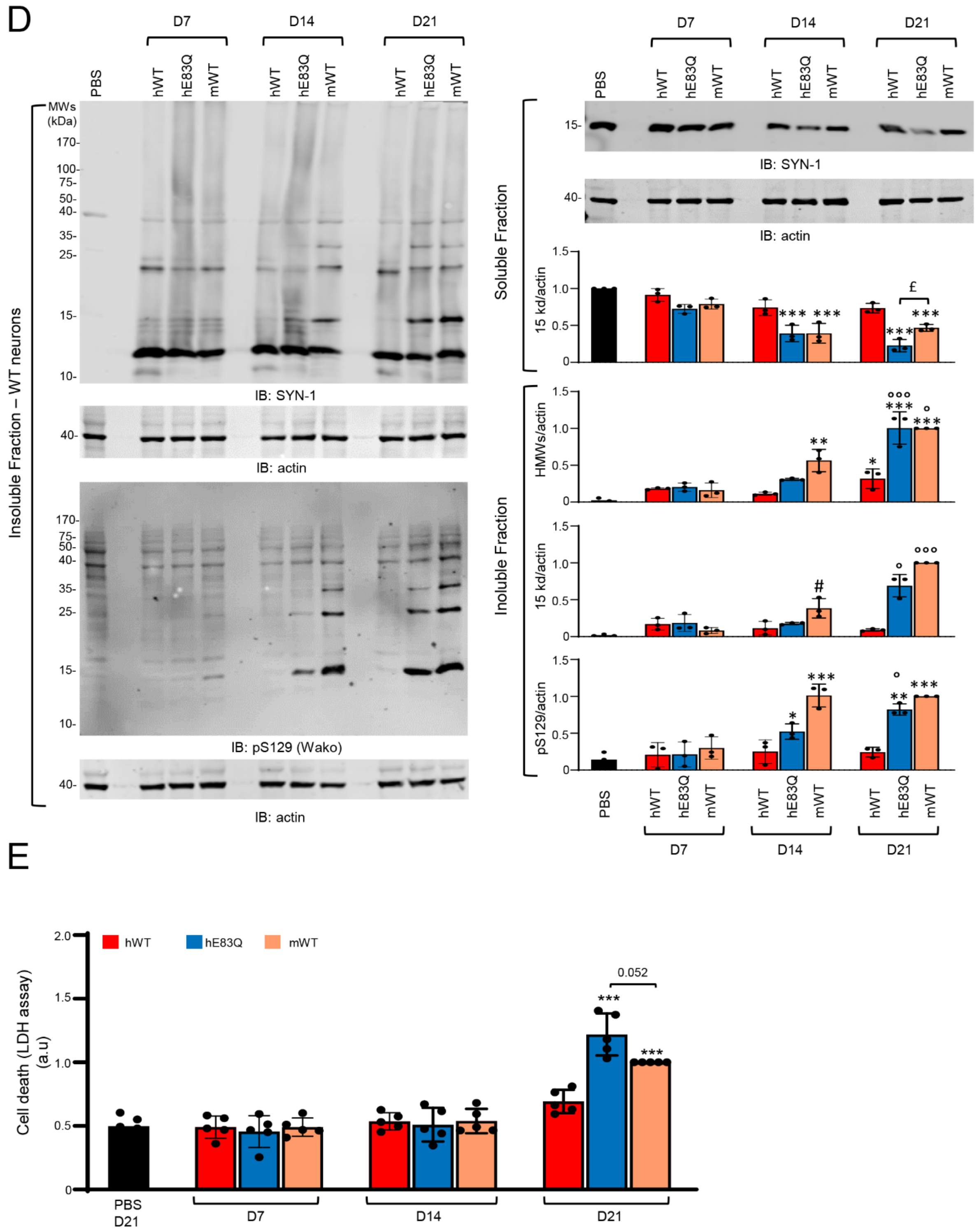


**Fig. S20. E83Q mutation is enough to restore the capacity of human PFFs to seed as efficiently as the mouse PFF in primary neurons.** Temporal analysis of the level of aSyn seeded-aggregates formed in WT neurons after the addition of 70 nM of hWT, hE83Q or mWT aSyn PFFs to the primary culture (at DIV 5) for 7 days (D7), 14 days (D14) or 21 days (D21). Control neurons were treated with PBS. **(A-C)** The level of seeded-aggregates was measured overtime in PBS- or PFF-treated neurons by high content analysis (HCA). Seeded-aggregates were detected by ICC using pS129 (81a) antibody and neurons were counterstained with microtubule-associated protein (MAP2) antibody, and the nucleus was counterstained with DAPI staining. For each independent experiment, a minimum of two wells was acquired per condition, and nine fields of view were imaged for each well. 3 independent experiments were performed. Representative images at D14 (A) and at D21 **(B)**. Scale bars = 100 µm. **(C)** Images were then analyzed using Cell Profiler software to quantify the total level of pS129 in MAP2 positive neurons as previously described [2]. **(D)** The level of seeding in PFF-treated neurons at D7, D14 and D21 was analyzed by WB. After sequential extractions, the soluble and insoluble fractions of neuronal cell lysates were analyzed by immunoblotting. Total aSyn, pS129 and actin were respectively detected by SYN-1, pS129 (Wako or MJFR-13) antibodies. The total loading level was assessed using the actin antibody. In the insoluble fraction, the higher molecular weights (HMWs) corresponding to the newly formed fibrils are detected from 25 kDa to the top of the gel. A minimum of three independent experiments was performed. **(E)** Cell death levels were assessed in WT neurons treated with PFFs (70 nm) at D7, D14 and D21 using lactate dehydrogenase (LDH) release assay. For each independent experiment, triplicate wells were measured per condition. 5 independent experiments were performed. **(C-E)** The graphs represent the mean +/- SD of a minimum of 3 independent experiments. **(C)** $p<0.05$=*, $p<0.005$=**, $p<0.0005$=*** (ANOVA followed by Tukey HSD *post-hoc* test, PBS vs. PFF-treated neurons). $p<0.0005$=### (ANOVA followed by Tukey HSD *post-hoc* test, hWT PFF-treated neurons vs. hE83Q PFF-treated neurons). $p<0.005$=$$, $p<0.0005$=$$$ (ANOVA followed by Tukey HSD *post-hoc* test, mWT PFF-treated neurons vs. hWT or hE83Q PFF-treated neurons). N.S (non-specific, D14 vs D21). **(D)** $p<0.05$=*, $p<0.005$=**, $p<0.0005$=*** (ANOVA followed by Tukey HSD *post-hoc* test, PBS vs. PFF-treated neurons). $p<0.05$=# (ANOVA followed by Tukey HSD *post-hoc* test, D7 vs. D14 mWT PFF-treated neurons). $p<0.05$=£, (ANOVA followed by Tukey HSD *post-hoc* test, mWT PFF-treated neurons vs. hE83Q PFF-treated neurons at D21). **(E)** $p<0.05$=*, $p<0.005$=**, $p<0.0005$=*** (ANOVA followed by Tukey HSD *post-hoc* test, PBS vs. PFF-treated neurons).

**Figure S21**

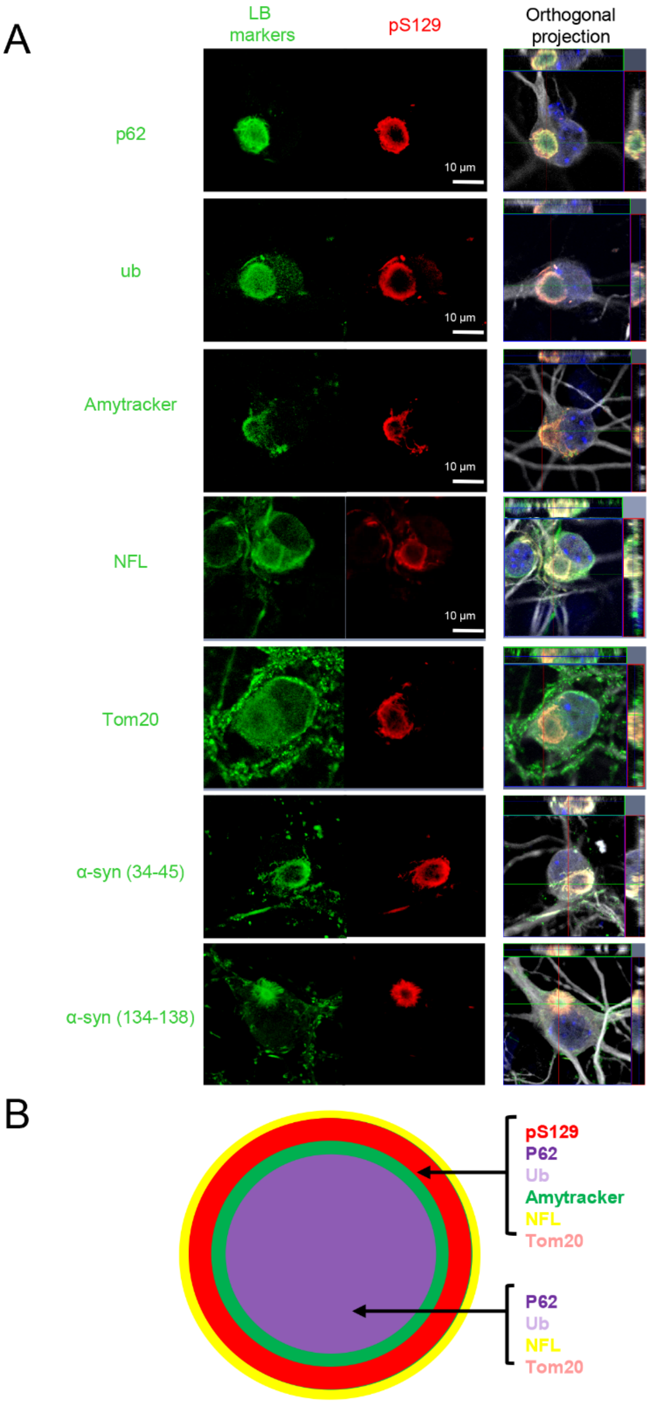


**Fig. S21. E83Q mutation induces the formation of LB-like inclusions that recapitulate the immunohistochemical and morphological features of brainstem LBs observed in PD patient brains. (A)** LB-like inclusions with a ring-like appearance, pS129 aSyn and the Amytracker signal are preferentially localized at the periphery of these inclusions. In contrast, p62 and ubiquitin were detected at both the periphery and the core of the ring-like inclusions as NFL cytoskeletal proteins and mitochondria. Images shown in panel A were from Figs. 9-10 or S12-S15. **(B)** Such organization resembles the highly organized skin-onion-like [3] brainstem LBs inclusions observed in human brain tissues from stage 3–5 PD patients [3, 4, 5, 6, 7, 8, 9].
